# Gene-level alignment of single cell trajectories

**DOI:** 10.1101/2023.03.08.531713

**Authors:** Dinithi Sumanaweera, Chenqu Suo, Ana-Maria Cujba, Daniele Muraro, Emma Dann, Krzysztof Polanski, Alexander S. Steemers, Woochan Lee, Amanda J. Oliver, Jong-Eun Park, Kerstin B. Meyer, Bianca Dumitrascu, Sarah A. Teichmann

**Author notes:** Corresponding authors (S.A.T.). These authors contributed equally to this work (Dinithi Sumanaweera, Chenqu Suo).

## Abstract

Single-cell data analysis can infer dynamic changes in cell populations, for example across time, space or in response to perturbation. To compare these dynamics between two conditions, trajectory alignment via dynamic programming (DP) optimization is frequently used, but is limited by assumptions such as a definite existence of a match. Here we describe **Genes2Genes**, a Bayesian information-theoretic DP framework for aligning single-cell trajectories. **Genes2Genes** overcomes current limitations and is able to capture sequential matches and mismatches between a reference and a query at single gene resolution, highlighting distinct clusters of genes with varying patterns of expression dynamics. Across both real world and simulated datasets, **Genes2Genes** accurately captured different alignment patterns, demonstrated its utility in disease cell state trajectory analysis, and revealed that T cells differentiated *in vitro* matched to an immature *in vivo* state while lacking expression of genes associated with TNFɑ signaling. This use case demonstrates that precise trajectory alignment can pinpoint divergence from the *in vivo* system, thus guiding the optimization of *in vitro* culture conditions.

## Introduction

Recent advances in single-cell genomics, the mainstay of which is single-cell RNA sequencing (scRNA-seq), have revolutionized our understanding of biology and opened up new avenues of research^1^. With their single-cell resolution and ability to observe thousands of genes simultaneously, these new technologies enable the identification of transitional cell states and the study of dynamic cellular processes (e.g. cell differentiation/development; cellular response to perturbations). The computational task of deriving a ‘timeline’ for each dynamic process (e.g. based on transcriptomic similarity) is referred to as ‘pseudotime trajectory inference’^2,3^. One key challenge is how to compare and align two (or more) trajectories, e.g., *in vitro* cell differentiation *versus in vivo* cell development, or responses in control *versus* drug treatment groups (**Fig. 1a**). Identifying genes that differ between *in vitro* and *in vivo* systems, for instance, can guide us to refine *in vitro* cell differentiation.

**Fig. 1.**
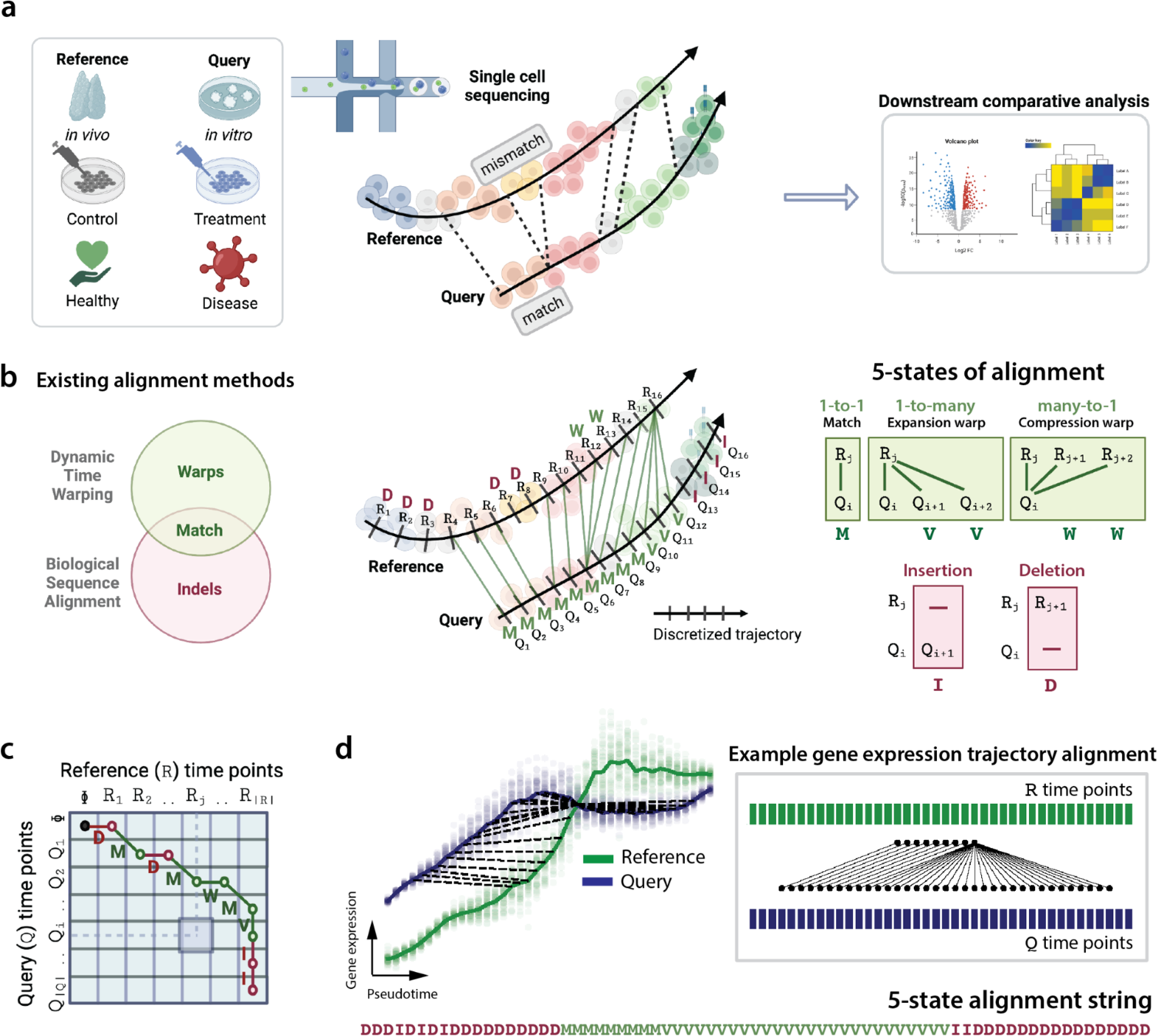
Computational alignment of single-cell transcriptomic trajectories. a, Trajectory alignment is important for comparing different single-cell reference and query systems that dynamically change. This could be between *in vivo* cell development and *in vitro* cell differentiation, or between control and drug-treated cells in response to the same perturbation, or between the responses to vaccination or pathogen challenge in healthy *versus* diseased individuals. A complete alignment between them can capture matches and mismatches for further downstream analysis. **b,** Schematic illustration of the different states of alignment and their theoretical origins. Left: dynamic time warping and biological sequence alignment are complementary to each other^14,15,19^, where both address matches yet capture either warps or indels, respectively. Middle: Illustration of an alignment (a nonlinear mapping) between time points of the discretized reference and query trajectories shown in **a.** Right: between discrete time points in R (reference trajectory) and Q (query trajectory), there may exist 5 different states of alignment: matches (1-1 correspondences), warps (1-to-many expansion or many-to-1 compression correspondences) and mismatches (insertions/deletions denoting a significant difference in one system compared to the other). **c,** Example alignment path across a pairwise time point matrix between R and Q trajectories. Diagonal lines (green) refer to matches (`M`); vertical lines refer to either insertions (`I`) (red) or expansion warps (`V`) (green); horizontal lines refer to deletions (`D`) (red) or compression warps (`W`) (green). Any matrix cell (*i, j*) denotes the pairing of a reference time point R_j_ and query time point Q_i_. **d,** An example gene alignment generated by Genes2Genes. Left: The plot of interpolated log1p normalized expression (*y*-axis) between reference (green) and query (blue) against their pseudotime (*x*-axis). The bold lines represent mean expression trends, while the faded data points are 50 random samples from the estimated expression distribution at each time point. The black dashed lines visualize matches and warps between time points. Right: Schematic illustration of the corresponding nonlinear mapping between R and Q time points shown in the left. Bottom: the corresponding five-state alignment string.

Trajectory comparison poses a time series alignment problem which can be addressed with dynamic programming^4^. The goal is to find an optimal mapping, i.e., a set of pairwise sequential correspondences between the time points of two single-cell trajectories, that captures the matched and mismatched cell states between them. To align two single-cell trajectories, a popular dynamic programming algorithm called dynamic time warping (DTW) is often used. Several studies^5–10^ including the widely-used CellAlign^6^ framework employ DTW to analyze correspondences between time-course profiles, allowing detection of timing differences in biologically similar processes through the notion of warps^11^. Current practice is to interpolate the gene expression time series prior to DTW, and then minimize the Euclidean distance of expression vectors between the matched time points to find an optimal alignment. While DTW is a powerful approach with numerous uses, we are motivated to overcome its main limitations, i.e., (1) the requirement that every time point in each trajectory needs to match with at least one time point in the other trajectory; (2) the inability to identify mismatches (missing data or substantial differences between two series), occurring in the form of insertions or deletions (indels); and (3) the use of a distance metric which relies only on the difference of mean expression, without considering their distributions.

Warps and indels are fundamentally distinct (**Fig. 1b-c**). This is particularly highlighted in discussions about integrating DTW with the concept of gaps^12,13^ (as in biological sequence alignment^14,15^). Both matches and mismatches between trajectories inform our understanding of temporal gene expression dynamics. A mismatch could either imply missing data or differential expression (DE). For instance, a sudden rise or drop in expression of one system relative to the other might indicate that it is transitioning through a different cell state. A mismatch can also occur when a considerable fraction of cells in one condition has a significantly different distribution of expression for some genes compared to the other condition. On the other hand, matches imply similar cell states, and warps indicate differences in their relative speeds of cell transition in diverse contexts. Matches and mismatches together can also inform specific patterns such as divergence and convergence (e.g. **Fig. 1d**), allowing their computational separation for downstream analysis. Moreover, an alignment is a means to properly identify DE genes between trajectories. For instance, Laidlaw et al. (2023)^16^ showed that trajectory alignment successfully captures DE genes undetectable by non-alignment methods^17,18^

Here we present Genes2Genes (G2G; **Fig. 2, Methods**), a novel framework for aligning single-cell transcriptomic trajectories of a reference and query system at single-gene resolution. G2G utilizes a DP alignment algorithm that accounts for matches, warps and indels by combining the classical Gotoh’s algorithm^15^ and DTW^19^, inspired by the concepts of biological sequence alignment discussed in the related literature^20–22^. Its scoring scheme employs a new Bayesian information-theoretic measure based on minimum message length inference^23–25^ to quantify the distance between two time points based on their gene expression distributions. G2G (1) generates descriptive pairwise alignments at gene-level, (2) identifies gene clusters of similar alignment patterns, (3) derives an aggregate (average) cell-level alignment across all or subset of genes, (4) identifies genes with differential dynamic expression profiles, and (5) explores their associated biological pathways. As it formally handles matches and mismatches in a principled way, G2G overcomes the need to impose ad hoc thresholds^6^ and/or post hoc processing of a usual DTW output (as in TrAGEDy^16^, the recent advancement built on CellAlign^6^) for capturing differential regions in gene expression.

**Fig. 2.**
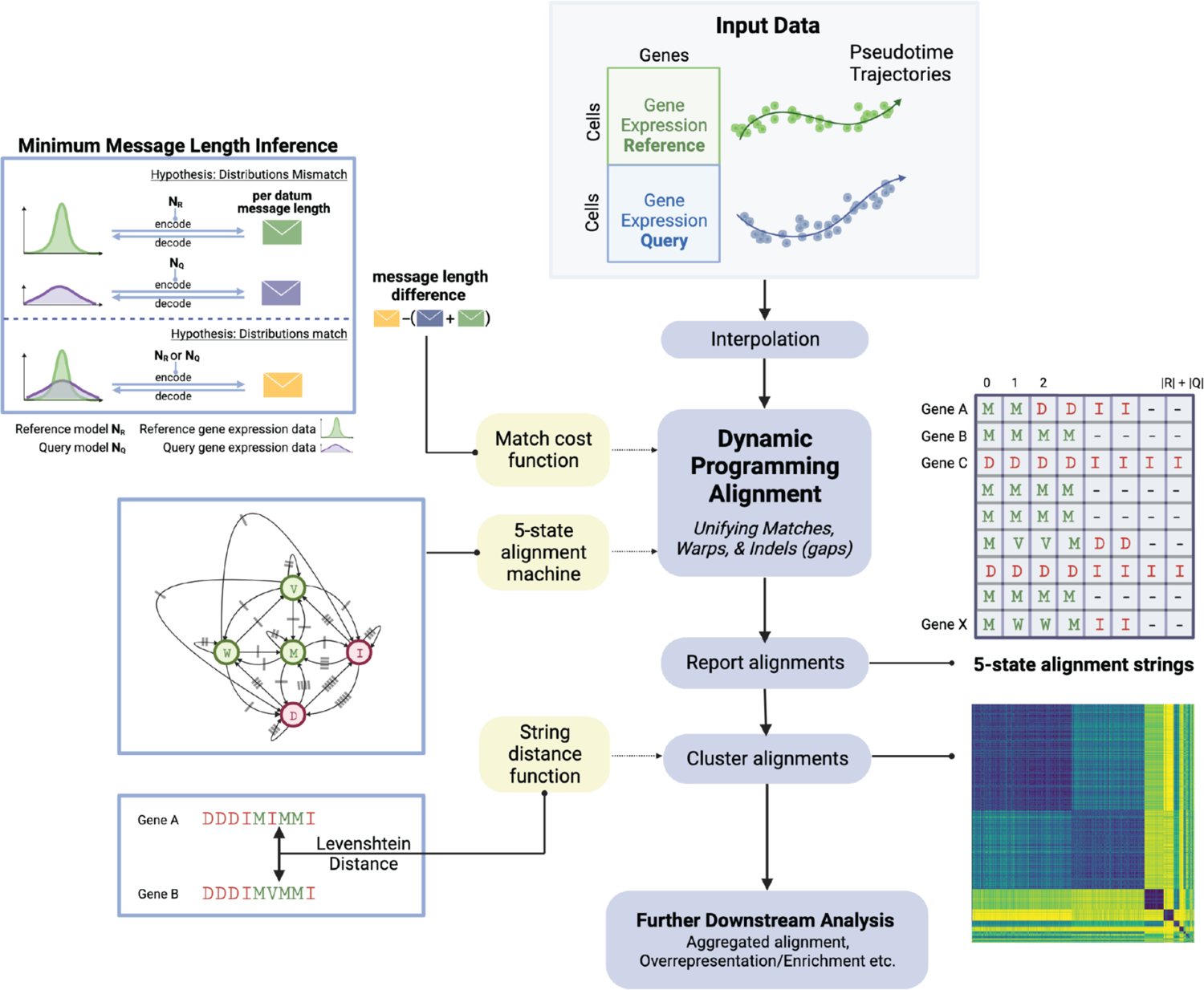
Overview of the Genes2Genes (G2G) alignment framework for comparing single-cell transcriptomic trajectories. Schematic illustration of G2G workflow. Given log1p normalized cell-by-gene expression matrices of a reference (R) and query (Q), and their pseudotime estimates, G2G infers individual alignments for a list of genes of interest. It first interpolates data by extending mean based interpolation in Alpert et al (2018)^6^ to distributional interpolation, and then runs Gotoh’s dynamic programming (DP) algorithm^15^ adapted for a five-state alignment machine, defining a match (`M`), compression warp (`W`), expansion warp (`V`), insertion (`I`) and deletion (`D`). All reported alignments are then clustered and used to deliver statistics on the overall alignment between R and Q, supporting further downstream analyses. **Top left,** the DP algorithm utilizes a match cost function defined under minimum message length (MML)^24^ statistical inductive inference. Given a hypothesis (a distribution model) and data, MML can define the total message length of encoding them for lossless compression along an imaginary message transmission. G2G defines two hypotheses: (1) Φ: *R*_j_ and *Q*_i_ time points mismatch, and (2) A: *R*_j_ and *Q*_i_ time points match. Under Φ, the message length is the sum of independent encoding lengths of their corresponding interpolated expression data and distributions. Under A, the message length is their joint encoding length of the corresponding interpolated expression data with a single Gaussian distribution (either of *R*_j_ or *Q*_i_). The match cost is taken as the difference of A and Φ per-datum encoding lengths. **Middle left,** The DP algorithm incorporates a (five-state) finite state machine which can generate a string over the alphabet, Ω = [`M`, `W`, `V`, `D`, `I`] that describes all the 5 possible states of alignment (**Fig. 1b**) between two time points. Each arrow represents a state transition. Arrows with the same hatch mark implies equal probability of state transition. **Bottom left,** G2G computes a pairwise Levenshtein distance matrix across all 5-state alignment strings to cluster genes of similar alignment pattern. **Top right,** example output of five-state alignment strings for each gene. **Bottom right,** example clustermap showcasing the clustering structure of alignment strings resulted from Agglomerative hierarchical clustering. The color represents the Levenshtein distance.

We first validate G2G’s ability to accurately capture matches, mismatches, and different alignment patterns in simulated datasets, benchmarking against CellAlign^6^ and TrAGEDy^16^ (the current state-of-the-art of single-cell trajectory alignment). We then demonstrate how G2G offers gene-level alignment between two stimulated groups of cells in a published real dataset^26^. We further explore how our framework can facilitate a healthy versus disease comparison, using a published idiopathic pulmonary fibrosis (IPF) dataset^27^. We finally show how G2G quantitatively aligns *in vitro* and *in vivo* T cell development. We find that the TNFα signaling pathway in the final stage of *in vivo* T cell maturation is not recapitulated *in vitro*, and validate G2G’s use in identifying potential targets for optimizing *in vitro* cell engineering.

## Results

### Genes2Genes (G2G) aligns single-cell trajectories with dynamic programming, employing a Bayesian information-theoretic measure

G2G is a new dynamic programming (DP) framework to infer and analyze gene trajectory alignments between a single-cell reference and query. Given a reference sequence *R* (a discrete series of time points: *R*_1_, *R*_2_,… *R*_j_,… *R*_|*R*|_) and query sequence *Q* (a discrete series of time points: *Q*_1_, *Q*_2_,… *Q*_j_,… *Q*_|&|_), a computational alignment between them can inform us of the one-to-one matches, one-to-many matches (expansion warps), many-to-one matches (compression warps), insertions, and deletions (indels) between their time points in sequential order, denoted by `M,V,W,I,D`, respectively (**Fig. 1b**). While matches and warps imply similarity between transcriptomic states of *R* and *Q*, indels (a.k.a gaps) imply mismatches (differential or missing transcriptomic states compared to each other). A standard DP alignment algorithm finds an optimal alignment between two sequences by constructing a pairwise cost matrix and generating the path which minimizes the cost (**Fig. 1c**). This relies on a scoring scheme to quantify correspondences between every pair of *R* and *Q* time points.

Unlike the current DP alignment approaches (i.e. DTW and DNA/protein sequence alignment), G2G implements a DP algorithm that handles both matches (including warps) and mismatches jointly, and it runs between the reference and query for each gene. This algorithm extends Gotoh’s three-state alignment algorithm^15^ to five-states (defined in **Fig. 1b**) for accommodating warps, allowing a nonlinear mapping between the pseudotime axes of *R* and *Q*. In the sequence alignment domain, the standard Gotoh’s algorithm defines time-efficient DP recurrences over `M,I,D` states to align sequences under an affine gap penalty scheme (i.e. different penalties for gap open and extension), which is easily extendable to define more alignment states and probabilistic state transitions between these states^20–22^. **Fig. 1d** illustrates an example of gene alignment generated by G2G, described as a 5-state string. Reading the 5-state string left-to-right defines the matches, warps, and mismatches of *R* and *Q* time points in sequential order, similar to the format reported for DNA/protein pairwise alignment.

Our DP scoring scheme incorporates a Bayesian information-theoretic cost function based on minimum message length (MML) inference^23–25^ (top left of **Fig. 2, Supplementary** Fig. 1), and the state transition probabilities from a five-state machine (middle left of **Fig. 2**). The MML criterion allows us to compute a symmetric cost (named MML distance) for matching any reference time point *R*_j_ and query time point *Q*_i_ based on their corresponding gene expression distributions. We evaluate their distributional difference in terms of both mean and variance, acknowledging that one trajectory may be noisier than the other. The five-state machine allows to compute a symmetric cost of assigning an alignment state for *R*_j_ and *Q*_i_ out of the five possible states. Each cost term is computed as the Shannon information^28^ measured in ‘nits’ based on the probability models defined for the corresponding events, i.e., Shannon information of event *E* with probability *Pr*(*E*) is: −*log*(*Pr*(*E*)) nits. See **Methods** for details.

### Overview of the G2G framework

The G2G framework is composed of several components to support single-cell trajectory comparison, which include input preprocessing, DP alignment algorithm, alignment clustering and downstream analysis (**Fig. 2**). G2G’s inputs are log1p normalized scRNA-seq matrices of some reference and query, and their pseudotime estimates. G2G then performs interpolation to smooth each gene expression trajectory. This first transforms the pseudotime axis to [0,1] range using min-max normalization, over which we take a predefined number of equispaced interpolation time points, similar to CellAlign^6^. For each interpolation time point, we estimate a Gaussian distribution of gene expression, taking into account all cells, kernel-weighted^6^ by their pseudotime distance to this interpolation time point. This way, our approach fits the entire distribution instead of estimating only the mean expression at each interpolation point. The interpolated gene trajectories of the reference and query are then aligned using our DP algorithm. This generates optimal trajectory alignments for all input genes, described as five-state strings (**Fig. 1d** and top right matrix of **Fig. 2**). The 5-state string of a gene informs the percentage of match calling (including one-to-one matches and warps), which we term as ‘alignment similarity’. These strings are then used to compute a pairwise distance matrix between genes (using the Levenshtein distance). The genes displaying similar alignment patterns are then clustered together using agglomerative hierarchical clustering. Alignments and their cluster memberships allow further downstream analysis such as gene set overrepresentation analysis. Clusters reveal the diversity of alignment patterns across genes (e.g. 100% mismatched, 100% matched, 20% early-matched and late-mismatched etc.). G2G also generates a representative alignment for each cluster by aggregating all its gene-level alignments (e.g. cluster of 100% mismatches, represented by a string over `I,D`; cluster of 100% matches, represented by a string over `M,V,W`). Finally, G2G aggregates all gene-level alignments into a single, cell-level alignment, informing an average mapping between the trajectories. Both gene-level and cell-level alignments are particularly useful when there is heterogeneity in alignment patterns among different genes.

### G2G expands the capacity of DTW

G2G infers statistically-consistent, sequential matches and mismatches between reference and query time points. Such output is impossible from DTW (e.g. CellAlign^6^) due to mapping all time points regardless of their transcriptomic differences (**Fig. 3a**). In contrast, G2G refrains from mapping time points with dissimilar expression. One could perform local DTW with user-defined thresholds^6^ or post-hoc processing of DTW alignment (as in TrAGEDy^16^) to unmap dissimilar time points, yet the underlying assumption of a definite match still remains. This is particularly broken in datasets with no shared process^16^. With G2G, a user no longer needs ad hoc thresholding nor post-processing, as our algorithm systematically disconnects time points of differential expression. Altogether, the algorithmic novelties and outputs fundamentally distinguish G2G from the existing approaches. (See **Fig. 3a-b** and **Supplementary Table 1** for summarized comparison between CellAlign^6^, TrAGEDy^16^, and G2G).

**Fig. 3.**
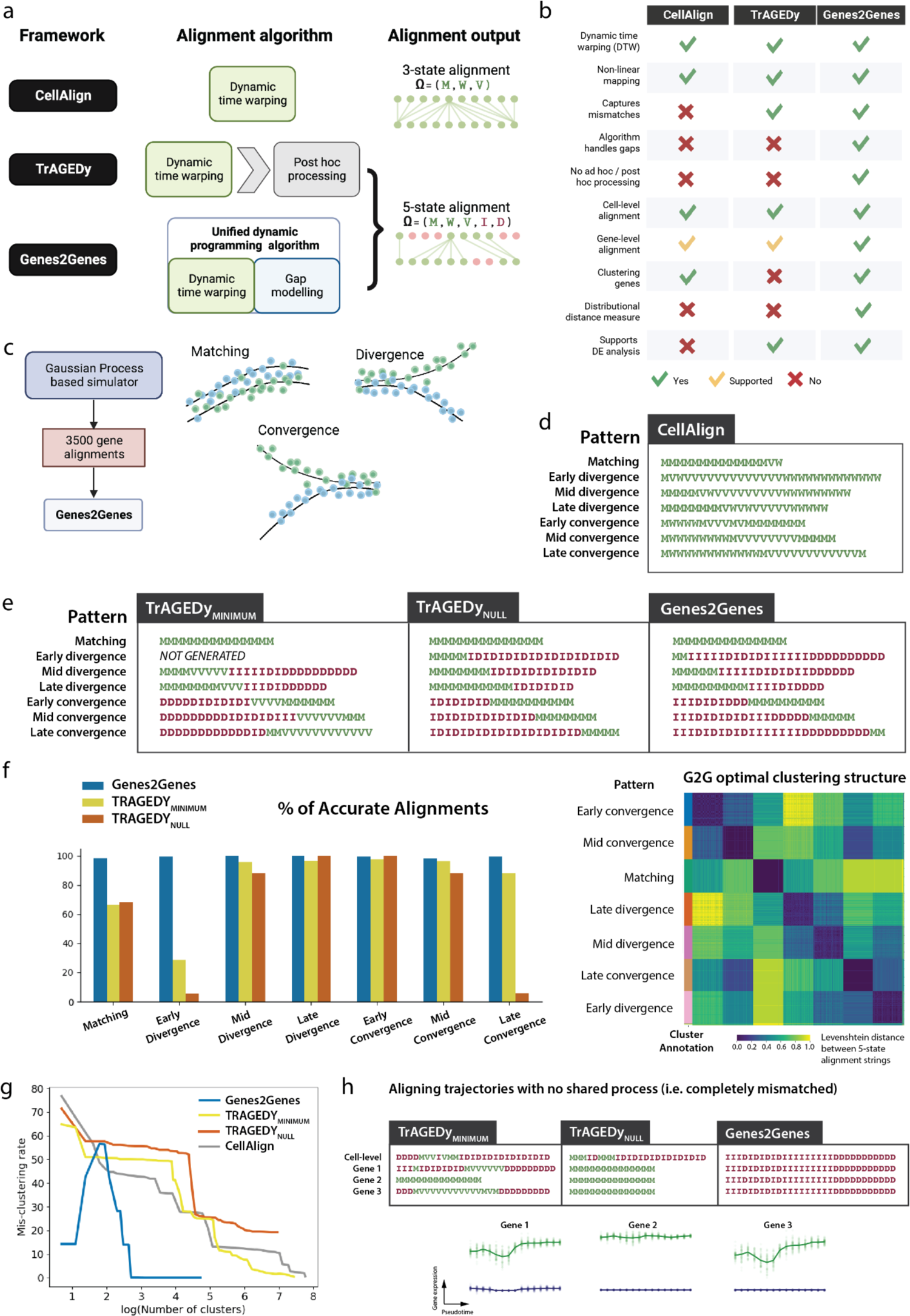
Genes2Genes (G2G) outperforms the current state-of-the-art of trajectory alignment. a, Illustration of the differences in the algorithms and alignment outputs of CellAlign, TrAGEDy, and G2G. CellAlign runs dynamic time warping (DTW), thus generating an alignment only defining the three match states: Ω = [`M`, `W`, `V`] (described in **Fig. 1b**). TrAGEDy performs post-hoc processing of DTW output to disconnect dissimilar time points. G2G runs a new dynamic programming algorithm that unifies DTW and gap modeling. Both TrAGEDy and G2G mappings define all five states of alignment: Ω = [`M`, `W`, `V,I,D`] (described in **Fig. 1b**). **b,** Tabular comparison of features across CellAlign, TrAGEDy, and G2G. **c,** Experiment 1 uses a Gaussian Process based simulator to generate 3500 simulated pairs of reference and query gene trajectories for testing G2G and benchmarking against CellAlign and TrAGEDy. This dataset covers three main classes of alignment pattern: (1) *Matching*, (2) *Divergence* and (3) *Convergence*. The *Divergence* and *Convergence* groups are further sub-categorized based on their approximate location of bifurcation (0.25, 0.5, and 0.75 within [0,1] pseudotime range), resulting in seven distinct classes of pattern (each with 500 alignments). **d,** The three-state, cell-level alignment generated by CellAlign for each pattern dataset (under 15 equispaced time points). **e,** The five-state, cell-level alignments generated by both modes of post-hoc processing implemented by TrAGEDy (referred to as TrAGEDy_MINIMUM_ and TrAGEDy_NULL_), and G2G. **f,** Left: Comparing TrAGEDy and G2G by their percentages of accurate alignments that capture the expected patterns across all the seven classes. Right: The clustergram of the pairwise Levenshtein distance matrix across all G2G alignments, which clearly shows the distinct classes of pattern as separated by agglomerative hierarchical clustering. **g,** The plot compares hierarchical clusterings of the alignment strings generated by CellAlign, TrAGEDy, and G2G, by varying the clustering resolution (from the smallest number of clusters to the maximum number of clusters). The x-axis is the number of clusters in log scale, and the y-axis is the mis-clustering rate (the percentage of the total number of outliers across all clusters). **h,** The cell-level alignment of two simulated trajectories with no shared process, and the gene-level alignment of three example genes, generated by TrAGEDy and G2G, where the expected alignment pattern is a complete 100% mismatch.

### G2G accurately identifies different patterns of alignment with simulated data experiments

To benchmark how well G2G performs alignment compared to CellAlign^6^ and TrAGEDy^16^, we tested on three simulated datasets: (1) an artificial dataset with seven alignment patterns, (2) a real dataset with artificial perturbations, and (3) a negative control dataset. All CellAlign^6^ and TrAGEDy^16^ alignments were converted into 5-state strings before comparison. TrAGEDy^16^ prunes DTW matches based on the minimum dissimilarity score (hereafter referred to as TrAGEDy_MINIMUM_), with an alternative option to disregard the minimum (hereafter referred to as TrAGEDy_NULL_); We tested both modes.

### Experiment 1

We simulated 3500 trajectory pairs under three main classes of pattern: (1) *Matching* (500 genes), (2) *Divergence* (1500 genes), and (3) *Convergence* (1500 genes), using Gaussian Processes and suitable kernels^29,30^ (**Fig. 3c**, Extended Data Fig. 1a-c**, Methods**). Each trajectory comprises 300 cells spread across pseudotime range [0,1]. Each *Divergence* and *Convergence* group has an equal number of pairs, bifurcating at three time points (approximately at time point *t*_*b*_ ∈ [0.25, 0.5, 0.75], indicating early, mid, and late bifurcation, respectively). We first examined G2G performance in identifying matched and mismatched regions accurately for each pattern (under 15 optimal time points in [0,1] pseudotime range). As illustrated in **Fig. 1b-d**, a subsequence over `M,W,V` denotes a matched region, while a subsequence over `I,D` denotes a mismatched region. We expect the correct alignment to be 100% match for *Matching* trajectories; a matched region (start-match) followed by a mismatched region (end-mismatch) for *Divergence*; and a mismatched region (start-mismatch) followed by a matched region (end-match) for *Convergence*. The match/mismatch lengths for *Divergence*/*Convergence* depend on their bifurcation locations (**Extended Data** Fig. 1d-e). Thus we also report the distributions of start-match lengths (following a false mismatch if there is any), end-mismatch lengths, and start-mismatch (false mismatch) lengths in all alignments under each bifurcation group, to see if they fall within the expected ranges. We used the accuracy rate (the proportion of correct gene alignments) to fine-tune the optimal parameter setting (i.e. state transition probabilities) of the 5-state machine used by our DP algorithm, which was set as the default for G2G (See **Methods**, **Supplementary Table 2**).

**Fig. 3d-e** reports cell-level alignments generated by all methods in comparison. The G2G aggregate alignments for the 7 groups (**Fig. 3e**, right-most) accurately reflect their distinct matching and mismatching patterns. In contrast, CellAlign^6^ cannot produce expected patterns for *Divergence* and *Convergence* (**Fig. 3d**), as it only covers the match states shown in **Fig. 1b**. TrAGEDy outputs expected patterns after post-hoc processing DTW alignments, with higher match lengths (**Fig. 3e**, left).

Across all the seven distinct patterns, G2G outperforms both modes of TrAGEDy with high accuracy in generating expected gene-level alignments (**Fig. 3f** left). The G2G accuracy rates are: 98.6%, 99.4%, 99.8%, 100%, 99.2%, 98.2%, 99.2% for *Matching, Divergence* (early, mid, late), and *Convergence* (early, mid, late) pairs, respectively. All distributions of start-match (start-mismatch) lengths and end-mismatch (end-match) lengths in *Divergence*/*Convergence* alignments fall within the expected ranges (**Extended Data** Fig. 2a-b). In contrast, TrAGEDy_MINIMUM_ gives 66.26%, 28.57%, 95.87%, 96.86%, 97.35, 96.15, 88.2% accuracy rates, respectively, with *Divergence*/*Convergence* alignments showing higher variability in their match and mismatch lengths compared to G2G, thus falling out of the expected length ranges. TrAGEDy_NULL_ gives 68.2%, 5.4%, 88%, 100%, 100%, 88%, 5.8% accuracy rates, respectively, with better length distributions than TrAGEDy_MINIMUM_. G2G shows fewer false mismatches on average for *Matching* alignments compared to TrAGEDy, while also having fewer intermediate false mismatches compared to TrAGEDy_MINIMUM_. Interestingly, TrAGEDy_NULL_ generates no intermediate false mismatches, yet results in higher inaccuracy due to 100% matched or expected-order-swapped alignments.

G2G clustering separates the seven distinct patterns very well (**Fig. 3f** right), i.e., hierarchical agglomerative clustering of alignments at the optimally-chosen 0.22 Levenshtein distance threshold gives 15 clusters, with only 0.1% mis-clustering rate. (**Extended Data** Fig. 2c, See **Methods** for details on choosing the optimal threshold). We first compared this to CellAlign’s^6^ k-means clustering of genes based on their pseudotime shifts (i.e. pseudotime differences between matched time points in gene-level DTW alignments) from k=7 to k=50. All mis-clustering rates were significantly higher [42.6%, 60.4%] (**Extended Data** Fig. 2d) than G2G’s low mis-clustering rate. For the hierarchical clustering of CellAlign^6^ and TrAGEDy alignment strings, we observe higher noise and mis-clustering rates, compared to G2G (**Fig. 3g**).

### Experiment 2

To test how G2G detects matching patterns in real scRNA-seq data, we used a dataset of E15.5 murine pancreatic development^31^ and considered 769 genes varying in expression during beta-cell differentiation. We randomly split cells into query and reference, and simulated a mismatch in the form of a deleted portion (perturbation scenario 1) or changed portion (perturbation scenario 2) of increasing size at the beginning of the trajectory (**Extended Data** Fig. 3a). We then performed gene-level alignments using G2G and TrAGEDy (with 50 interpolation time points) for all different cases under each scenario, and calculated their alignment similarity percentages (**Extended Data** Fig. 3b-c). For perturbation scenario 1, both G2G and TrAGEDy alignment similarity decreases with increasing deletion sizes as expected across smaller perturbation sizes, although the detected mismatch length is shorter than expected for deletions larger than 20%. This could be due to the relatively non-varying gene expression between pseudotime bin 10 to 20 (**Extended Data** Fig. 3d), which causes warps instead of mismatches. For perturbation scenario 2, the alignment similarity has an expected maximum and minimum (**Extended Data** Fig. 3e).

Accordingly, the observed trends for both methods generally follow the expected trends, falling within the expected ranges for larger perturbation sizes. Interestingly, TrAGEDy_NULL_ performs better than TrAGEDy_MINIMUM_ in both perturbation scenarios. TrAGEDy_NULL_ also shows higher accuracy for perturbation sizes < 6%. Overall, both G2G and TrAGEDy_NULL_ show close performance with better match accuracy detection than TrAGEDy_MINIMUM_. However, G2G shows relatively less variability in results across all cases.

### Experiment 3

We also examined two simulated datasets with no shared process (referred to as negative control and tested by TrAGEDy^16^). G2G generated an aggregate alignment with solely insertions and deletions (i.e. 100% mismatch) as expected, while both modes of TrAGEDy^16^ alignment falsely included segments of matches (**Fig. 3h**). Similar results were seen for three individual genes with completely mismatched trajectories. This is an example where the DTW assumption breaks, highlighting the benefit of G2G.

In conclusion, G2G outperforms existing methods by accurately aligning and clustering genes with different alignment patterns, while advancing the existing framework for trajectory alignment.

### G2G captures mismatches and offers gene-level resolution alignment

To further demonstrate features of our framework, we performed G2G alignment on the same dataset^26^ tested by CellAlign^6^ (**Fig. 4a**). This is a time-course dataset of murine bone marrow-derived dendritic cells treated with PAM3CSK (PAM) or lipopolysaccharide (LPS) to simulate responses to different pathogens.

**Fig. 4.**
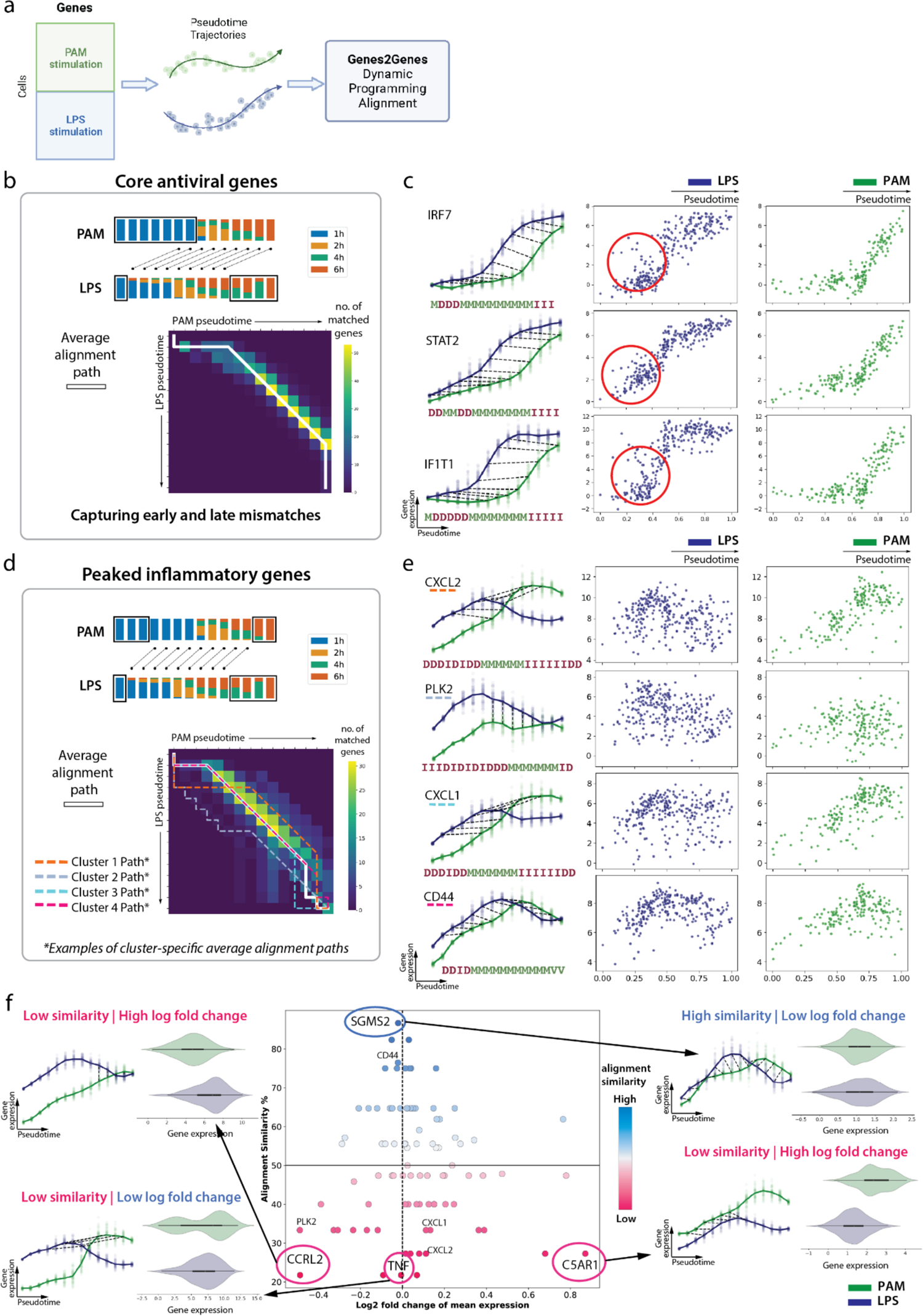
G2G captures mismatches and offers gene-level resolution alignment. a, G2G alignment was performed on a published time-course dataset^26^ of murine bone marrow-derived dendritic cells stimulated with PAM or LPS. Both the gene expression data and the inferred pseudotime were taken from Alpert et al. 2018^6^. **b,** Top: schematic illustration of the aggregate alignment result for all the genes in the “core antiviral module”. The stacked bar plots represent the cell compositions at each time point (14 equispaced time points on pseudotime [0,1]), colored by the time of sampling post stimulation. Boxed segments represent mismatched time points. The black dashed lines represent matches and warps between time points. Bottom: pairwise time point matrix between PAM and LPS pseudotime. The color represents the number of genes showing match or warp for the given pair of a PAM time point and an LPS time point. The white line represents the main average alignment path. **c,** Gene expression plots for three representative genes (*IRF7, STAT2* and *IFIT1*) from the “core antiviral module” in LPS-stimulated (blue) and PAM-stimulated (green) data along their pseudotime. Left column: the interpolated log1p normalized expression (*y*-axis) against pseudotime (*x*-axis). The bold lines represent mean expression trends, while the faded data points are 50 random samples from the estimated expression distribution at each time point. The black dashed lines represent matches and warps between time points. Right two columns: the actual log1p normalized expression (*y*-axis) against pseudotime (*x*-axis). Each point represents a cell. The five-state alignment string for each gene is shown below the expression plots. Red circles highlight the cells with high expression values at early time points, which are referred to as ‘precocious expressers’. **d,** The same plots as in **b** for genes in the “peaked inflammatory module”. In the pairwise time point matrix, the white line represents the main average alignment path. The genes are also clustered based on their alignment results (**Extended Data** Fig. 2), and dashed lines with different colors represent examples of cluster-specific alignment paths. **e,** The same plots as in **c** for four representative genes (*CXCL2, PLK2, CXCL1, CD44*) from each cluster shown in **d**. **f,** Plot of alignment similarity (*y*-axis) against log_2_ fold change of mean expression (*x*-axis) for all genes in the “peaked inflammatory module” (middle). The color also represents the alignment similarity. The surrounding plots show the interpolated log1p normalized expression (*y*-axis) against pseudotime (*x*-axis) on the left, and the violin plot of total gene expression on the right for four selected genes (*SGMS2, CCRL2, TNF, C5AR1*).

G2G’s ability to capture mismatches is revealed when aligning genes from the “core antiviral module” of this dataset (**Extended Data** Fig. 4a). CellAlign^6^ demonstrated a “lag” in gene expression after PAM stimulation compared to LPS^6^. This is also captured by G2G aggregate alignment whereby later PAM pseudotime points were mapped to earlier LPS pseudotime points (**Fig. 4b**). In addition, G2G identified mismatches in the early and late pseudotime points. Clustering alignments did not reveal much diversity, implying that all genes generally follow the average pattern (**Extended Data** Fig. 4b). At early pseudotime points, the gene expression was consistently low in the PAM condition, whereas some LPS-stimulated cells were already showing elevated expression (e.g. *IRF7, STAT2, IF1T1* in **Fig. 4c**). These have also been noticed and described as “precocious expressers” in the original paper^26^. The mismatch in late LPS pseudotime points was caused by the peaked expression, while the expression of PAM-stimulated cells was still on the rise, not reaching a peak yet.

For genes in the “peaked inflammatory module”, **Fig. 4d** shows their G2G aggregate alignment. Clustering of genes revealed cluster-specific average alignments different from the main alignment (**Fig. 4d**, Extended Data Fig. 4c-e). Representative genes from different clusters (shown in **Fig. 4e**) display subtle differences in the length and position of the time points being matched. Using G2G alignment similarity statistics (**Fig. 4f**), we also identified *SGMS2* as the most similar gene (with low log fold change), and *CCRL2* and *C5AR1* as highly dissimilar genes (with high log fold change) between PAM- and LPS-stimulated trajectories. *CCRL2* alignment shows a late convergence, as it was expressed highly at the beginning and peaked early in the LPS-stimulated cells, whereas the expression grew as a slow incline following PAM stimulation. We also note *TNF*, which is highly dissimilar (low alignment similarity) despite its almost negligible log fold change. This difference would be undetectable by a standard DE test (e.g. Wilcoxon rank-sum P-value=0.2), hence highlighting the importance of trajectory alignment. The above results again showcase how G2G captures mismatched regions between scRNA-seq trajectories.

### G2G captures early and late differences in disease state epithelial cell differentiation

To highlight G2G algorithm’s versatility in single-cell studies, we explored its ability to discern disease-related genes from those in healthy states. We compared two cell differentiation trajectories from healthy lung versus diseased lung in idiopathic pulmonary fibrosis (IPF). IPF is an incurable and irreversible disease characterized by deposition of extracellular matrix by myofibroblasts, scarring and progressive loss of lung function, with an estimated survival rate of 3-5 years after diagnosis^32^. Using the Adams et al. (2020) dataset^27^, we investigated the differentiation of alveolar type 2 (AT2) cells into alveolar type 1 (AT1) cells in the healthy lung versus AT2 differentiation into aberrant basaloid cells (ABC) in the IPF lung (**Fig. 5a**), as suggested by several studies^33,34^. ABCs have only recently been described in single-cell studies from IPF patients^33,35–37^, and their origin and role in IPF pathogenesis is still not understood.

**Fig. 5.**
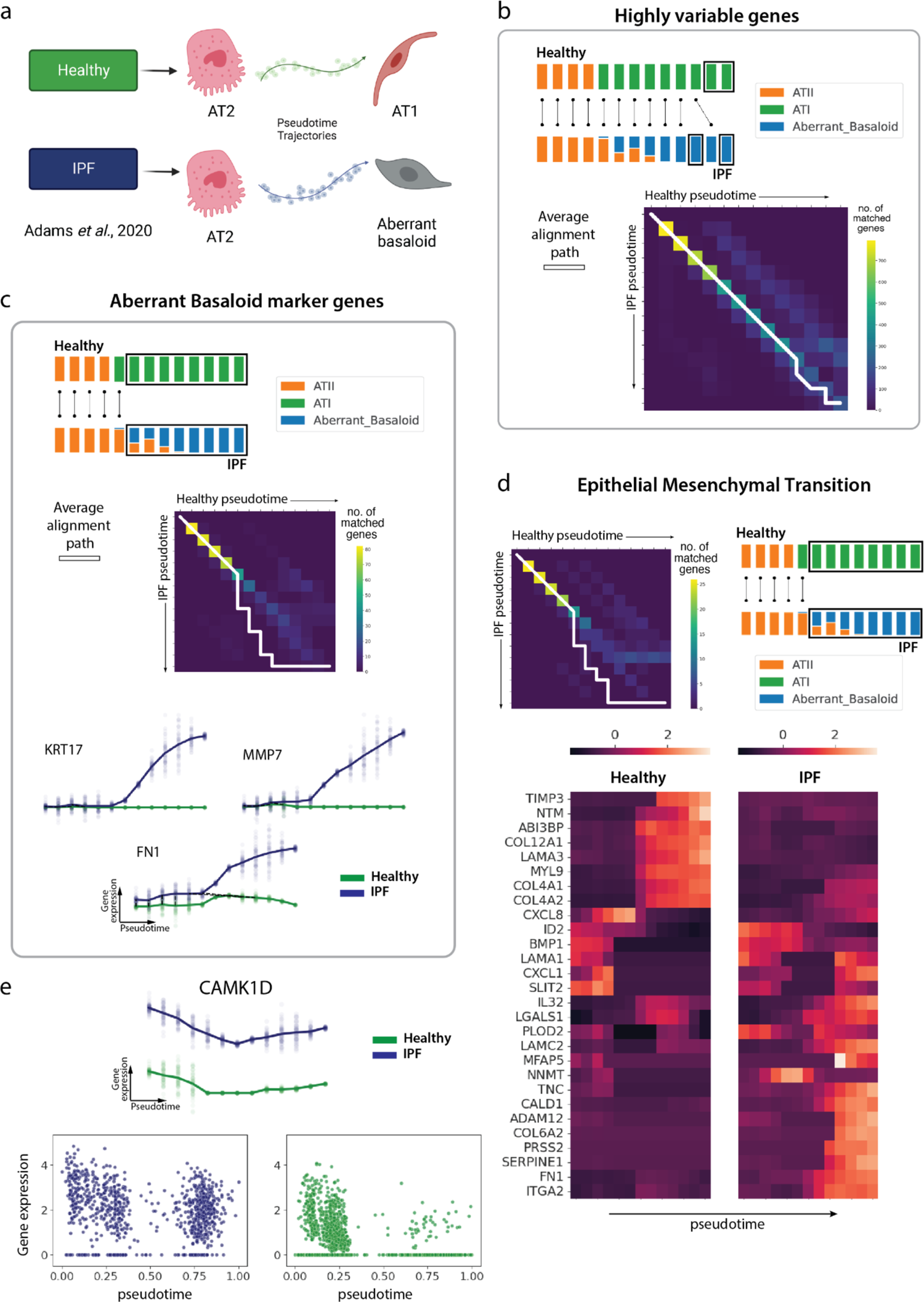
Healthy versus disease case study: aligning differentiation of alveolar type 2 (AT2) cells into alveolar type 1 (AT1) cells in the healthy lung versus aberrant basaloid cells (ABC) in the IPF lung with G2G. a, Schematic illustration of the healthy and IPF cell differentiation trajectories of focus. b, Top: schematic illustration of the aggregate alignment result for all the highly variable genes (HVGs). The stacked bar plots represent the cell type composition at each time point (13 equispaced time points on pseudotime [0,1]). Boxed segments represent mismatched time points. The black lines represent matched time points. Bottom: The pairwise time point matrix between healthy and IPF pseudotime. The color represents the number of genes showing match or warp for the given pair of healthy and IPF pseudotime points. The white line represents the main aggregate alignment path. c, The aggregate alignment result for 88 ABC marker genes (Supplementary Fig. 3) plotted as in b, with the schematic illustration on top, and the pairwise time point matrix in the middle. Bottom: Gene expression plots for three example ABC marker genes (*KRT17, MMP7, FN1*) between IPF (blue) and healthy (green) data along their pseudotime. These plot the interpolated log1p normalized expression (*y*-axis) against pseudotime (*x*-axis). The bold lines represent mean expression trends, while the faded data points are 50 random samples from the estimated expression distribution at each time point. The black dashed lines represent matches and warps between time points. d, Top: (Left) The aggregate alignment path for all Epithelial Mesenchymal Transition (EMT) pathway genes, plotted on the pairwise time point matrix between healthy and IPF as in b. (Right) The schematic illustration of the aggregate alignment. Bottom: Heatmap of the smoothened (interpolated) and z-normalized mean log1p gene expression of genes in the EMT pathway along the pseudotime.e, Gene expression of *CAMK1D* between IPF (blue) and healthy (green) data along their pseudotime. Top: interpolated log1p normalized expression (*y*-axis) against pseudotime (*x*-axis) as in c. Bottom: The actual log1p normalized gene expression vs. pseudotime plots.

We inferred pseudotime trajectories for healthy and IPF data using diffusion pseudotime^38^ (**Supplementary** Fig. 2) and aligned them using G2G across 994 highly variable genes (under 13 equispaced time points in [0,1] pseudotime range). The alignment distribution (**Extended Data** Fig. 5a) shows ∼62% mean similarity. The aggregate alignment for all genes showed matches at the beginning and mismatches at the end (**Fig. 5b**) as expected, given that both healthy and IPF lung epithelial differentiation start from AT2 cells, but give rise to AT1 in healthy versus ABC in IPF. Moreover, examining the alignments of ABC-specific marker genes (**Fig. 5c, Supplementary** Fig. 3), we observe a diverging pattern as reported by other studies^27,33,35^, again validating the accuracy of G2G alignment.

We further performed gene set overrepresentation analysis over the top mismatched genes (alignment similarity <40%) and found that Epithelial Mesenchymal Transition (EMT) was the most significantly enriched pathway (**Fig. 5d, Supplementary Table 3**). While the majority of EMT genes show mismatches only at later pseudotime points, consistent with dysregulated EMT being implicated in ABC development in IPF^27,33,34,36,37^, it is interesting to note that some EMT genes also show differences at early/mid differentiation stages (e.g. *NNMT, CXCL1, CXCL8*). These could be potential therapeutic targets to prevent differentiation into the pathological ABC cell state.

Downstream clustering of all gene alignments revealed additional alignment patterns (**Extended Data** Fig. 5b-c). For example, cluster 3 represents genes with almost complete mismatches. Among those, *CAMK1D* (**Fig. 5e)** shows significant upregulation, which is known to be upregulated by TGF-β1^39^ that has a key role in IPF development^40^. Overall, G2G was able to capture the expected alignments, as well as some new early/mid mismatches between the healthy and IPF trajectories.

### *In vivo* - *in vitro* human T cell development comparison using G2G reveals differences in TNFα signaling

We next employed G2G to compare *in vitro* and *in vivo* human T cell development. The thymus is the key site for T cell development in humans, where lymphoid progenitors differentiate through stages of double negative (DN) and double positive (DP) T cells to acquire T cell receptor (TCR) (illustrated in **Fig. 6a** top, **Extended Data** Fig. 6). If the TCR recognizes self antigen presented on MHC during the process of positive selection, the developing T cells further differentiate through abT(entry) cells and finally mature into single positive (SP) T cells. There are different subsets of SP T cells, including CD4+T, CD8+T and regulatory T (Treg) cells, as well as the newly recognized unconventional type 1 and type 3 innate T cells and CD8AA^41,42^. To investigate human T cell development in a model system, we differentiated induced pluripotent stem cells (iPSCs) into mature T cells using the artificial thymic organoid (ATO) system^43^. We previously harvested differentiated cells from week 3, 5, and 7, and reported that the mature T cells in ATO were most similar to the *in vivo* type 1 innate T cells^42^. To explore this further, we performed scRNA-seq analysis of cells harvested at regular intervals throughout differentiation, i.e., including the early time points (**Fig. 6a**, bottom, **Extended Data** Fig. 6a). Cell types were annotated using a combination of logistic regression based predictions from CellTypist^44^ and marker gene analysis (**Extended Data** Fig. 6b**, 7-8**). The ATO system captures the differentiation from stem cells, through mesodermal progenitors and endothelium to the haematopoietic lineage, and then further down to the T cell lineage.

**Fig. 6.**
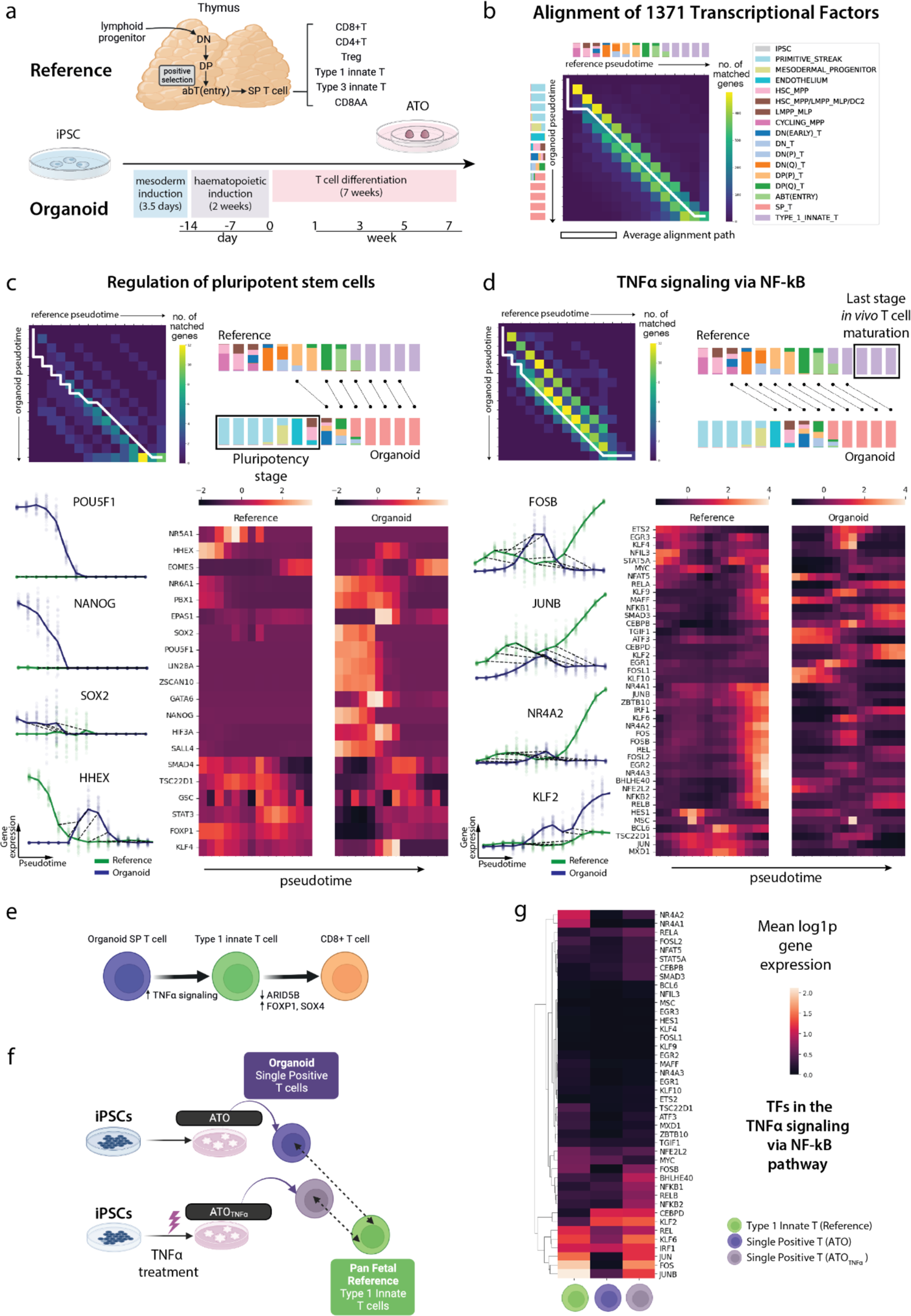
*in vivo, in vitro* human T cell development alignment with G2G. a, Schematic illustration of T cell development in the human thymus. **b,** Aggregate alignment result for all 1371 TFs between *in vitro* organoid (i.e. ATO) and *in vivo* reference (i.e. pan fetal reference from Suo et al. 2022^42^) human T cell development data shown in the pairwise time point matrix between organoid and reference pseudotime. The color represents the number of genes showing match or warp for the given pair of an organoid time point and a reference time point. The white line represents the main average alignment path. The stacked bar plots represent the cell compositions at each time point (14 equispaced time points on pseudotime [0,1]), colored by the cell types, for reference (top) and organoid (left) separately. **c,** Alignment results for all TFs in the pluripotency signaling pathway. Top left: pairwise time point matrix between organoid and reference pseudotime. The color represents the number of genes showing match or warp for the given pair of an organoid time point and a reference time point. The white line represents the main aggregate (average) alignment path. Top right: schematic illustration of the mapping between time points described by the aggregate alignment. The stacked bar plots represent the cell compositions at each time point, colored by the cell types. The black dashed lines represent matches and warps between time points. Boxed segment represents the mismatched pluripotency stage in the organoid. Bottom left: the interpolated log1p normalized expression (*y*-axis) against pseudotime (*x*-axis) for selected genes. Bottom right: heatmap of the smoothened (interpolated) and z-normalized mean gene expression along the pseudotime. **d,** The same plots as in **c** for all TFs in the TNFɑ signaling via NFκB pathway. The boxed segment in the right top plot represents the mismatched last stage *in vivo* T cell maturation. **e,** Schematic illustration of potential targets for further optimization of *in vitro* T cell differentiation towards either type 1 innate T cells or conventional CD8+T cells. **f,** Schematic illustration of the comparison between SP T cells from the wild type ATO and the TNFɑ treated ATO against *in vivo* type 1 innate T cells. SP T cells from the ATO after TNFɑ treatment show more maturity towards *in vivo* type 1 innate T cells. **g**, Heatmap of mean log1p normalized gene expression of TFs within the TNFɑ signaling via NFκB pathway (same gene list as in **d**) in reference (*in vivo* type 1 innate T cells), SP T cells from the wild type ATO, and SP T cells from the TNFɑ-treated ATO (ATO_TNFɑ_).

We then integrated the *in vitro* ATO data with the relevant *in vivo* cell types from our developing human immune atlas^42^ (hereafter referred to as the pan fetal reference) into a common latent embedding using scVI^45^, and estimated their pseudotime (**Extended Data** Fig. 6c-e). For ATO data, the pseudotime was estimated using Gaussian Process Latent Variable Model (GPLVM)^46^ with sampling times as priors. GPLVM has previously been successfully applied in single-cell trajectory inference to incorporate useful priors^47–51^. The pan fetal reference cells’ pseudotime was computed similarly by estimating their time priors from the nearby ATO cells.

Alignment between *in vitro* ATO and *in vivo* pan fetal reference trajectories was performed with G2G (under 14 equispaced time points in [0,1] pseudotime range) using all transcription factor (TF) genes^52^ (1371 TFs), as many TFs function as ‘master regulators’ of cell states and have been used to induce cell differentiation. The aggregate alignment of all TFs showed a mismatch at the beginning and a mismatch at the end (**Fig. 6b**), with ∼66% mean alignment similarity in their distribution (**Extended Data** Fig. 9a).

### Clustering alignments captures biologically interesting groups of genes

Using G2G, the TFs were hierarchically clustered based on their alignments and explored at several sub-optimal resolutions to identify groups with different alignment patterns (**Extended Data** Fig. 9b-c). At low clustering resolution (**Extended Data** Fig. 9c), Cluster 2 captures the pluripotent TFs showing insertions at early pseudotime (**Supplementary Table 5**). The expression of many of them, such as the well-known *POU5F1, NANOG* and *TBX3*^53^, are present in early ATO development, but missing from the reference. This is expected for TFs involved in the regulation of pluripotent stem cells (**Fig. 6c**), since *in vitro* differentiation started from iPSCs, whereas the earliest *in vivo* cells were haematopoietic stem cells (HSCs). Among them, *HHEX*, which is known to be expressed in HSC^54,55^ and early DN T cells^56^ notably demonstrates another pattern: a matching between *in vivo* and *in vitro* HSC and DN T cells as expected, although with the maximum *HHEX* expression in *in vitro* cells being lower than that of *in vivo* cells (**Fig. 6c**). Interestingly, clustering also reveals TFs with mismatches only in the middle time points. (e.g. *POU6F1, SOX18, CSRNP3* in cluster 0 at low resolution, and *BATF2* in cluster 13 at high resolution). This might represent a missing cell state, e.g., *BATF2* is expressed sparsely in endothelial cells which are present in the *in vitro* but not in the *in vivo* system. On the other hand, *LEF1* (necessary for early stages of thymocyte maturation^57^) stands out as a single cluster showing almost 100% matching between the trajectories, while two other clusters include almost 100% mismatching TFs such as *GATA6, SALL4, HOXB6, NACC2,* and *PRDM6*. See **Supplementary** Fig. 4 for gene expression and alignment plots of all aforementioned examples.

### TNFɑ signaling as a potential target for in vitro optimization

We performed gene set overrepresentation among the most mismatched genes (alignment similarity < 40%, **Supplementary Table 4**), and found that genes associated with TNFɑ signaling via NF-ϰB pathway showed significant differences. Many of the TFs in TNFɑ signaling via the NF-ϰB pathway (e.g. *FOSB, JUNB* and *NR4A2*) show an increasing trend at the last stage of *in vivo* T cell development, while this increase is missing in the *in vitro* T cells (**Fig. 6d**). There are exceptions to this overall pattern, e.g., *KLF2*, whose expression is higher *in vitro* than *in vivo* (**Fig. 6d**). This might be due to each gene being regulated by more than one signaling pathway. G2G alignment of all 196 genes in the TNFɑ pathway also consistently identifies a significant mismatch in the last stage (**Extended Data** Fig. 10a), thus suggesting this pathway as a potential target for further *in vitro* optimization. We further validated our results by restricting the analysis to T cell lineages, i.e., DN T cells onwards (**Extended Data** Fig. 10b left). TNFɑ signaling via NF-ϰB pathway remained the most enriched gene set among the mismatched TFs (**Supplementary Table 6**). We remark that although it is possible to recover these differences by doing direct DE analysis between cell subsets, e.g., end products of ATO vs *in vivo* T cells, a key advantage of trajectory alignment is the ability to systematically identify the time point where the mismatch occurred during differentiation. This in turn informs us when to introduce TNFɑ in *in vitro* optimizations.

### In vitro SP T versus in vivo CD8+T lineages

The above alignments were performed using *in vivo* type 1 innate T cells and the relevant precursors, as we previously found that the *in vitro* mature T cells were most similar to the *in vivo* type 1 innate T cells^42^. However, *in vitro* cell differentiation to conventional CD8+T cells might also provide promising routes for cell therapies. We therefore performed another G2G alignment using *in vivo* conventional CD8+T cells and the relevant T lineage precursors (DN T cells onwards). Differences in the two alignment results suggest that potential targets such as *SOX4, FOXP1* and *ARID5B* may tune cells towards *in vivo* CD8+T cells (Details in **Supplementary Note**, Extended Data Fig. 10b-c**, Supplementary Table 7)**.

### Preliminary experiment targeting TNFɑ signaling

Overall, G2G alignment between *in vivo* and *in vitro* human T cell development revealed potential targets for further optimization of *in vitro* T cell differentiation (illustrated in **Fig. 6e**). We experimentally validated the impact of TNFɑ signaling by adding TNFɑ into the ATO media between week 6 to 7 (**Fig. 6f**), and comparing SP T cells of our *in vitro* ATO (wild type) and the TNFɑ treated ATO (ATO_TNFɑ_) to the *in vivo* type 1 innate T cells in the pan fetal reference. We observed that, in the scVI^45^ latent space of all *in vivo* and *in vitro* cells, the euclidean distance between the mean latent dimensions of *in vitro* SP T cells and the *in vivo* type 1 innate T cells decreased by ∼5% after TNFɑ treatment. This suggests that in the latent space, the transcriptomic signature of *in vitro* SP T cells is closer to the *in vivo* transcriptomic signature of type 1 innate T cells after TNFɑ treatment. Further examining the effect, the TFs and all genes within the TNFɑ signaling pathway (**Fig. 6g**, Extended Data Fig. 10d) showed higher expression in ATO_TNFɑ_ T cells compared to the ATO T cells, as expected. TNFɑ treatment also shortened the mean distance of gene expression distributions for all genes and TFs that were significantly distant between ATO T cells and *in vivo* type 1 innate T cells (see **Methods** and **Supplementary Note** for more details on the statistical tests). We also note the change of expression distance in several known SP T cell maturation markers (*IL7R, KLF2, FOXO1, S1PR1, SELL*)^58,59^ (**Extended Data** Fig. 10e-f**, Supplementary** Fig. 6). *IL7R* has shown to initiate in mature SP thymocytes, with its expression dependent on NF-ϰB signaling^60–6364^. The distance of *in vitro* and *in vivo IL7R* expression significantly drops after TNFɑ treatment with an increased expression as expected from mature T cells. *KLF2* has also further upregulated. The rest of the markers maintain expression. Overall, these results suggest more mature SP T cells in ATO_TNFɑ_.

It is worth noting that in another *in vitro* T cell differentiation protocol, TNFɑ was added throughout T cell differentiation to improve T cell production efficiency^65,66^. However, G2G identified that the mismatch in TNFɑ pathway happens at the end of T cell differentiation, and therefore we have added TNFɑ in the last week of differentiation to improve T cell maturation. In conclusion, the target pathway identified by G2G alignment enabled us to successfully push *in vitro* T cells to become more similar to *in vivo* cells (**Fig. 6f**). Our results suggest that targeting TNFɑ is a potential direction to refine the ATO protocol towards mature type 1 innate T cells, subjected to more functional studies in future.

## Discussion

Genes2Genes (G2G) offers a structured framework to align single-cell pseudotime trajectories at single-gene resolution. We validated G2G’s accuracy in identifying matches, mismatches and different alignment patterns on simulated datasets, by benchmarking against the DTW-based trajectory alignment methods, CellAlign^6^ and TrAGEDy^16^. Moreover, using a published dataset^27^, we demonstrated G2G’s potential in identifying genes and pathways that drive pathogenicity in IPF through basaloid cell differentiation. Finally, we captured genes and pathways that can guide the refinement of T cell differentiation in an organoid protocol, and successfully validated that G2G alignments are helpful for inducing the *in vitro* cells to transition towards a more *in vivo* like state.

Given cell-by-gene matrices and pseudotime estimates of a reference and query, G2G generates a five-state alignment string for each gene of interest by running a DP algorithm that unifies DTW and gap modeling. The gene sets to compare can be either all genes, or restricted to gene sets of interest, e.g., TFs, regulons, highly variable genes, or genes associated with a certain biological/signaling pathway. G2G outperforms existing methods through more descriptive and accurate alignments, highlighting both matched (including warps) and mismatched regions of a gene’s expression over time. It provides a powerful addition to the current repertoire of comparative analysis toolboxes for any pseudotime alignment task, e.g., *in vivo*/*in vitro*, healthy/disease, treatment/control, cross-species etc.

An important feature of G2G is gene-specific alignments. Most existing approaches produce a single alignment for all genes by computing high-dimensional Euclidean distances over their mean gene expression vectors. Such metrics suffer from ‘the curse of dimensionality’ by losing accuracy as the number of genes increases^67^. Importantly, in many contexts, an alignment across all genes masks gene heterogeneity along trajectories in the reference and query systems. Alpert et al (2018)^6^ discuss how to choose the right alignment resolution, recommending alignment of the largest gene set that shows significant differential expression over time. This is to remove stably expressed genes which may add noise and skew the alignment results. Our method goes further and fully resolves all gene groups with distinct matching and mismatching patterns at different stages along trajectories.

G2G alignment allows users to identify clusters of genes with broadly similar alignments. We show that pathway overrepresentation analysis on each gene cluster can reveal specific biological signaling pathways driving the differences in pseudotime trajectories at different stages. These pathways and gene sets can be a starting point for protocol intervention strategies in the case of *in vivo*/*in vitro* alignments, and for mechanistic molecular interpretation of differences between trajectories in other cases. Our work provides proof-of-concept by demonstrating the power of gene-level alignment to gain biological insights (e.g. discovering differential genes and their associated pathways along pseudotime) for future investigation.

The reliability of a trajectory alignment depends on how trustworthy the given pseudotime estimates are. Thus we recommend the user to select a reliable method suitable for their datasets to infer pseudotime trajectories^2^. Future work is also needed to develop suitable methods to calibrate G2G’s input (i.e. pseudotime estimates and interpolation). For instance, an adaptive window size for the Gaussian kernel-based interpolation may optimize the method’s sensitivity to the variance of expression in nearby cells. Furthermore, the current G2G version can only compare two linear trajectories without considering branching processes. We are aware of other efforts in aligning branched trajectories with DTW based tree alignment^9^. Output from such alignments, i.e., identified pairs of correspondences, could be input into G2G for a comprehensive pairwise lineage alignment to capture mismatches.

In summary, G2G is a formal trajectory alignment framework for scRNA-seq data, and is able to capture matches and mismatches accurately at gene-resolution. It enables deeper understanding of the diversity of gene alignments between single-cell datasets. The G2G package is easy-to-use and freely available at https://github.com/Teichlab/Genes2Genes with a tutorial. We have demonstrated that regenerative medicine can specifically benefit from such trajectory comparisons by extracting cues to guide refinement of *in vitro* cell engineering to recapitulate *in vivo* development. We envision that the software will be useful to the community for exploring other biological scenarios such as cell activation/stimulation responses in control and disease, generating new insights to advance our understanding of cell development and function in health and disease.

## Methods

### Genes2Genes (G2G): A new alignment framework for single-cell trajectories

Dynamic programming (DP) remains central to many sequence alignment algorithms. As described in the main text, Genes2Genes performs DP alignment independently for all genes of interest, between a reference trajectory *R* and a query trajectory *Q*. In other words, each gene-level (i.e. gene-specific) trajectory alignment is an independent DP task of pairwise time series alignment. The aim is to generate an optimal sequence of matched time point pairs and mismatched time point pairs between *R* and *Q* for each gene. As illustrated in **Fig. 1b**, there are five different alignment states which denote these matches and mismatches between two time points. For each time point in any gene trajectory, there is a respective distribution of gene expression, as explained by an observed dataset of single-cell (scRNA-seq) measurements. G2G evaluates the similarities of these reference and query expression distributions over time, to determine matches and mismatches between their time points. The following sections introduce the problem of pairwise time series alignment, and describe the main components of our framework (**Fig. 2**).

### Pairwise time series alignment for trajectory comparison

A trajectory is a continuous path of change through some feature space, along some axis of progression (such as time)^68^. In single-cell transcriptomics, this feature space is usually defined by genes, and a trajectory through a high-dimensional gene space can describe the transcriptomic state of a cell as a function of time. A temporal (e.g. pseudotime) ordering of a set of single cells represents a discretization of the respective cell state trajectory, and their entire gene expression dataset forms a multivariate time series. On the other hand, their expression of a single gene forms a univariate time series. In this work, we consider a pairwise alignment of univariate time series, which allows us to perform gene-specific trajectory alignment.

Given two time series (sequences), reference *R* and query *Q* of length (i.e. a finite number of time points) |*R*| and |*Q*|, their pairwise alignment describes a sequential mapping between their time points. As an optimization problem, computational alignment has two key properties: (1) an optimal substructure, and (2) overlapping set of subproblems, which make it dynamic programmable (DP)^4^. Property (1) means, the optimal alignment of any two prefixes *R_1_*_..j_ and *Q_1_*_..i_ depends on the optimality of three sub-alignments: (i) *R_1_*_..j\**1*_ and *Q_1_*_..i\**1*_, (ii) *R_1_*_..j\**1*_ and *Q_1_*_..i_, and (iii) *R_1_*_..j_ and *Q_1_*_..i\**1*_. Property (2) means, there exists subproblems (prefix alignments) that are overlapping. DP begins optimizing the alignment of prefixes, starting from null (Φ) sequences until it completes an alignment of the entire two sequences. During this process, it computes overlapping subproblems only once and reuses them through a memoization (history) matrix *H*. In the standard DP alignment algorithm, any cell (*i, j*) in *H* stores the optimal alignment cost of the two prefix sequences: *R_1_*_:j_ and *Q_1_*_:i_, by optimizing an objective function which quantifies the alignment through a set of recurrence relations. Once all *H* matrix cells are computed, the optimal alignment can be retrieved by backtracking, starting from the right-most bottom cell (|*Q*| + *1*, |*R*| + *1*) until reaching the matrix cell (*0*,*0*). The time complexity of this algorithm depends on how its alignment scoring scheme is designed. (The standard scheme has a quadratic complexity to find the best alignment out of the factorially growing all possible number of alignments).

### Preprocessing a trajectory time series by distributional interpolation

Interpolation is a necessary preprocessing step that a time series has to undergo prior to taking part in an alignment. This is to ensure smoothly changing and uniformly distributed data that are in phase (i.e. having the same rate of sampling) at least approximately; otherwise a reliable alignment cannot be guaranteed^6,12^. Here we chose to extend the mean gene expression based interpolation method used by CellAlign^6^ to a distributional interpolation, for preprocessing a reference and query time series of gene expression before their alignment.

Given a pseudotime series *t* of (log1p normalized) expression in some gene *g*_j_ of a single-cell dataset, our distributional interpolation method first min-max-normalizes the pseudotime of *t* as to be in the range of [*0*,*1*]. Then, *m* equally spaced artificial (interpolated) time points are determined within [*0*,*1*], where for each artificial time point *t′*, we estimate a Gaussian distribution (of mean *g*_j_(*t′*)_*mean*_ and standard deviation *g*_j_(*t′*)_*std*_) by using the Gaussian kernel-based weighted approach. For each cell *i* annotated with pseudotime *t*_i_, an associated weight is computed w.r.t each artificial time point *t′* as:

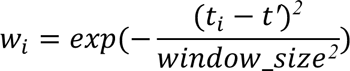

where default *window_size* = *0.1*. The below equations are then used to compute the Gaussian distribution parameters *g*_j_(*t*′)_*mean*_ and *g*_j_(*t*′)_*std*_:

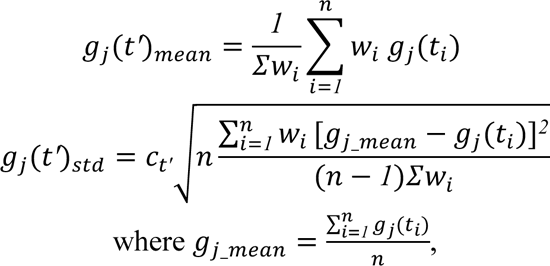

*n* is the total number of cells, and *c*_*t*_*_′_* is the weighted cell density (abundance of cells) at the interpolated time point *t′*. The weighted cell density computed as:

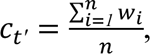

is the expected weight of a cell at *t′*. A higher expected weight is indicative of a higher cell density. This way, we account for cell abundance when deciding the variance for the interpolated point (otherwise with a very low number of cells, we may get a very high variance). Next, we generate *k* = *50* random data points from the Gaussian distribution *N*(*g*_j_(*t′*)_*mean*_, *g*_j_(*t′*)_*std*_) for each interpolated time *t′*, representing the interpolated distribution of single-cell gene expression. Note: In this work, we use a fixed number of interpolated time points *m* for both reference and query. As *m* controls the resolution of the alignment, it should suffice to represent the entire trajectory. This interpolation also adds an O(*nm*) time complexity due to taking a weighted contribution from all cells at each *t*′. To overcome this, a general solution is to subsample datasets before the analysis and/or to reduce the number of contributing cells by considering only the nearest neighborhood.

The reference pseudotime series and query pseudotime series of each gene is preprocessed using the above described distributional interpolation method. The interpolated time series are then input to our G2G dynamic programming algorithm as detailed in the next section.

### A new dynamic programming algorithm for time series alignment of a single gene

Here we describe a new dynamic programming (DP) algorithm to generate an alignment between a reference time series *R* and query time series *Q* of log-normalized expression of a specified gene. This algorithm jointly adapts Gotoh’s sequence alignment^15^ with classical dynamic time warping (DTW)^19^ to accommodate five states of alignment (**Fig. 1b**), i.e., one-to-one match (`M`), many-to-one warp (`W`), one-to-many warp (`V`), insertion (`I`), and deletion (D) between time points in the two time series. We denote the five-state space as Ω = [M, W, V, D, I]. Our approach unifies matches and mismatches within a single DP algorithm unlike DTW which only handles matches (including warps).

The DTW algorithm originated from the speech recognition domain^19^ under the family of dynamic programming algorithms for optimization^4^. It has been extensively used to align time series with shifts (warps). Sankoff and Kruskal (1983)^12^ had previously discussed how to capture warps and indels both from a single alignment algorithm. They provided a DP recurrence relation involving evaluations of the five alignment states to decide the optimal state for each pair of *R* and *Q* time points when aligning two time series. Extending this idea further, we implemented the Gotoh’s O(|*R*||*Q*|) DP algorithm to generate an optimal five-state alignment hypothesis for *R* and *Q* time series by:

● Defining a Bayesian information-theoretic measure of distance between two gene expression distributions using the minimum message length statistical inductive inference framework^12,24^.
● Defining a five-state machine that models state transitions in the alignment hypothesis across the five different matching and mismatching states, defining one-to-one matches, warps (i.e. many-to-one compression matches and one-to-many expansion matches), and insertions-deletions (indels).

The scoring scheme of the DP algorithm evaluates every pair of reference time point *j* and query time point *i* to generate an optimal alignment across all time points. This involves computing two types of cost: (1) the cost of matching the two time points, *j* and *i* (denoted by *Cost*_*match*_(*i, j*)) based on their respective (interpolated) gene expression distributions, and (1) the cost of assigning an alignment state *x* ∈ Ω for the two time points *j* and *i*. The following sections first detail on how we compute these costs, and then describe how our DP optimization works using this scoring scheme.

Note: The reference time point *j* and query time point *i* are also denoted by *R*_j_ and *Q*_i_ in the main text.

### The DP scoring scheme

The cost of match between the reference time point *j* and query time point *i*

We expect a match between the reference time point *j* and query time point *i* if they have similar distributions of gene expression. Thus, to score the likelihood of a match, we define a distance measure between the two gene expression distributions corresponding to the *j* and *i* time points, respectively. To compute this distance, we first take the interpolated single-cell expression datasets at time point *j* of *R* (denoted by *R*(*j*)) and time point *i* of *Q* (denoted by *Q*(*i*)). We already know the mean (μ) and standard deviation (σ) statistics for *R*(*j*) and *Q*(*i*) separately, as they were estimated during the time series interpolation step. Thus we define the Gaussian distribution *N*(μ_*R*(j)_, σ_*R*(j)_) for *R*(*j*), and the Gaussian distribution *N*(μ_&(i)_, σ_&(i)_) for *Q*(*i*) using their respective μ and σ statistics. Accordingly, if *D*_*R*(j)_ = [*d*_1_, *d*_2_,…, *d*_|*R*(j)|_] and *D*_&(i)_ = [*d*_1_, *d*_2_,…, *d*_|&(i)|_] are the expression data vectors of the *R*(*j*) and *Q*(*i*) datasets, respectively, then:

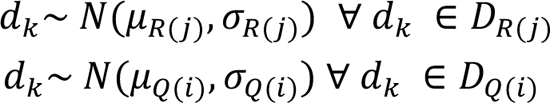

To be brief, we denote *N*(μ_*R*(j)_, σ_*R*(j)_) distribution by *N*_*R*(j)_, and *N*(μ_&(i)_, σ_&(i)_) by *N*_&(i)_.

Next, we implement the cost function, *Cost*_*match*_(*i, j*), to consider both:

1. **Data:** *D*_*R*(j)_ and *D*_&(i)_ expression vectors of the *R*(*j*) and *Q*(*i*) datasets, respectively,
2. 2. Models: Gaussian distributions, *N*_*R*(j)_ and *N*_&(i)_,

when computing the distance between *R*(*j*) and *Q*(*i*). To do so, we use the minimum message length (MML) criterion^24,25^ and define a Bayesian information-theoretic distance measure. Fig. 2 (top left) illustrates an abstract overview of our MML framework and its place in the overall G2G alignment framework. Supplementary Fig. 1a further expands this illustration to explain how the *Cost*_*match*_(*i, j*) computation works for a pair of reference time point *j* and query time point *i*, as detailed by the following sections.

### Primer on minimum message length inference (MML)

MML is an inductive inference paradigm for model comparison and selection, grounded on Bayesian statistics, information and coding theory. It facilitates designing hypothesis test schemes specific to a problem domain. Given a hypothesis (model) *H* and some data *D*, it lays an imaginary message transmission from a sender who jointly encodes *H* and *D*, aiming for their lossless decoding at a recipients’ side. Bayes theorem defines their joint probability as:

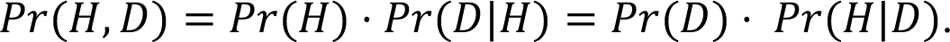

Separately, Shannon information^28^ defines the optimal length of a message that encodes some event *E* with a probability *Pr*(*E*) as:

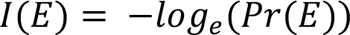

measured in nits, where *I* denotes information. By applying the Shannon information^28^ to Bayes theorem, we can map the respective probability elements in *Pr*(*H*, *D*) = *Pr*(*H*) ⋅ *Pr*(*D*|*H*) onto the information space, describing the amount of Shannon information needed to encode *H* and *D* jointly as:

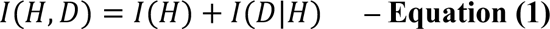

This gives a two-part total message length of encoding *H* and *D* jointly. The first part *I*(*H*) refers to the message length of encoding the hypothesis *H* itself, whereas the second part *I*(*D*|*H*) refers to the message length of encoding the data points in *D* using *H*.

When there are two hypotheses, *H*_1_ and *H*_2_, that describe the same data *D*, MML enables us to select the best hypothesis that gives a model-complexity vs. model-fit tradeoff, by evaluating a compression statistic Δ = *I*(*H*_1_, *D*) − *I*(*H*_2_, *D*). Here, Δ is also the log odds posterior ratio between the two hypotheses.

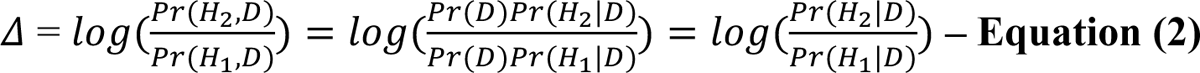

If Δ>0, this implies that the hypothesis *H*_2_ is *e*^C^ times more likely than *H*_1_, and vice versa.

### Casting the cost of matching between reference time point *j* and query time point *i* under MML

Given the expression data *D* (containing both *D*_*R*(j)_ and *D*_&(i)_) and their estimated Gaussian distributions (*N*(μ_*R*(j)_, σ_*R*(j)_) and *N*(μ_&(i)_, σ_&(i)_) denoted by *N*_*R*(j)_ and *N*_&(i)_, respectively), we formulate two different hypotheses:

1. **Hypothesis A:** assumes that the two time points match, and thus explains data *D* with a single, representative Gaussian distribution *N*(μ_∗_, σ_∗_) denoted by *N*_∗_ (which is either ^*N*^*R*(j) ^or *N*^&(i)^).^
2. **Hypothesis** Φ: assumes that the two time points mismatch, and thus explains data *D*_*R*(j)_ with *N*_*R*(j)_, and data *D*_&(i)_ with *N*_&(i)_, independently.

We then compute the two message lengths: *I*(*A, D*) and *I*(Φ, *D*) according to Equation 1 in the above described MML formulation:

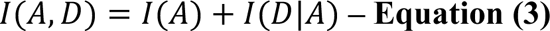

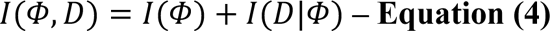

where, *A* = [*N*_∗_] and Φ = [*N*_*R*(j)_, *N*_&(i)_]. *I*(*A, D*) is the total message length of encoding *A* and *D* jointly, where *I*(*A*) refers to the encoding length of the parameters μ_∗_, σ_∗_ of *N*_∗_distribution. *I*(*D*|*A*) refers to the encoding length of all data points in *D* based on their likelihood under *N*(μ_∗_, σ_∗_). Accordingly, we can re-write and expand Equation 3 to:

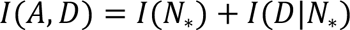

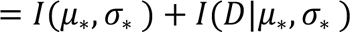

On the other hand, *I*(Φ, *D*) is the total message length of encoding Φ and *D* jointly, where *I*(Φ) refers to the sum of the independent encoding lengths of parameters μ_*R*(j)_, σ_*R*(j)_ of *N*_*R*(j)_ and μ_&(i)_, σ_&(i)_ of *N*_&(i)_. *I*(*D*|Φ) refers to the sum of the independent encoding lengths of all data points in *D*_*R*(j)_ and all data points in *D*_&(i)_ based on their likelihood under their respective Gaussian distributions: *N*(μ_*R*(j)_, σ_*R*(j)_) and *N*(μ_&(i)_, σ_&(i)_). Accordingly, we can re-write and expand **Equation 4** to:

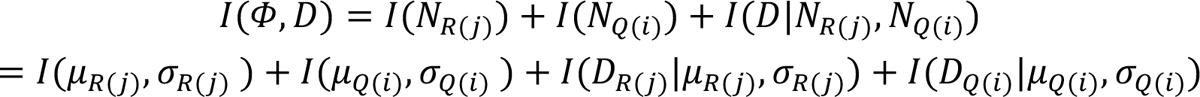

Note: See the next section for the equations used to compute each term in Equation 3 and Equation 4.

Next, we normalize each total message length to compute a per datum message length (i.e. entropy), by dividing them by the total number of datapoints (single-cells) in *D*.

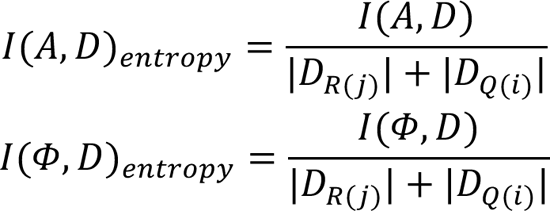

Note: to make the *I*(*A, D*)_*entropy*_ measure symmetric, we take the average:

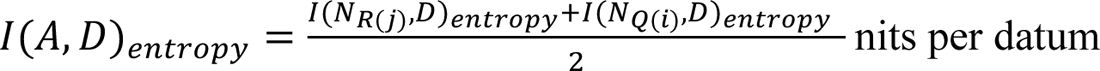

Afterwards, we compute a compression statistic Δ, which is taken as our *Cost*_*match*_(*i, j*):

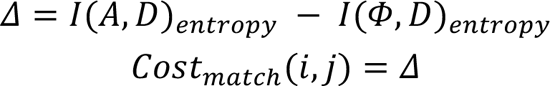

As in Equation 2, this reflects the log odds posterior ratio:

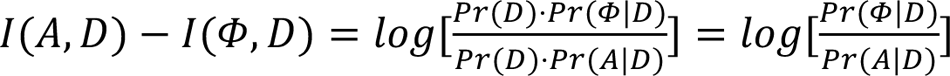

When *R*(*j*) and *Q*(*i*) are very dissimilar, the total encoding length under hypothesis A (i.e. the time points match) results in a larger value compared to that of hypothesis Φ (i.e. the time points mismatch). Thus, *Cost*_*match*_(*i, j*) increases as the distributions deviate from each other (Extended Data Fig. 1b, Supplementary Fig. 1b,c).

### Computing the total encoding message length for any Gaussian model *N*_∗_ and data *D* of size *N*

The above described *Cost*_*match*_(*i, j*) distance measure is computed using the standard MML Wallace Freeman approximation^24,28^ defined for a Gaussian distribution^25,69^. As defined by Equation 1, for any dataset *D* and a hypothesis *H* that describes *D* = [*x*_1_, *x*_2_,…, *x*_*N*_] under a Gaussian distribution *N*(μ, σ) with parameters ^θ⃗^ = (μ, σ), the total message length of encoding *H* and *D* jointly is given by:

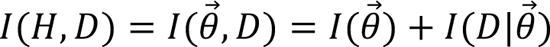

MML Wallace Freeman approximation expands this to:

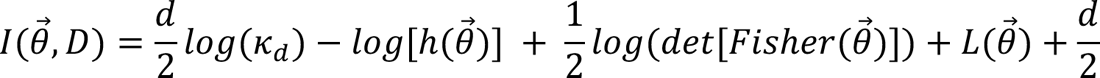

where *d* is the number of free parameters (*d* = 2 for a Gaussian), and *κ_*d*_* is the Conway lattice constant^70^ for k_d_ is 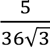 = 2). Here: ℎ(<sup>θ⃗) is the prior over the parameters. μ is defined with a uniform prior over a predefined range of length *R*_μ_. *log*(σ) is defined with a uniform prior over a predefined range of length *R*_σ_. Accordingly,

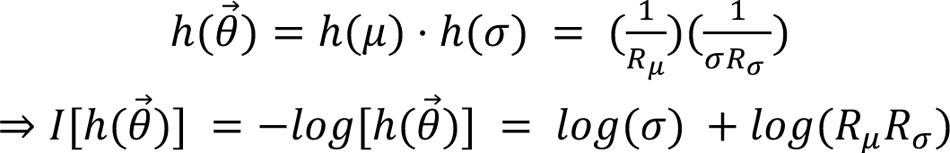

We use *R*_μ_=15.0 and *R*_σ_= 3.0 as reasonable for log normalized gene expression data (e.g. across 20,240 genes, we observe ∼8.1 maximum log normalized expression and ∼1.7 maximum standard deviation in the pan fetal reference dataset).

*L*(^θ⃗^) is the negative log likelihood:

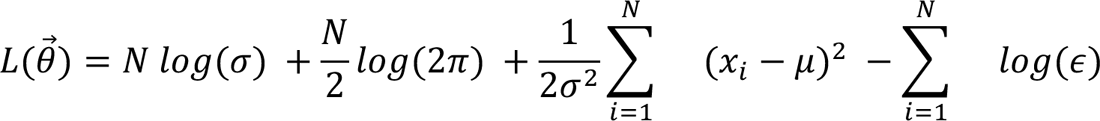

where, ∈ is the precision of measurement for each data point (taken as ε=0.001). *det*[*Fisher*(^θ⃗^)] is the determinant of the expected Fisher matrix (i.e. the matrix of the expected second derivatives of the negative log likelihood function). This determinant has the closed form: 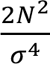

### The cost of alignment state assignment for the reference time point *j* and query time point *i*

The DP scoring scheme also involves computing a cost of assigning a certain alignment state *x* ∈ Ω = [M, W, V, D, I] for the two time points *j* and *i*. This state assignment cost is computed as the amount of Shannon information^28^ required to encode state *x* given the assigned state *y* for the preceding time points. As previously said, in information theory, Shannon information defines the optimal length of a message that encodes some event *E* with a probability *Pr*(*E*) as: *I*(*E*) = −*log*_*e*_(*Pr*(*E*)) measured in nits, where *I* denotes information. Accordingly, the cost of assigning state *x* given a previous state *y* is: *I*(*x*|*y*) = −*log*_*e*_(*Pr*(*x*|*y*)). We define a five-state machine (middle left of Fig. 2) to explain these conditional probabilities of state assignments (a.k.a state transitions).

This finite state machine extends the general three-state alignment machine^20,21^ which has a match (M) state, delete (D) state, and insert (I) state, by adding two new states: compression warp (W) state and expansion warp (V) state (Fig. 1b). The warp states are equivalent to the match state but are extensions to accommodate one-to-many and many-to-one matches between the two series, respectively. As in the three-state alignment machine^22^, we enforce symmetry, while prohibiting an invalid transition from an indel state to a warp state. That is, we do not allow I→W and D→V, as they can be covered by a single M state in the first place. On the other hand, we have the choice of allowing D→W and I→V, as there can be a legitimate case of a warp match after an insertion or deletion. Note: all the outgoing transitions of each state in this finite state machine add up to a probability of 1. It also treats <I and D> and <W and V> equivalently. Consequently, there are 23 total number of state transitions in this machine, yet there are only three free transition probability parameters: [*Pr*(M|M), *Pr*(I|I), *Pr*(M|I)], due to the symmetry and characteristics of the machine. These probabilities control the expected lengths of a match and a mismatch (analogous to how an affine gap penalty function works in sequence alignment). In this work, we have chosen [*Pr*(M|M)=0.99, *Pr*(I|I)=0.1, *Pr*(M|I)=0.7] as the optimal parameter setting based on a grid search which minimized the alignment inaccuracy rate in our dataset of simulated experiment 1. We also note that these parameters can be automatically inferred using an added layer of optimization and time complexity on top of the main DP optimization, which is an interesting future direction to follow.

Altogether, our G2G DP scoring scheme utilizes the *Cost*_*matc*h_(*i, j*) function, and the above described state assignment costs (i.e. all possible state transition costs evaluated as *I*(*x*|*y*) for all possible *y* → *x* state transitions under the five-state machine), which are used to define the recurrence relations of our DP algorithm.

### Dynamic programming recurrence relations

We formulate the DP problem using five history matrices [*Hist*_*M*_, *Hist*_*W*_, *Hist*_V_, *Hist*_*D*_, *Hist*_*I*_], where each matrix corresponds to each alignment state in Ω, respectively. Any history matrix, *Hist*_*x*_ for state *x* ∈ Ω, has the dimensions (|*Q*| + *1* × |*R*| + *1*). In other words, the columns correspond to the time points in the reference series *R*, while the rows correspond to the time points in the query series *Q*. Each cell *Hist*_*x*_(*i, j*) stores the optimal cost of aligning the prefix time series *R*[*1*.. *j*] and *Q*[*1*.. *i*] ending in state *x*. The DP recurrence relations to compute each matrix cell (*i, j*) of each history matrix for *i* > *0, j* > *0* are:

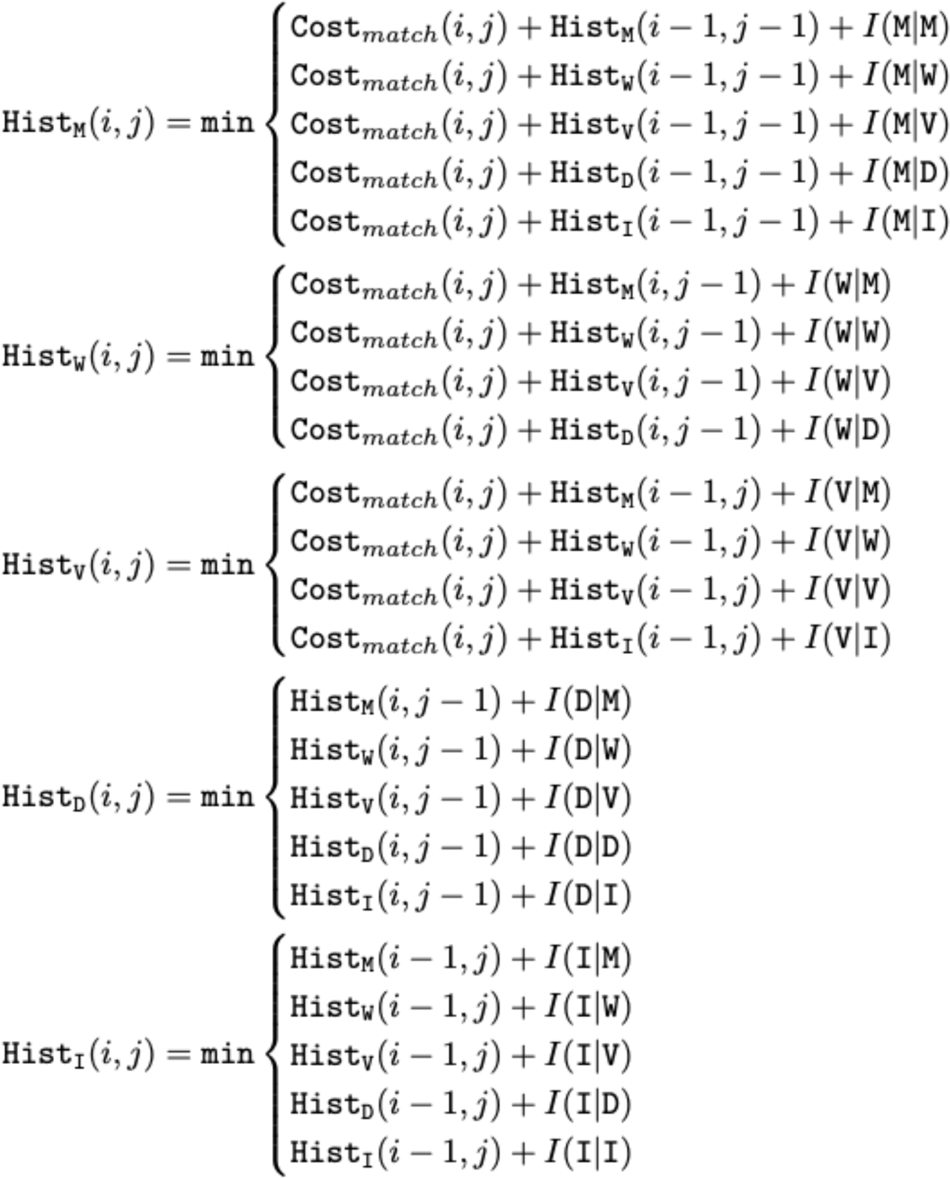

As previously described, *Cost*_*matc*_*_*h*_*(*i, j*) measures the distance between the two interpolated gene expression distributions corresponding to the reference time point *j* and query time point *i*. The cost term *I*(*x*|*y*) ∀ *x*, *y* ∈ Ω refers to the Shannon information of a state transition *y* → *x* (e.g. *I*(`I|M`) is the cost of `M`→`I`, computed as −*log*_*e*_[*Pr*(`I|M`)]), based on the five-state alignment machine as explained before. Prior to computing the above relations, the history matrices are initialized as:

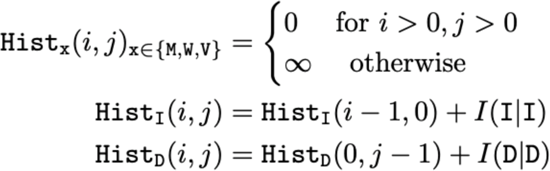

Note: For the cases of <*i* = *1* and *j* = *1*> (i.e. before the first state transition), we assign a uniform transition cost: *I*(`M`) = *I*(`I`) = *I*(`D`) = −*log*_*e*_(*1*/*3*). All the five history matrices are computed by running the aforementioned DP algorithm. We then generate the optimal alignment *Y*^∗^ as a five-state string by backtracking, starting from the cell:

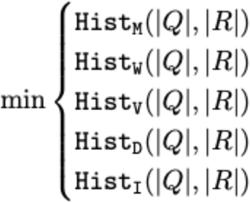

Note: The optimal alignment cost landscape matrix *L* can be visualized by constructing:

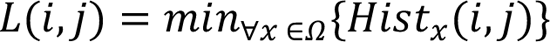

Let *Y*^∗^ be the optimal alignment string between *R* and *Q* time series that minimizes the total DP alignment cost under our DP scoring scheme and recurrence relations. In other words, *Y*^∗^ describes the optimal set of reference time point and query time point pairs that are matched, as well as the optimal set of reference time points and query time points that are mismatched. The *k*th character in *Y*^∗^, i.e., *Y*^∗^[*k*], gives the alignment state for the corresponding reference and query time points (*Y*^∗^[*k*] ∈ Ω =[M, W, V, D, I]). Let the set of matched time point pairs (*i, j*) in *Y*^∗^ be denoted by *T*_*matched*_. Then, the total alignment cost of *Y*^∗^ is the sum of the total match cost (*C*_*matc*h_) and the total state assignment cost (*C*_*state*_), where:

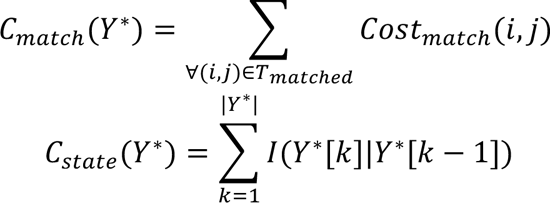

under our scoring scheme.

Overall, the optimal alignment *Y*^∗^ is generated by optimizing the following objective function:

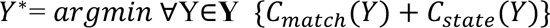

where Y is the space of all possible five-state alignment hypotheses.

#### *Note on using a custom Cost*_*matc*h_(*i, j*) *function*

Our MML-based *Cost*_*matc*h_(*i, j*) function defines a distribution-based distance measure to compute the cost of matching the reference time point *j* and query time point *i* based on their gene expression distributions (as explained in the previous sections). Considering expression distributions rather than just the mean expression values allows us to make technical/batch variations implicit. However, we note that this can be any cost function (e.g. KL-divergence) which can measure the distance between two Gaussian distributions. However, our MML-based compression statistic Δ enables us to define a complete description of each hypothesis, which considers both model-complexity and data-fit. On the other hand, KL-divergence is equivalent to the expected log-likelihood ratio, which does not take the complexity of model parameters into account.

#### Note on aligning extreme cases

The underlying assumption during our time series interpolation step before alignment is that each trajectory has a smooth trend of change (i.e. no abrupt changes). However, in reality, this may not always be the case. For instance, some pluripotent genes are highly expressed in early stages of development, but suddenly drop expression to zero mid-way. In such cases, the aforementioned assumption breaks down and thus, the interpolated variance may not reflect the true observation. In addition, when a gene is completely or almost zero expressed, there is no distribution to model. Hence, to handle such extreme cases where the basic assumptions break down, G2G applies the following steps:

– When interpolating a gene trajectory (either reference or query), check if there are significantly inconsistent, zero expressed regions in the trajectory by running a sliding-window scanning to identify interpolation pseudotime points with almost zero expression (< 3 cells expressed as per the standard check). A region in the trajectory is taken as significantly inconsistent if the length of the adjacent zero expressed pseudotime region exceeds 0.2. Then apply a common standard deviation (which is 10% of the minimum standard deviation estimated for all interpolation pseudotime points in the trajectory) to represent those inconsistent regions (such that those regions will always have a very low standard deviation with complete or almost zero mean expression).
– If both reference and query genes are zero-expressed, they are identical and completely matching.
– If either of the reference or query gene is almost zero expressed (< 3 cells expressed as per the standard check) across pseudotime, apply 10% of the minimum standard deviation estimated for all interpolation pseudotime points in the other trajectory as the standard deviation for the extreme-case trajectory, and vice versa.

#### Reporting alignment statistics over gene-level alignments

G2G generates alignments for all genes of interest by running the DP alignment algorithm (detailed in the previous section) independently for each gene between the reference and query. Each optimal alignment output is a five-state alignment string describing the matches and mismatches. These gene-level alignments are further analyzed to generate useful statistics and insights as below.

#### Distribution of Alignment similarities

The percentage of total matching (i.e. one-to-one matches and warps) (termed as ‘alignment similarity’ percentage) in each five-state gene-alignment string, as well as its average across all genes, provide quantitative measures of the degree of concordance between the reference and query trajectories. The genes can be ranked from the most distant to most similar genes across time using the alignment similarity percentage.

#### Aggregate alignment

G2G also generates a single aggregate (average) alignment across all genes (or any specified subset of genes) using each of their optimal alignment landscapes. Recall that any matrix cell (*i, j*) in the optimal alignment landscape (i.e. *L*(*i, j*) = *min*_∀*x* ∈A_{*Hist*_*x*_(*i, j*)}) refers to the optimal ending alignment state of the prefix time series *R_1_*_..j_ and *Q_1_*_..i_. Thus, across such matrices of all genes, there is a five-state frequency distribution for each (*i, j*). To generate an average alignment, we start a traversal from the right-most bottom cell (|*Q*| + *1*, |*R*| + *1*), and choose the most probable alignment state *x* ∈ Ω for *R*_|*R*|_ and *Q*_|&|_ time points as the most frequent state across all genes. According to this state, we traverse to the next matrix cell (i.e. if it is *m*, we go to (*i* − *1, j* − *1*); if it is *d*, we go to (*i, j* − *1*) and so on). By the time we reach the cell (*0*,*0*), we have a representative five-state alignment string.

#### Clustering alignment patterns

Given a set of five-state alignment strings (i.e. gene-specific alignments), we employ a string clustering approach to identify groups of genes that show similar temporal matching and mismatching patterns. Given a pairwise distance matrix between all alignment strings, computed using a suitable string distance measure, we run the standard agglomerative hierarchical clustering algorithm (under the average linkage method; using the Python *sklearn.cluster* package). The distance threshold parameter for the linkage distance in hierarchical clustering controls the level at which the cluster merge stops, allowing inspection of the clusters at different levels of the clustering hierarchy.

#### Defining a distance measure for five-state alignment strings

Clustering gene alignments require defining a distance measure between two alignment paths. While the polygonal area based distance measure^71^ is ideal for three-state alignment strings, it is unable to distinguish between warps and indels. The commonly used string distance measures are the Hamming distance (for equal-length strings) and the Levenstein distance (for any-length strings). Levenstein distance is the minimum number of edits (i.e. character substitutions, inserts and deletes) needed to transform one string to another, computed using dynamic programming. G2G computes the pairwise Levenstein distances between 5-state alignment strings (using the *leven* python package) normalized by the maximum length of each pair of strings in comparison. On the other hand, Hamming distance is the minimum number of single-character substitutions needed to transform one string to another. G2G can compute the pairwise Hamming distances using *scipy.spatial.distance.cdist*, by first encoding the five-state alignment strings as equal-length binary strings (**Supplementary** Fig 5a). We define a binary encoding scheme that transforms each five-state alignment string into a binary vector of size |*R*| + |*Q*|. This is done by traversing through the alignment path, recording for each trajectory, the match/mismatch state of their respective pseudotime points (i.e a match state *x* ∈ [*m, w*_*d*_, *w*_i_] is encoded by 1; a mismatch state *x* ∈ [*i, d*] is encoded by 0). The resultant binary strings of *R* and *Q* are then concatenated to numerically represent their alignment path. Note: Both Levenstein and Hamming distances are normalized to be in the range [0,1].

#### Choosing the right string distance measure

We tested the clusterings under both metrics, Levenshtein distance and Hamming distance, to choose the one which gives the best grouping of genes. In hierarchical clustering, the number of clusters decreases as the distance threshold increases. Ideally, the bottom level of an optimal hierarchical structure of string clusters should represent distinct clusters for all unique strings in the given string set. In other words, the maximum number of gene clusters (at the minimum distance threshold) should be equal to the number of unique alignment strings in all alignment strings. Thus in general, this is a good criterion to evaluate between different distance measures.

Accordingly, we inspected the difference in clustering structures (at the minimum distance threshold) resulting from Levenstein distance versus Hamming distance across the datasets (**Supplementary** Fig 5b). We found that the minimum distance threshold under the Levenstein distance always gives the highest maximum number of clusters equal to the number of unique alignment strings, whereas the Hamming distance does not always guarantee such capture. This overall observation agrees with the theoretical expectation that the Levenstein distance can better distinguish the differences in all 5 states, compared to Hamming distance which can only distinguish between matches and mismatches. Therefore, we recommend using Levenstein distance for alignment string clustering.

#### Choosing the distance threshold for hierarchical clustering

A common strategy to select the optimal distance threshold of a hierarchical clustering is to optimize the distance based on the mean silhouette coefficient ^72^ across all data samples.

Given cluster assignments of a set of data samples, the silhouette coefficient score is computed for each sample as the difference between the mean intra-cluster distance (i.e. the mean distance from the sample to all other samples in the same cluster) and the mean nearest-cluster distance (i.e. the minimum of the mean distances from the sample to all samples in every other cluster), normalized by the maximum among those two distance measures. The range of Silhouette coefficients is [-1,1], where a high positive value indicates well separated clusters, a low positive value closer to 0 indicates overlapping clusters, and a low negative value indicates incorrect assignments. To empirically determine an appropriate distance threshold, we obtain clusterings for all eligible thresholds in the range (0, 1.0] with 0.01 step size, and compute their mean Silhouette coefficients using *sklearn.metrics.silhouette_score*.

#### Hierarchical clustering of alignment strings requires a tradeoff

In general, the best clustering structure is considered as the one with the highest mean Silhouette Coefficient. However, for strings, we observe that this value is given by the maximum possible number of clusters (equal to the number of unique alignment strings). In trajectory alignment, a large number of unique alignment patterns are possible due to subtle differences in the optimal alignment states across their pseudotime bins for different genes. For instance, in our simulated dataset, there are 113 unique alignment strings covering the 7 major alignment patterns (i.e. *Match, Divergence* and *Convergence* under 3 different bifurcation points), and the size of their clustering structure with the highest mean Silhouette Coefficient is always equal to their number of unique alignment strings (Extended Data Fig. 2c).

Our objective is to obtain a less noisy, biologically interpretable clustering structure. Thus, this involves selecting the distance threshold which gives a good tradeoff between mean silhouette score and the number of clusters capturing the main alignment patterns. In our simulated dataset, we observe that this tradeoff is given by the distance threshold that corresponds to the second highest locally optimal mean silhouette score. The distance threshold 0.22 gives the second highest locally optimal mean silhouette score of 0.82 with 15 clusters, including the 7 major clusters with only a 0.1% mis-clustering rate (the percentage of the total number of outliers in all clusters) (**Fig. 3e**). The rest of the clusters are mini clusters covering 31 (0.8%) alignments separated due to noise such as warps.

Accordingly, we conclude that it is reasonable to choose the distance threshold by inspecting the clustering structures corresponding to the locally optimal mean silhouette scores. We note the importance of a manual inspection to decide on the trade-off between biologically interpretable number of clusters vs. cluster resolution. Our G2G package enables the user to inspect cluster diagnostics through distance vs. mean silhouette score plots, as well as the average alignment pattern of each cluster, to make a final decision.

#### Pathway overrepresentation analysis

We select the top *k* mismatching genes (i.e. genes with <=40% alignment similarity) from the list of genes ranked according to their alignment similarity percentage, to analyze their pathway overrepresentation. The identified clusters of genes are also analyzed. We use the GSEApy Enrichr^71,73,74^ wrapper against the *MSigDB_Hallmark_2020*^75^ and *KEGG_2021_Human* pathway genesets^75,76^. For all analyses, a 0.05 significance threshold of the adjusted P-value (with the default FDR correction method used by GSEApy) was applied.

#### Determining the best parameter setting

The G2Gframework has several key parameters: the number of interpolation time points with the corresponding pseudotime binning structure, the window_size of the Gaussian kernel used for interpolation, and the transition probabilities of the 5-state machine.

##### Pseudotime binning structure

For any number of interpolation time points, we can define an equispaced binning over the [0,1] range of the pseudotime axis (as also used by CellAlign^6^). A binning with low resolution (a low number of interpolation points) can be less representative of the dynamic process, while a binning with high resolution (a high number of interpolation points) can be too noisy or redundant. Therefore the optimal number of bins is a trade-off depending on the dataset at hand. We use the *optBinning*^77^ python package that implements a convex mixed-integer programming method, to heuristically decide the optimal number of bins in pseudotime range [0,1]. The *ContinuousOptimalBinning* module in *optBinning* can infer an optimal split structure given an array of continuous values. We obtain the optimal binning structure for reference and query pseudotime distributions separately. We then use the inferred optimal number of bins for equispaced binning of the pseudotime axis.

In all our datasets except for the T cell datasets, *optBinning* returned an equal number of optimal bins for both of the pseudotime distributions. For T cell datasets (i.e. pan fetal reference and ATO), we got 15 and 14 bins, respectively. For consistency, we selected the minimum of those two (i.e. 14 bins). We do not use the optimal splits returned by *optBinning* as it is an irregular binning structure which will not be consistent for alignment. As also used by both CellAlign^6^ and TrAGEDy^16^, equispaced binning is the safest and consistent choice.

##### Window_size

This parameter controls the effective neighborhood of contributions from all cells towards computing an interpolated mean and variance for each interpolation time point. CellAlign^6^ has found that *window_size* = 0.1 is the most effective for standard single-cell datasets (with a trade-off between noise and locality) and thus we use the same across all our experiments and analyses.

##### 5-state machine

As described previously, this machine has only 3free state transition probabilities: [*Pr*(*m*|*m*), *Pr*(*i*|*i*), *Pr*(*m*|*i*)]. We performed a grid search over the possible space of combinations, while fixing *Pr*(*m*|*m*)=0.99 to enforce that we have the highest probability of observing a continuous region of matches rather than a single point match. We found that [*Pr*(*i*|*i*) = 0.1, *Pr*(*m*|*i*) = 0.7] gives the lowest rate of false mismatch across the G2G alignments on our simulated benchmark dataset. This setting remained optimal when varying *Pr*(*m*|*m*) in the range [0.1, 1.0) (Supplementary Table 2). Therefore we use [*Pr*(*m*|*m*) = 0.99, *Pr*(*i*|*i*) = 0.1, *Pr*(*m*|*i*) = 0.7] as the G2G default setting.

### Benchmarking against CellAlign^6^ and TrAGEDy^16^ alignment

A DTW alignment for a given gene (i.e. gene-level alignment) or a set of genes (cell-level high-dimensional alignment) was generated using CellAlign’s *globalAlign* function, following the interpolation and scaling steps defined in the package documentation. The gene-level DTW alignments were clustered using CellAlign’s *pseudotimeClust* function. Similarly, TrAGEDy’s^16^ post-hoc-processed DTW alignments were generated using the functions in the published script provided by Laidlaw et al (2023), following documentation. An equal number of interpolation time points (i.e. 15 time points for our simulated dataset, as determined using *optBinning*^77^) were used for CellAlign^6^, TrAGEDy^16^ and G2G for fair comparison. To compare G2G alignments against their corresponding CellAlign^6^ and TrAGEDy^16^ alignments, we first converted them into their respective five-state alignment strings. As CellAlign only captures the match states [`M, W, V`], CellAlign mappings are always three-state strings. Note: CellAlign^6^ uses Euclidean distance, whereas TrAGEDy^16^ recommends Spearman correlation with the option to use Euclidean distance. Spearman correlation is mathematically undefined for a single gene dimension, and therefore cannot be run for gene-level alignment. Thus, we use TrAGEDy^16^ under Euclidean distance for both cell-level and gene-level alignment.

## Datasets

### Datasets for simulated experiments

#### Simulating pairwise datasets with different alignment patterns using Gaussian Processes

We modeled log-normalized expression of a gene *x* as a function *f* of time *t* using a Gaussian Process (GP), a stochastic process where any finite instantiation of it follows a multivariate Gaussian distribution. In other words, it is a distribution of functions, from which we can sample an *f*(*t*):

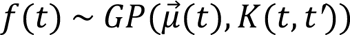

where, μ is the mean vector, and *K*(*t, t′*) is a kernel function which evaluates a covariance matrix covering every pair of finite time points where the *f*(t) is evaluated. The characteristics of this function are controlled by the class of the kernel being used (e.g. a Radial Basis Function (RBF) kernel for generating smooth, non branching functions; a change point kernel for generating branching functions). Therefore, a GP with an appropriate kernel is ideal to simulate different trajectory patterns in single-cell gene expression across pseudotime. Following the standard textbook and kernels discussed in literature^29,30^, we implemented a simulator for three different types of pairwise trajectory patterns: (1) *Matching*, (2) *Divergence*, and (3) *Convergence*, across a pseudotime range [0,1] with 300 total number of simulated cells for each trajectory.

#### Generating a *Matching* pair of reference and query gene trajectories

We used a GP with a constant *c* mean vector rμrr*c*’ (*c* ∈ [0.5,9.0] uniform random sampled) and RBF kernel *K* to first sample a function μ(*t*) that describes an average expression value for each time point. Next, we sampled two gene expression trajectories: **GEX**_*re*]_(*t*) and *GEX*_*query*_(*t*), from a GP where the mean is μ(*t*) and the covariance matrix is σ^2^*I* (σ ∈ [0.05,1.0] uniform random sampled, and *I*= identity matrix).

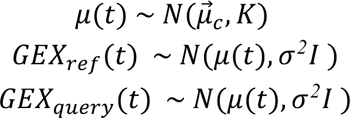

#### Generating a *Divergence* pair of reference and query gene trajectories

Here we use a Change Point (CP) kernel which imposes a bifurcation in the trajectory as it reaches an approximate time point *t*_*C*P_ (a.k.a. change point). The idea is to activate one covariance function before *t*_*C*P_ and another covariance function after *t*_*C*P_. We used the below CP kernel^29,30^ *K*_*C*P_:

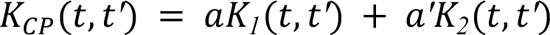

where,

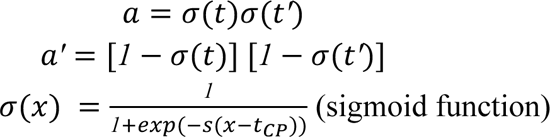

with **s** acting as a steepness parameter, deciding how steep the change point is. Penfold, et al (2018)^29^ defines a branching process by enforcing a zero kernel (*K_1_*) imposed before *t*_*C*P_and another suitable kernel (*K_2_*) afterwards. We used an RBF for *K_2_*. Following is the generative process starting with a base mean function μ(*t*) sampled from a separate GP with a constant *c* mean vector rμrr_*c*_’ (*c* ∈ [*0.5*,*9.0*] uniform randomly sampled), and an RBF kernel *K*.

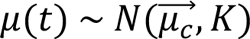

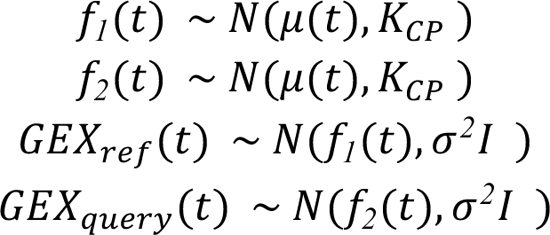

Next, two functions were sampled from a GP with base μ(*t*) and CP(t,t’), which were then used as mean vectors to generate *GEX*_*re*]_(*t*) and *GEX*_qu*ery*_(*t*) with covariance matrix σ*^2^I* (σ=0.3). This was run for [*t*_*C*P_=0.25, *t*_*C*P_=0.5, *t*_*C*P_= 0.75] to obtain 3 different groups of *Divergence* with varying bifurcation points. We use a constant σ which is not too low and not too high, and also apply a filtering criteria to ensure that the final dataset includes pairs of simple and clear divergence (a stable ground truth with no complex patterns). Pairs are filtered through basic heuristics such as the difference between mean expression before divergence and at the end terminals of reference and query.

Extended Data Fig. 1a-c shows that the branching effect may start approximately before the change point. Therefore, we expect the early non-divergent segment to continue at least until time point *i* < *t*_*C*P_ where we begin to see > 0.01 covariance in the change point kernel. Accordingly, given our approx_bifurcation_start_point = *i*, we expect the range of match lengths to fall between a lower limit = *n_tota*l_*pse*u*dotime*_*points* × *i* and upper limit = *n_tota*l_*pse*u*dotime*_*points* × *c*h*ange*_*point*. Equivalently, we expect mismatch lengths to fall between *n_tota*l_*pse*u*dotime*_*points* × (*1* − *i*) and upper limit = *n_tota*l_*pse*u*dotime*_*points* × (*1* − *c*h*ange*_*point*).

### Generating a *Convergence* pair of reference and query gene trajectories

For *Convergence*, we simply inverted the above generated divergent trajectory pairs, as the *Convergence* and *Divergence* patterns are complementary to each other.

### Simulating mismatches on real scRNA-seq data

We downloaded the mouse pancreas development dataset^31^ from the CellRank python package^78^. We subset the dataset to include only cells in the beta-cell lineage, using the annotations from the original authors (selecting cells labeled as “Ngn3 low EP”, “Ngn3 high EP”, “Fev+”, “Beta”), retaining 1845 cells. To select genes varying along beta-cell differentiation, we ran the CellRank pipeline (v1.5.1) to compute lineage drivers, following the package tutorials (https://cellrank.readthedocs.io/en/stable/auto_examples/estimators/compute_lineage_drivers.html) and selected genes significantly associated with differentiation potential to beta-cells (at 1% FDR we selected 769/2000 highly variable genes). For the pseudotime axis, we used the diffusion pseudotime estimated by the CellRank authors. To simulate trajectories for alignment, we divided the diffusion pseudotime (between 0 and 1) equally into 50 bins, assigned cells to the bins based on their estimated pseudotime and randomly split cells into query and reference datasets in each bin. To simulate deletions of *n* bins, we excluded query cells from the first *n* pseudotime bins (i.e. cells where the pseudotime value was less or equal to the upper margin of the *n*th bin). To simulate mismatches of *n* bins, we found the pseudotime bin with highest mean expression for the gene of interest in the query cells, and calculated mean (*max_mean*) and standard deviation (*max_std*) of expression of query cells for this bin; then, for each of the first *n* pseudotime bins, we substitute expression values of the query cells with a sample from a normal distribution with mean = *max_mean + max_std* and standard deviation = *max_std*. Pseudotime values for the query cells were min-max normalized to [0,1] after perturbation. We then ran the G2G alignment on each tested gene and calculated their alignment similarity (match calling) percentages.

### Datasets for benchmarking G2G

#### Dendritic cell stimulation dataset

The normalized single-cell expression datasets of PAM/LPS stimulation and their pseudotime estimates were downloaded from the CellAlign^6^ github repository and converted into *Anndata* objects. These contain two gene sets: ‘core antiviral module’ (99 genes) and ‘peaked inflammatory module’ (89 genes), pre-selected from the original publication^26^ and referred to as global and local modules, respectively by Alpert et al (2018)^6^. The datasets include 179 PAM-stimulated cells and 290 LPS-stimulated cells for which the pseudotime has been estimated by CellAlign^6^ authors using Diffusion-maps. CellAlign^6^ used 200 points for their PAM/LPS analysis as they observed adequate resolution. With *optBinning*^77^, we see that 14 bins are sufficient to represent both trajectories.

#### Simulated dataset containing trajectories with no shared process

This is a simulated, negative control dataset which was generated using the published script by Laidlaw, et al (2023) (in the TrAGEDy^16^ Git repository). Their script uses DynGen^79^, a single-cell data simulator for dynamic processes. The resulting negative dataset contains two trajectories simulated under two different gene regulatory networks and TF activity, ensuring that there is no shared process between them. The reference and query trajectories have 619 genes across 2000 and 1940 cells, respectively.

#### Dataset for Healthy versus IPF Case study

We downloaded the healthy and IPF dataset^27^ from GEO (GSE136831) and extracted our lineages of interest: AT2to AT1 cell differentiation for healthy; AT2 to Aberrant Basaloid Cell (ABC) differentiation for IPF (based on original author annotations). We identified a subset of 54 AT2 cells of low quality (with high percentage of ribosomal gene expression) which were filtered out prior to further analysis. Next we subsetted healthy and IPF cells to independently preprocess and estimate their pseudotime. This includes 3157 healthy cells (2655 AT2, 502 AT1) and 890 IPF cells (442 AT2, 448 ABCs). We first normalized their per-cell total transcript count to 10,000 followed by log1p normalization, and selected 4000 highly variable genes to estimate pseudotime of cells using Diffusion Pseudotime^38^ implemented in SCANPY^80^. The preprocessing steps were based on SCANPY documentation for trajectory inference, which included neighborhood graph construction, UMAP projection, denoising the graph by Diffusion Map construction, and running Diffusion Pseudotime estimation under default parameter settings. The root cell for each trajectory was chosen based on the expression score of known AT2 progenitor cell markers: AXIN2, FGFR2, ID2, FZD6, LRP5, LRP6^81,82^ (using scanpy.tl.score_genes function).

### Dataset preparation for *in vivo in vitro* T cell development comparison

#### Cell cultures for artificial thymic organoid (ATO) and single-cell RNA sequencing experiment

MS5 line transduced with human DLL4 was obtained from G. Crooks (UCLA) as a gift. The MS5-hDLL4 cells were cultured in DMEM (Gibco) with 10% FBS. Two iPSC lines were used in this study. Cell lines HPSI0114i-kolf_2 (Kolf) and HPSI0514i-fiaj_1 (Fiaj) were obtained from the Human Induced Pluripotent Stem Cell initiative (HipSci: www.hipsci.org) collection. All iPSC lines were cultured on vitronectin (diluted 1:25 in PBS; Gibco) coated plates, in TeSR-E8 media (Stemcell Technologies).

We followed the PSC-ATO protocol as previously described^43^. iPSC cells were harvested as a single-cell suspension and seeded (3×10^6^ cells per well) in GFR reduced Matrigel (Corning) - coated 6-well plates in X-VIVO 15 media (Lonza), supplemented with rhActivin A, rhBMP4, rhVEGF, rhFGF (all from R&D Systems), and ROCK inhibitor (Y27632; LKT Labs) on day −17, and only rhBMP4, rhVEGF and rhFGF on days −16 and −15. Cells were harvested 3.5 days later (day−14), and isolated by FACS for CD326^-^CD56^+^ (PE anti-human CD326 antibody, Biolegend, 324205; APC anti-human CD56 antibody, Biolegend, 318309) human embryonic mesodermal progenitors (hEMPs).

Isolated hEMPs were combined with MS5-hDLL4 at a ratio of 1:50. Two or three cell-dense droplets (5×10^5^ cells in 6 μl hematopoietic induction medium) were deposited on top of an insert in each well of a six-well plate. Hematopoietic induction medium composed of EGM2 (Lonza) supplemented with ROCK inhibitor and SB blocker (TGF-β receptor kinase inhibitor SB-431542; Abcam) was added into the wells outside the inserts so that the cells sat at the air-liquid interface. The organoids were then cultured in EGM2 with SB blocker for 7 days (days −14 to −7), before the addition of cytokines rhSCF, rhFLT3L, rhTPO (all from Peprotech) between days −6 to 0. These 2 weeks formed the hematopoietic induction phase. On day 1, media was changed again to RB27 (RPMI supplemented with B27 (Gibco), ascorbic acid (Sigma-Aldrich), penicillin/streptomycin (Sigma-Aldrich) and glutamax (Thermo Fisher Scientific)) with rhSCF, rhFLT3L and rhIL7. The organoids can be maintained in culture for 7 more weeks in this medium.

For dissociation of organoids on day −7, they were removed from culture insert and incubated in digestion buffer, which consisted of collagenase type IV solution (StemCell Technologies) supplemented with 0.88mg/ml collagenase/dispase (Roche) and 50U DNase I (Sigma), for 20 minutes at 37°C. Vigorous pipetting was performed in the middle of the incubation and at the end. After complete disaggregation, single cell suspension was prepared by passing through a 50-μm strainer.

For dissociation of organoids from day 0 onwards, a cell scraper was used to detach ATOs from cell culture insert membranes and detached ATOs were then submerged in cold flow buffer (PBS (Gibco) containing 2% (v/v) fetal bovine serum (FBS; Gibco) and 2 mM EDTA (Invitrogen)). Culture inserts were washed and detached ATOs were pipetted up and down to form single-cell suspension before passing through a 50-μm strainer.

Cells were then stained with designed panels of antibodies and analyzed by flow cytometry. FACS was performed at the same time and live human DAPI^-^anti-mouse CD29^-^ (APC/Cy7 anti-mouse CD29 antibody, Biolegend, 102225) cells were sorted for day −7, day 0 and week 3 samples, and live (DAPI^-^) cells were sorted for week 5 and week 7 samples before loading onto each channel of the Chromium chip from Chromium single cell V(D)J kit (10X Genomics). The metadata for all the ATO samples can be found in **Supplementary Table 8.** For the day −14 sample, some sorted (both hEMP and the rest of DAPI^-^ fraction) and unsorted cells were stained with hashtag antibodies (TotalSeq-C antibodies from Biolegend, see **Supplementary Table 9**, following 10X cell surface protein labeling protocol) before being mixed together with some mouse stromal cells (MS5-hDLL4) for 10X loading. For week 1 sample, hashtag antibodies were added in at the same time as the FACS antibodies i.e. before sorting.

Single-cell cDNA synthesis, amplification and gene expression (GEX) and cell surface protein (CITE-seq) libraries were generated following the manufacturer’s instructions. Sequencing was performed on the Illumina Novaseq 6000 system. The gene expression libraries were sequenced at a target depth of 50,000 reads per cell using the following parameters: Read1: 26 cycles, i7: 8 cycles, i5: 0 cycles; Read2: 91 cycles to generate 75-bp paired-end reads.

### ATO data preprocessing and annotation

Raw scRNA-seq reads were mapped with cellranger 3.0.2 with combined human reference of GRCh38.93 and mouse reference of mm10-3.1.0. Low quality cells were filtered out (minimum number of reads = 2000, minimum number of genes = 500, maximum number of genes = 7000, Scrublet^83^ (v0.2.3) doublet detection score < 0.15, mitochondrial reads fraction < 0.2). Cells where the percentage of counts from human genes was < 90% were considered as mouse cells and excluded from downstream analysis. Cells were assigned to different cell lines (Kolf, Fiaj) using genotype prediction with Souporcell (v2.4.0)^84^. The mapping outputs of the eight samples were merged, with the sample ID prepended to the barcode IDs in both the BAM and barcodes.tsv to prevent erroneous cross-sample barcode overlap. Souporcell was run with `--skip_remap True –-K 2` and the common variants file based on common (>= 2% population allele frequency) SNPs from 1000 genomes data, as distributed in the tool’s repository. We selected 2 clusters due to the already known 2 cell lines. Next the data went through the standard pipeline of filtering out genes (cell cycle^85^ genes, genes detected in less than 3 cells), and normalizing the per cell total count to 10,000 followed by log1p transformation and scaling to zero mean and unit variance (with max_value = 10 to clip after scaling), using SCANPY^80^. The final dataset had 31,483 ATO cells with 23,526 genes which were input to CellTypist^44^ (for prediction using pre-trained logistic regression classifier – *Pan_Fetal_Human* model under majority voting). We then obtained a Uniform Manifold Approximation and Projection (UMAP) embedding for this dataset based on its scVI^45^ batch corrected embedding (v0.14.5 with 10 latent dimensions (default), 2 hidden layers, 128 nodes per hidden layer (default), and 0.2 dropout rate for the neural network), and subsetted cells to non-hematopoietic lineage, T/ILC/NK lineage, and other hematopoietic lineage cells (**Extended Data Fig. 8**) using their Leiden clustering. By default, scVI models gene counts with zero-inflated negative binomial distribution, defines a normal latent distribution, and handles batch effects. For each lineage, scVI latent dimensions and UMAP embedding were re-computed and cell types were annotated (low-level annotations in **Extended Data** Fig. 6b, with more refined annotations in **Extended Data** Fig. 7a) by inspecting both the CellTypist results (**Extended Data** Fig. 7b) and marker gene expression (**Extended Data** Fig. 8).

### Joint Embedding of reference and organoid in preparation for pseudotime estimation

We downloaded the annotated human fetal atlas dataset from https://developmental.cellatlas.io/fetal-immune and extracted the cell types (79,535 cells in total) representing the T cell developmental trajectory starting from progenitor cells towards type 1 innate T cells (T1 dataset), including Cycling MPP, HSC_MPP, LMPP_MLP, DN(early) T, DN(P) T, DN(Q) T, DP(P) T, DP(Q) T, ABT(entry) and type 1 innate T cells. We then compiled a reduced representation (20,384 cells) that preserve their underlying cell type composition. This was done by random subsampling from each cell type (with minimum sample size = 500 cells, aiming ∼20,000 total number of cells) based on their originally published annotations. Such stratified-sampling approach is practical for dealing with massive single-cell datasets to reduce computational resource requirements. Separately, we extracted the cell types from the ATO dataset (19,013 cells) representing the trajectory starting from iPSCs towards SP T cells, including iPSC, primitive streak, mesodermal progenitor, endothelium, HSC_MPP, HSC_MPP/LMPP_MLP/DC2, DN(early) T, DN T, DP(P) T, DP(Q) T, ABT(entry), SP T cells.

Then the T1 and ATO datasets were merged and preprocessed together by filtering out cells with more than 8% total mitochondrial UMI, cell cycle genes^85^ and genes expressed in less than 3 cells (`min_cells = 3`). Next, the highly variable genes were selected after per cell count normalization to 10,000 reads per cell and log1p normalization. The T1 pan fetal reference had 33 batches (due to different 10X chemistry 3’ *versus* 5’ and different donors), while the ATO had only 2 batches (due to 2 cell lines, Kolf and Fiaj). This merged dataset was then input to constructing a joint latent embedding using the scVI (v0.14.5) variational autoencoder implementation^45^ (with 10 latent dimensions (default), 2 hidden layers, 128 nodes per hidden layer (default) and 0.2 dropout rate for the neural network). The joint embedding was then taken to build the cell neighborhood graph and UMAP embedding using SCANPY^80^. The final T1 and ATO datasets comprise 20,327 cells and 17,176 cells respectively. 18,436 cells of T1 and 10,089 cells of ATO belong to T cell lineage, i.e., DN T onwards.

We followed a similar preprocessing for the pan fetal reference representing the trajectory towards CD8+T (CD8 dataset) (including Cycling MPP, HSC_MPP, LMPP_MLP, DN(early) T, DN(P) T, DN(Q) T, DP(P) T, DP(Q) T, ABT(entry) and CD8+T cells). The initially extracted CD8 subset (83,177 cells) was reduced to 20,412 cells, which was then merged with the 19,013 ATO cells and subjected to the same filtering and normalization as for the T1+ATO merge prior to scVI integration. The final CD8 dataset comprises 20,324 cells of which, 18,490 cells are DN T onwards.

### Pseudotime estimation using the Gaussian Process Latent Variable Model

The differentiation pseudotime was estimated separately for T1 reference, CD8 reference and ATO by employing the Gaussian Process Latent Variable Model (GPLVM)^46,49^. GPLVM is a probabilistic nonlinear dimensionality reduction method which models observed gene expression as a function *f*(*X*) of a set of latent covariates X. It enables us to incorporate Gaussian time priors when estimating pseudotime as a latent dimension. We used the Pyro^86^ GPLVM implementation (Pyro v1.8.0) with Sparse Gaussian Process inference (32 inducing points) and Radial Basis Function kernel to obtain a 2D latent embedding, where the first latent dimension corresponds to pseudotime and the second latent dimension corresponds to a second level of latent effects (e.g. batch). The first dimension was assigned a Gaussian prior with cell capture times as the mean. The second latent dimension was zero initialized to allow for a second level of latent effects (e.g. batch). The model used Adam optimizer to infer the optimal latent embedding with 2000 iterations (where the loss curve reasonably converged).

For the ATO, the GPLVM was initialized with cell capture days as the prior. Since there was no temporal data present in the pan fetal reference, we first approximated time prior for each reference cell as the weighted average of their k-nearest organoid neighborhood (kNN) capture time. A k=3 organoid neighborhood for a reference cell was obtained using the cKDTree based search method implemented in BBKNN^87^ on their scVI based UMAP embedding. Contribution of each organoid neighbor was weighted according to its distance. (kNN distance vector was softmax transformed, and the normalized reciprocal of each distance was taken as the associated weight, enforcing less contribution from distant neighbors towards the weighted average). This approximation may introduce outliers due to the spatial arrangement of different cell types in the UMAP. Thus, we leveraged the known cell-type annotations to refine the approximation by assigning each reference cell with the average approximated capture time of its cell type. These approximated capture times were scaled to be in [0,1] range and input as the mean prior to the previously described GPLVM. For T1 and CD8 GPLVMs, the input gene space was 2608 genes and 2616 genes respectively (same as at scVI integration). To ensure no outliers, the GPLVM estimated pseudotime was further refined by correcting outliers of each cell type using the cell-type specific average of estimated pseudotime. Outliers were selected based on the Interquartile Range (IQR) rule (i.e. 1.5 times IQR below the first quartile and above the third quartile of the cell-type specific pseudotime distribution).

### Genes2Genes alignment

For the complete T1 vs ATO comparison using G2G, the total common gene space of 20,240 genes was considered upon filtering genes with less than 3 cells expressed, 10,000 total count per cell normalization, and log1p normalization. For the DN T onwards comparison, there were 17,718 genes for T1 vs. ATO, and 20,183 genes for CD8 vs. ATO. All pseudotime estimates were min-max normalized to ensure [0,1] range. These total gene spaces were subsetted to include only the transcription factors^52^ (1371 TFs) and relevant signaling pathways focused in this work. G2G alignment was performed using 14 equispaced pseudotime points (the number of optimal bins was determined using *optBinning*^77^).

### ATO TNFɑ validation experiment

For the TNFɑ validation experiment, we followed the PSC-ATO protocol described previously using Kolf iPSC line. One plate (12 organoids) was set for each condition, with 4 conditions which were 1) control i.e., no TNFɑ addition, 2) 1 ng/ml TNFɑ (final concentration), 3) 5 ng/ml TNFɑ, and 4) 25 ng/ml TNFɑ. TNFɑ was added into the media between week 6 to 7. Organoids were harvested at week 7, stained with hashtag antibodies (TotalSeq-C antibodies from Biolegend) and a designed panel of FACS antibodies, and sorted for CD45^+^ cells before 10X loading. Library construction, sequencing and data preprocessing was the same as the WT ATO organoids (see sections above).

### Assessing T cells from ATO before and ATO after TNFɑ treatment

The ATO with TNFɑ treatment (ATO_TNFɑ_) resulted in 123 good quality T cells (data preprocessing and filtering as above, annotated by CellTypist^44^ as type 1 innate T cells) where majority (∼94% cells) are 1ng/ml TNFɑ treated (as opposed to wild-type, 5ng/ml, and 25ng/ml). Thus we performed the analysis using T cells treated with 1ng/ml TNFɑ (i.e. 116 cells). As our specific goal was to assess only the maturity level of type 1 innate T cells, this excludes the 3 cells annotated by CellTypist^44^.

The wild type ATO without TNFɑ treatment has 6558SP T cells and the pan fetal reference (Ref) has 1413 type 1 innate T cells. To check whether there is an improvement in the maturity of resultant T cells in ATO_TNFɑ_, we evaluated and compared the degree of similarity between ATO_TNFɑ_ and the reference against that between wild-type ATO and the reference. This was done via two directions: a cell-level assessment and a gene-level assessment. For cell-level assessment, we constructed an scVI^45^ latent embedding by integrating the reference, ATO and ATO_TNFɑ_ together, following the same preprocessing steps as done before for the wild-type ATO and the reference. We then computed the euclidean distance between the mean latent dimensions of in vitro SP T cells and the in vivo type 1 innate T cells. For gene-level assessment, we computed the MML distance of gene expression distributions by first identifying significantly distant genes across all the TFs, as well as across several curated pathway gene subsets from KEGG and MSigDB that are associated with TNFɑ signaling (See Supplementary Note), before and after TNFɑ treatment. We next tested if the difference between those distance distributions is significant or not using the Mann-Whitney U test. The following section describes our approach to identifying genes with a statistically significant distance.

### Estimating an empirical null distribution of MML distances to assess significance of a distance

We expect that there is no significant difference in gene expression between any two random subsets of cells belonging to the same cell type in the same system. Thus, we generated such random pairs of cell subsets within our pan fetal reference (type 1 innate T cell) dataset as well as within the ATO (Single Positive T cell) dataset across all 1371 TFs, to construct an empirical null distribution for MML distance. This empirical distribution enables us to compute a p-value for any given MML distance *x*. The p-value indicates how likely *x* is to occur randomly by chance.

For each TF, we performed uniform random sampling of a subset pair without replacement (using *numpy.random.choice*) for 50 iterations, resulting in 137,100 total number of pairs. Each subset contains 50 cells. Then, for each pair, we computed the MML distance. All computed MML distances together act as a null distribution which we then used to evaluate whether some MML distance measure between two expression distributions is significant or not.

The resultant empirical null distribution of MML distances is highly left skewed. We constructed its empirical cumulative density function (CDF) using *distributions.empirical_distribution.ECDF* function in the *statsmodels* package to test significance in distances under no assumption about the family of distribution. The p-values were adjusted for multiple testing using *statsmodels.stats.multitest.fdrcorrection* (benjamini– hochberg method) correcting for false discovery rate, before identifying significantly distant genes at a given level of significance.

### Testing statistical significance of the change in gene expression distances to reference from wild-type ATO versus ATO_TNFɑ_

Given the MML distances of the genes which are significantly different: (1) between the reference and wild-type ATO (sample 1), and (2) between the reference and ATO_TNFɑ_ (sample 2), we tested whether the average gene expression distance dropped after TNFα treatment. To test the significance of the change in distance, we performed the non-parametric Mann-Whitney U test implemented in *scipy.stats.mannwhitneyu*. Given two independent samples of scores, the null hypothesis of the Mann-Whitney U test assumes that their underlying distributions are the same, while the alternative hypothesis assumes that the distribution underlying the first sample is stochastically greater than the distribution underlying the second sample.

### Testing the closeness of SP T cells in the ATO to the reference type 1 innate T cells before and after TNFα treatment

We compared the 6558 SP T cells in the wild-type ATO and the 115 T cells in the TNFα treated ATO (ATO_TNFɑ_), against the 1413 type 1 innate T cells from the pan fetal reference. This was done in terms of all the 1371 human TFs, as well as the 196 genes in the TNFα signaling via NF-ϰB pathway (defined and curated by MSigDb).

Given each geneset, we first computed the MML distances of the genes between: (1) reference and ATO, and (2) reference and ATO_TNFɑ_. We then computed a p-value for each MML distance under the empirical cumulative density function (CDF) estimated by computing MML distances between randomly sampled homogeneous subsets across the reference and ATO cells. This allowed us to identify the gene set which is significantly different between reference and ATO (diffset_UNTREATED_), and the gene set which is significantly different between reference and ATO_TNFɑ_ (diffset_TREATED_). Next we compared the average distance of genes in diffset_UNTREATED_ to the average distance of genes in diffset_TREATED_, while testing the significance of the change in the distances using Mann-Whitney U test.

### Software packages used in the work

Genes2Genes framework and all analysis related code (including plot generation) were implemented using the standard Python libraries (Numpy, Pandas, Seaborn, scikit-learn, SciPy, statmodels), GPyTorch, GSEApy. Illustrations were made using Adobe Illustrator 2023 and BioRender.

## Code and data availability

**Genes2Genes** is implemented as an open-source package in Python 3 (v3.8) with tutorial available at: https://github.com/Teichlab/Genes2Genes. Code and data used to generate figures and perform analyses in the manuscript are available at: https://github.com/Teichlab/G2G_notebooks and https://drive.google.com/drive/folders/15LKmo3yRB-cR8Aq3aE59Taq2r0KtJSdX?usp=sharing. All generated alignments are available as Supplementary Data. Raw sequencing data for newly generated sequencing libraries have been deposited in ArrayExpress (accession number E-MTAB-12720).

## Acknowledgements

This paper is dedicated to the memory of our dear friend and colleague Dr. Daniele Muraro who contributed to this work. We thank the Crooks lab (A. Montel-Hagen, S. Lopez, and G. Crooks from University of California, Los Angeles) for their kind help in setting up the ATO experiments; N Huang, L Dratva, R Lindeboom, R Elmentaite, A Maartens from the Wellcome Sanger Institute, CH Ek from University of Cambridge, Z Miao from Guangzhou Laboratory, Y Chen and T Wang from London School of Economics and Political Science for their helpful discussions; and BioRender.com for graphical illustrations. We gratefully acknowledge the Sanger Flow Cytometry Facility and Sanger Core Sequencing pipeline for support with sample processing and sequencing library preparation. We acknowledge Wellcome Sanger Institute as the source of HPSI0114i-kolf_2 and HPSI0514i-fiaj_1 human induced pluripotent cell lines which were generated under the Human Induced Pluripotent Stem Cell Initiative funded by a grant from the Wellcome Trust and Medical Research Council, supported by the Wellcome Trust (WT098051) and the NIHR/Wellcome Trust Clinical Research Facility, and acknowledge Life Science Technologies Corporation as the provider of Cytotune. This publication is part of the Human Cell Atlas - www.humancellatlas.org/publications.

## Funding

D.S. is supported by the Marie Curie Individual Fellowship; This project has received funding from the European Union’s Horizon 2020research and innovation programme under the Marie Skłodowska-Curie grant agreement No: 101026506. C.S. is supported by a Wellcome Trust Ph.D. Fellowship for Clinicians. S.A.T. is funded by Wellcome (WT206194), the ERC Consolidator Grant ThDEFINE (646794). This work was also supported by a grant from the Wellcome Sanger Institute’s Translation Committee Fund.

## Author contributions

D.S., C.S. and S.A.T. conceived the initial project. D.S. and C.S. set up and directed the study. D.S., C.S., A.C, E.D. and K.P. performed bioinformatic analyses. D.S. designed and developed the software. C.S., A.S.S., W.L., and J.P. performed cell culture experiments. D.M., E.D., A.J.O., K.B.M., B.D. and S.A.T. provided intellectual input. S.A.T. acquired funding. D.S., C.S., and A.C. wrote the manuscript. All authors read and/or edited the manuscript.

## Competing interests

In the past three years, S.A.T. has received remuneration for Scientific Advisory Board Membership from Sanofi, GlaxoSmithKline, Foresite Labs and Qiagen. S.A.T. is a co-founder and holds equity in Transition Bio.

## Extended Data Figures

**Extended Data Fig. 1.**
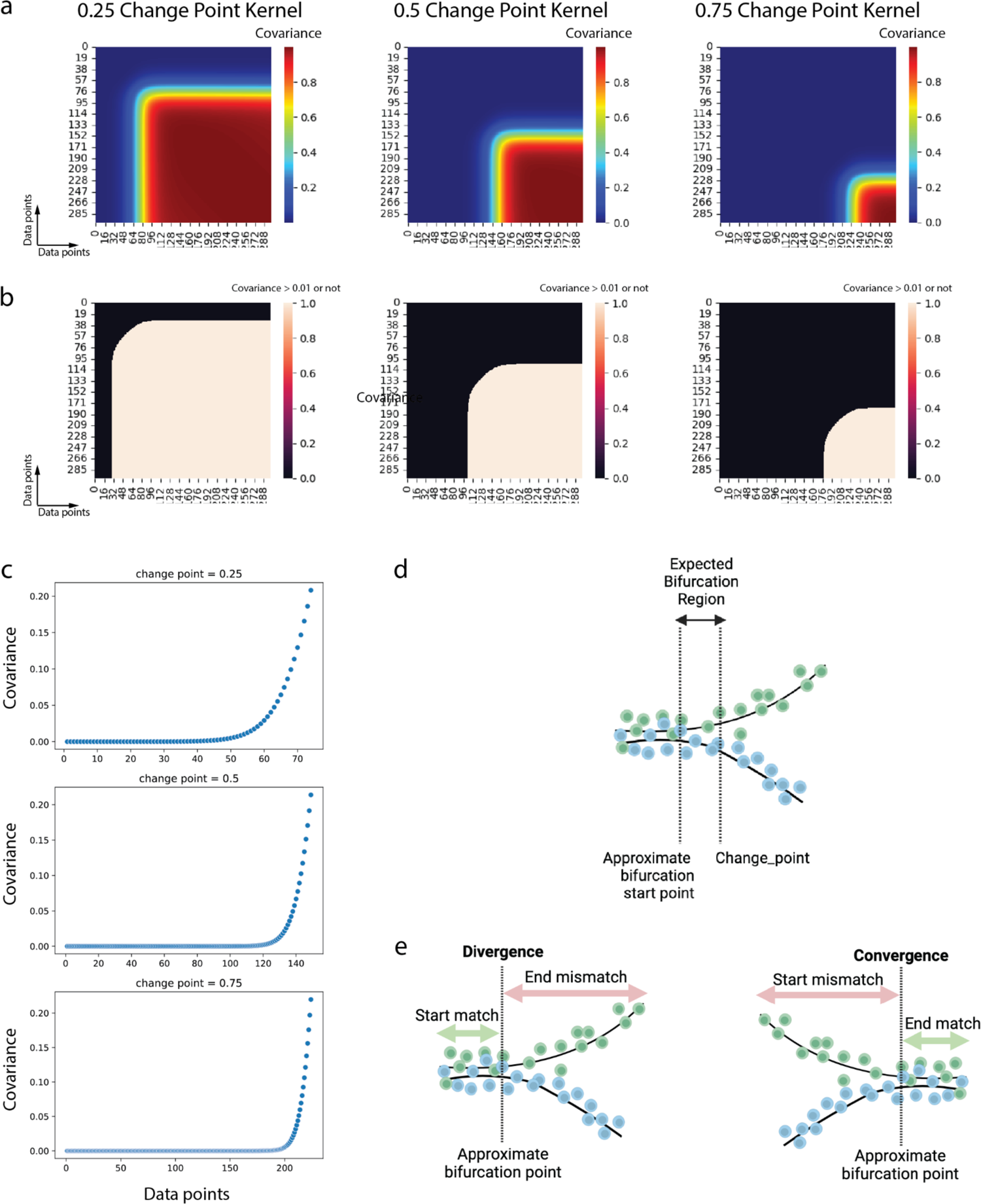
Simulating the bifurcation of reference and query trajectories using change point kernels. **a**, Change point kernel heatmaps for each approximate bifurcation point (change point) ∈ [0.25,0.5,0.75]. **b**, The same change point kernels binarized based on the 0.01 covariance threshold (top), **c**, The average covariance plotted for each *i* × *i* sub square matrix from *i* = 0 to *i* = change point, showing that the branching effect can approximately start before the specified change point. **d**, Expected bifurcation region is taken from the point where we begin to see > 0.01 covariance in the change point kernel, until the particular change point. **e**, Illustration of the main regions of match and mismatch expected in trajectory alignment under *Divergence* class (left) and *Convergence* class (right). A *Divergence* alignment is described by a start-match region followed by an end-mismatch region, whereas a *Convergence* alignment is described by a start-mismatch region followed by an end-match region.

**Extended Data Fig. 2.**
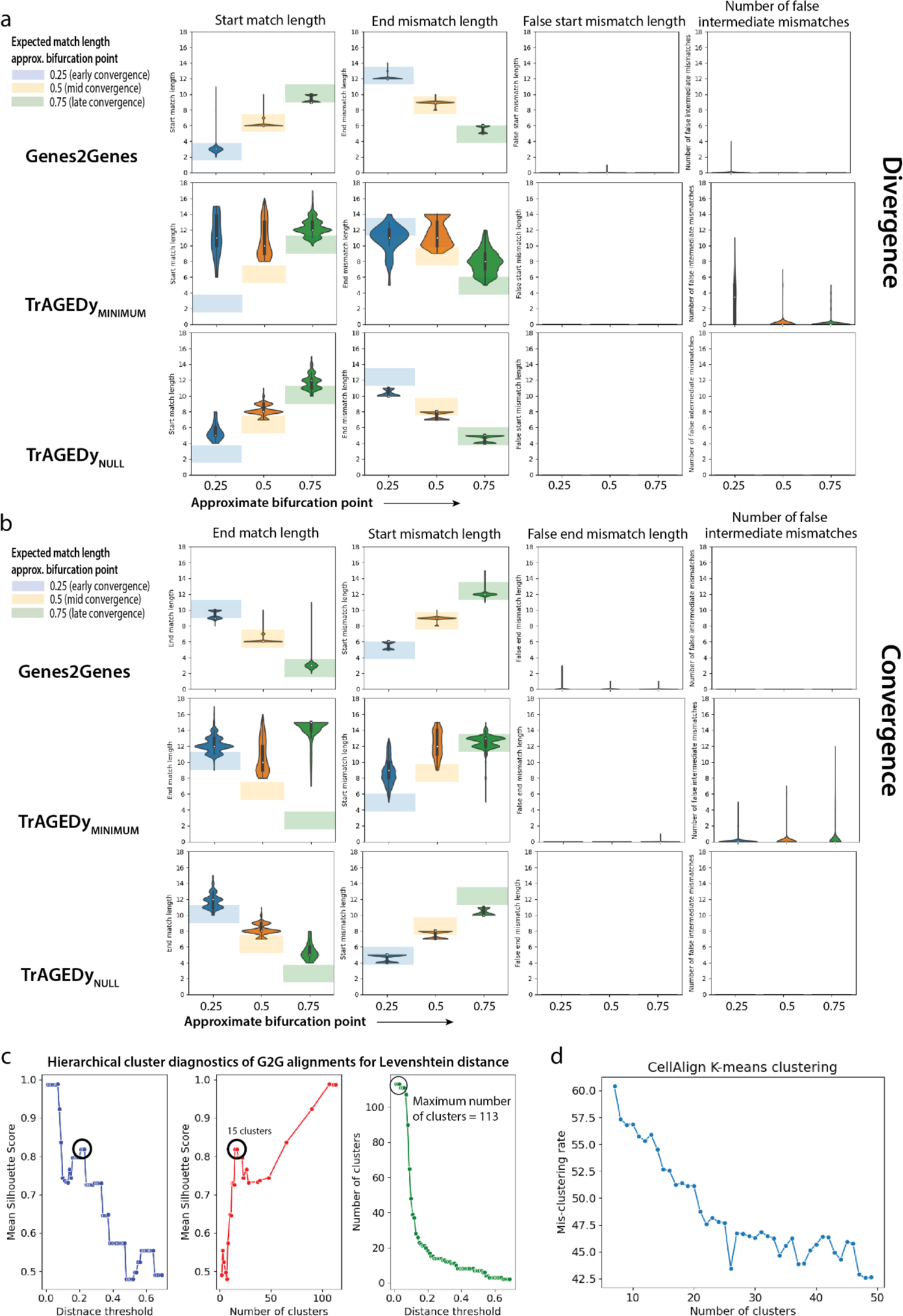
Simulation Data Experiment 1. **a**, Distributions of start-match lengths (following a false mismatch if there is any), end-mismatch lengths (prior to a false match if there is any), start-mismatch lengths (number of false mismatches starting from time point 0), and the number of intermediate false mismatches within the match regions, in the 1500 *Divergence* alignments across the three bifurcation subgroups (i.e., under approximate bifurcation point ∈ [0.25,0.5,0.75]), generated by Genes2Genes, TrAGEDy_MINIMUM_, and TrAGEDy_NULL_. 15 equispaced time points on pseudotime [0,1] were used for distribution interpolation and alignment. **b**, Similar statistics as in **a**, reported for the 1500 *Convergence* alignments across the three bifurcation subgroups, generated by Genes2Genes, TrAGEDy_MINIMUM_, and TrAGEDy_NULL_. Distributions of end-match lengths (prior to a false mismatch if there is any), start-mismatch lengths (following a false match if there is any), end-mismatch lengths (number of false mismatches until time point 1), and the number of intermediate false mismatches within the match regions. **c**, Cluster diagnostic plots for the hierarchical agglomerative clustering of the 3500 alignments across all pattern classes (including 500 *Matching* alignments, 1500 *Divergence* alignments, 1500 *Convergence* alignments), in terms of the mean Silhouette score when varying the Levenshtein distance threshold (or the number of clusters). The highest number of clusters represent the number of all unique 5-state alignment strings (i.e. 113 strings). Bold highlighted circles mark the local optimal mean Silhouette score which gives 15 optimal clusters for the genes at 0.22 distance threshold. **d**, Mis-clustering rates of the CellAlign k-means clustering outputs for all 3500 alignments, versus the number of clusters (k) ranging in [7,50].

**Extended Data Fig. 3.**
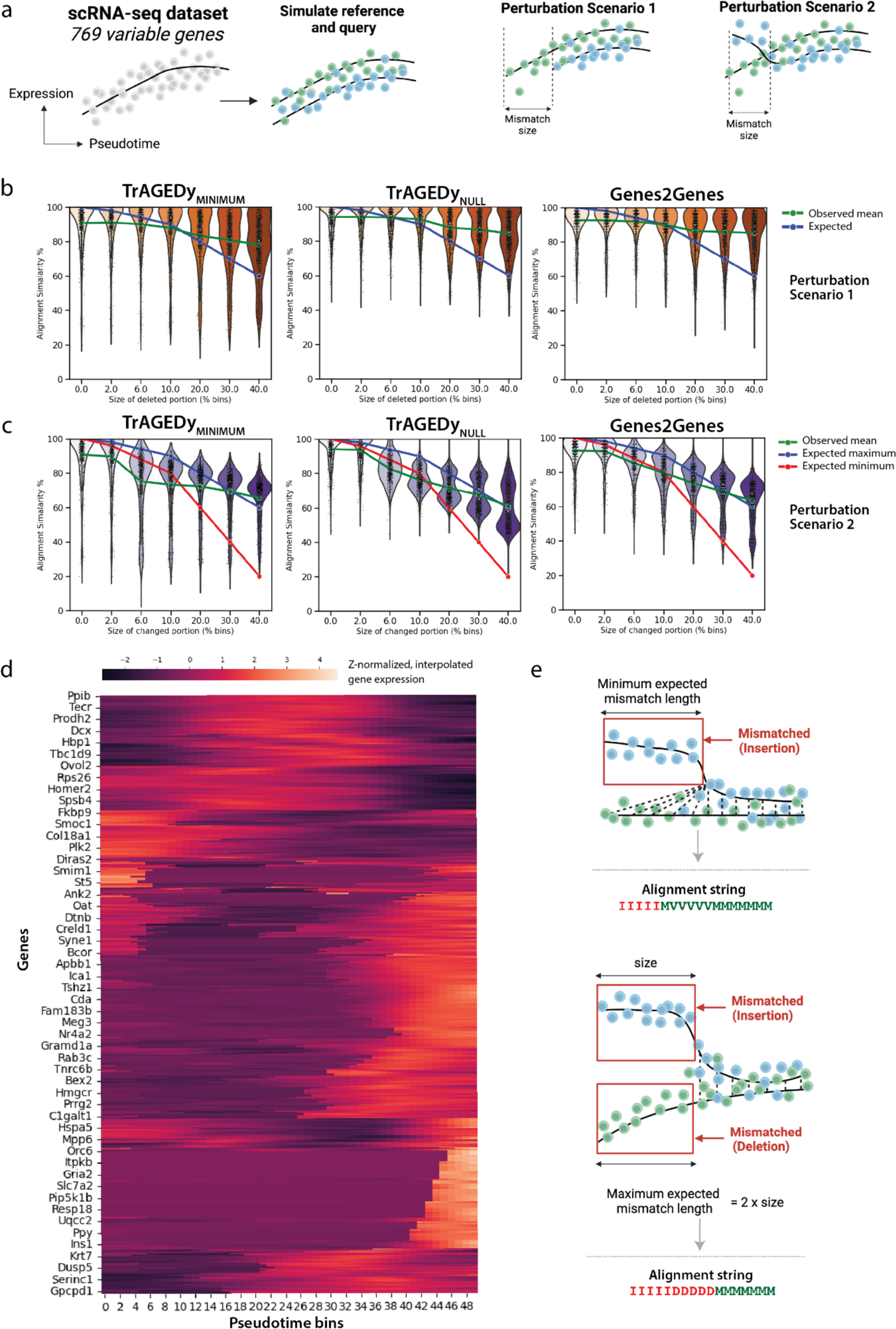
Simulation Data Experiment 2. a, Experiment 2 uses 769 genes in the mouse pancreas development dataset (Beta lineage) scRNA-seq dataset ^31^ to generate perturbed pairs of alignment from the expected Matching alignments. Perturbation scenario 1 deletes the start region from the reference trajectory, whereas perturbation scenario 2 changes the start region from the reference trajectory. b, The alignment similarity distributions for varying sizes (percentage of 50 pseudotime bins) of perturbation under perturbation scenario 1 (top) and perturbation scenario 2 (bottom), resulted from gene-level alignment using TrAGEDy_MINIMUM,_ TrAGEDy_NULL_, and Genes2Genes. Each point in the violin plot represents a gene (total number of genes n = 769). In each plot, the observed average alignment similarity across different perturbation sizes is shown by the green line. For perturbation scenario 1 (top), the blue line shows the expected alignment similarity across different perturbation sizes. For perturbation scenario 2 (bottom), there are two expected lines: maximum (in blue) and minimum (in red). The maximum mismatch length is expected when both reference and query time points form insertions and deletions, making the maximum expected length size*2. The minimum mismatch length is expected when only the changed reference time points are mismatched as insertions, while the corresponding query time points are matched to the non-perturbed reference time points (illustrated in e). d, Overall smoothened (interpolated) and z-normalized mean gene expression along the pseudotime (equally divided into 50 bins) for all genes in the dataset. e, Example illustrations of the two types of trajectory alignment that gives the minimum expected mismatch length and the maximum expected mismatch length under the perturbation scenario 2, where a start portion of a particular size in the query trajectory (in blue) is changed with respect to the reference trajectory (in green).

**Extended Data Fig. 4.**
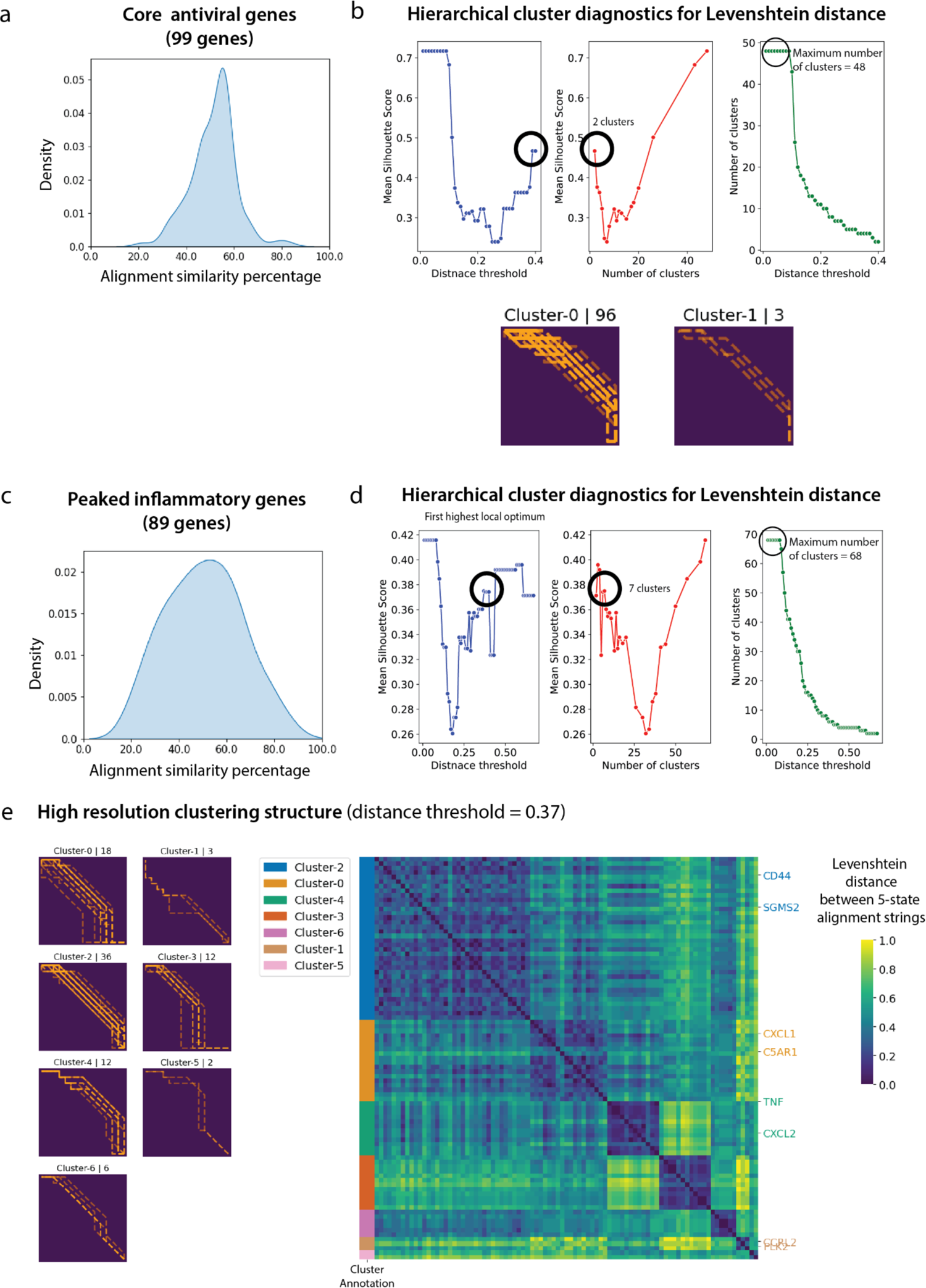
The PAM vs. LPS alignment using Genes2Genes. **a**, Density plot of the alignment similarity distribution (i.e. distribution of the percentage of matches/warps across all the alignment outputs) for the 99 genes in the ‘core antiviral module’. **b**, Top: Cluster diagnostic plots for the hierarchical agglomerative clustering of those 99 gene alignments in terms of the mean Silhouette score when varying the Levenshtein distance threshold (or the number of clusters). The highest number of clusters represent the number of all unique 5-state alignment strings (i.e. 48 strings). Bold highlighted circles mark the local optimal mean Silhouette score which gives two optimal clusters for the genes at 0.4 distance threshold. Bottom: Each plot titled by “Cluster-x | n” is the pairwise matrix of reference time points (columns) and query time points (rows), visualizing alignment paths for the total of n genes in cluster x. **c**, Density plot of the alignment similarity distribution (i.e. distribution of the percentage of matches/warps across all the alignment outputs) for the 89 genes in the ‘peaked inflammatory module’. **d**, Cluster diagnostic plots for the hierarchical agglomerative clustering of those 89 gene alignments, reported similarly to **b.** The identified optimal clustering structure has 7 clusters (at distance threshold=0.37 corresponding to the local optimal mean Silhouette score, highlighted by bold circles). **e**, The clustermap of the pairwise Levenshtein distance matrix of all 89 gene alignments, which illustrates the identified 7 clusters.

**Extended Data Fig. 5.**
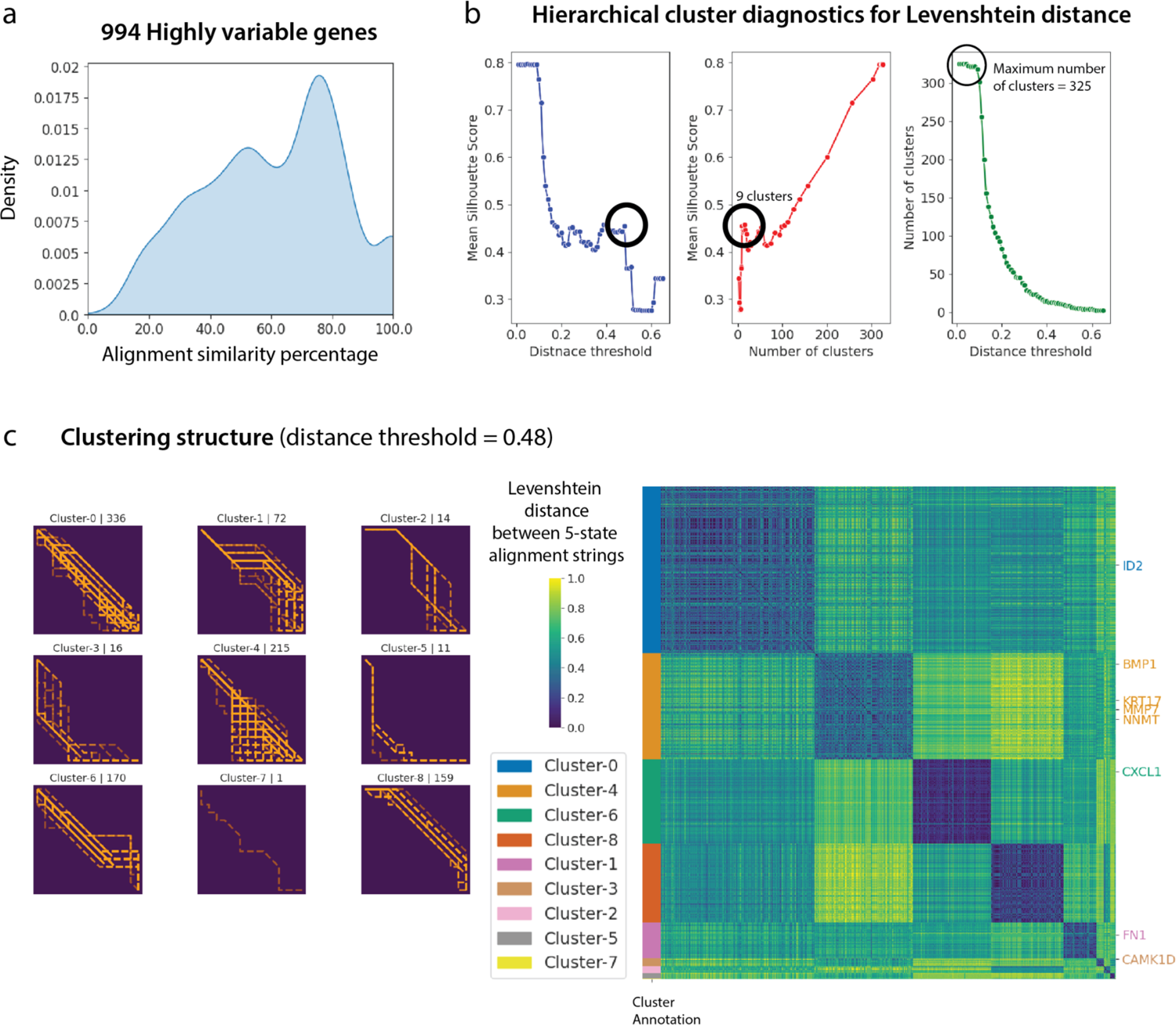
The healthy versus Idiopathic Pulmonary Fibrosis (IPF) disease case study alignment clustering outputs. **a**, Density plot of the alignment similarity distribution (i.e. distribution of the percentage of matches/warps across all the alignment outputs) for the 994 highly variable genes in the dataset. **b**, Cluster diagnostic plots for the hierarchical agglomerative clustering of those 994 gene alignments in terms of the mean Silhouette score when varying the Levenshtein distance threshold (or the number of clusters). The highest number of clusters represent the number of all unique 5-state alignment strings (i.e. 325 strings). Bold highlighted circles mark the local optimal mean Silhouette score which gives nine optimal clusters for the genes at 0.48 distance threshold. **c**, The identified clustering structure. Left: Each plot titled by “Cluster-x | n” is the pairwise matrix of reference and query time points, visualizing alignment paths for all the genes (one alignment per gene and a total of n genes in the cluster) in a cluster x. Right: The clustermap of the pairwise Levenshtein distance matrix of all 994 gene alignments.

**Extended Data Fig. 6.**
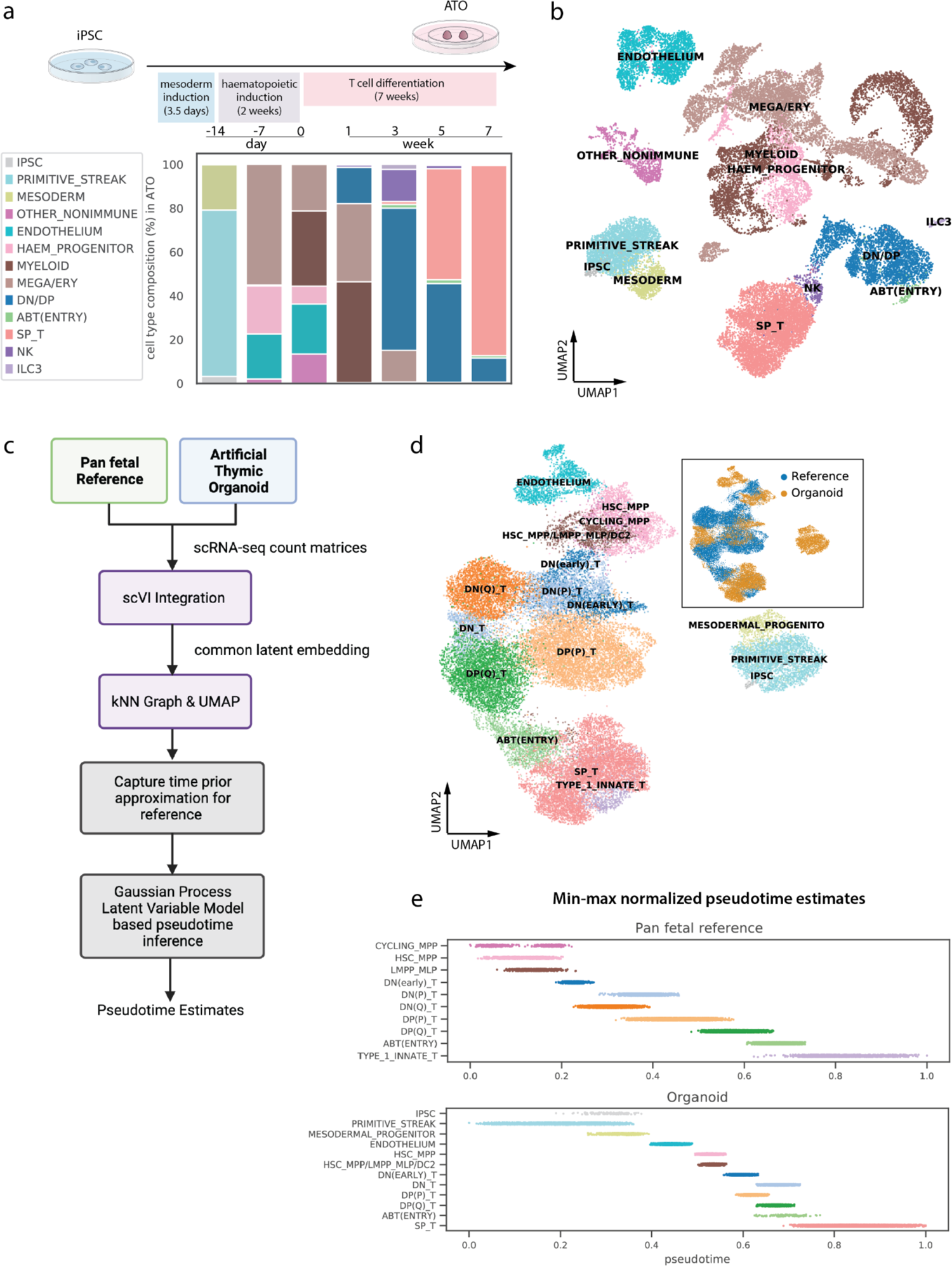
in vivo, in vitro human T cell development data integration and pseudotime inference. **a**, Top: schematic showing the experimental set-up of T cell differentiation from iPSCs in ATOs. Bottom: barplot of cell type composition in ATO at different time points during differentiation. **b**, UMAP visualization of different cell types in the ATO dataset (low-level annotation, number of cells n = 31,483), with more refined annotation in **Extended Data Fig. 7a**. **c**, Workflow of integrating *in vitro* (i.e. ATO) and *in vivo* (i.e. pan fetal reference from Suo et al. 2022^42^) human T cell development data and pseudotime inference using GPLVM. **d**, Main: UMAP visualization of integrated *in vivo* and *in vitro* human T cell development data, colored by the cell types. Right insert: the same UMAP visualization colored by the data source. **e**, Stripplot of the inferred pseudotime (*x*-axis) against different cell types (*y*-axis), colored by the cell types, of *in vivo* pan fetal reference data (top) and *in vitro* organoid data (bottom).

**Extended Data Fig. 7.**
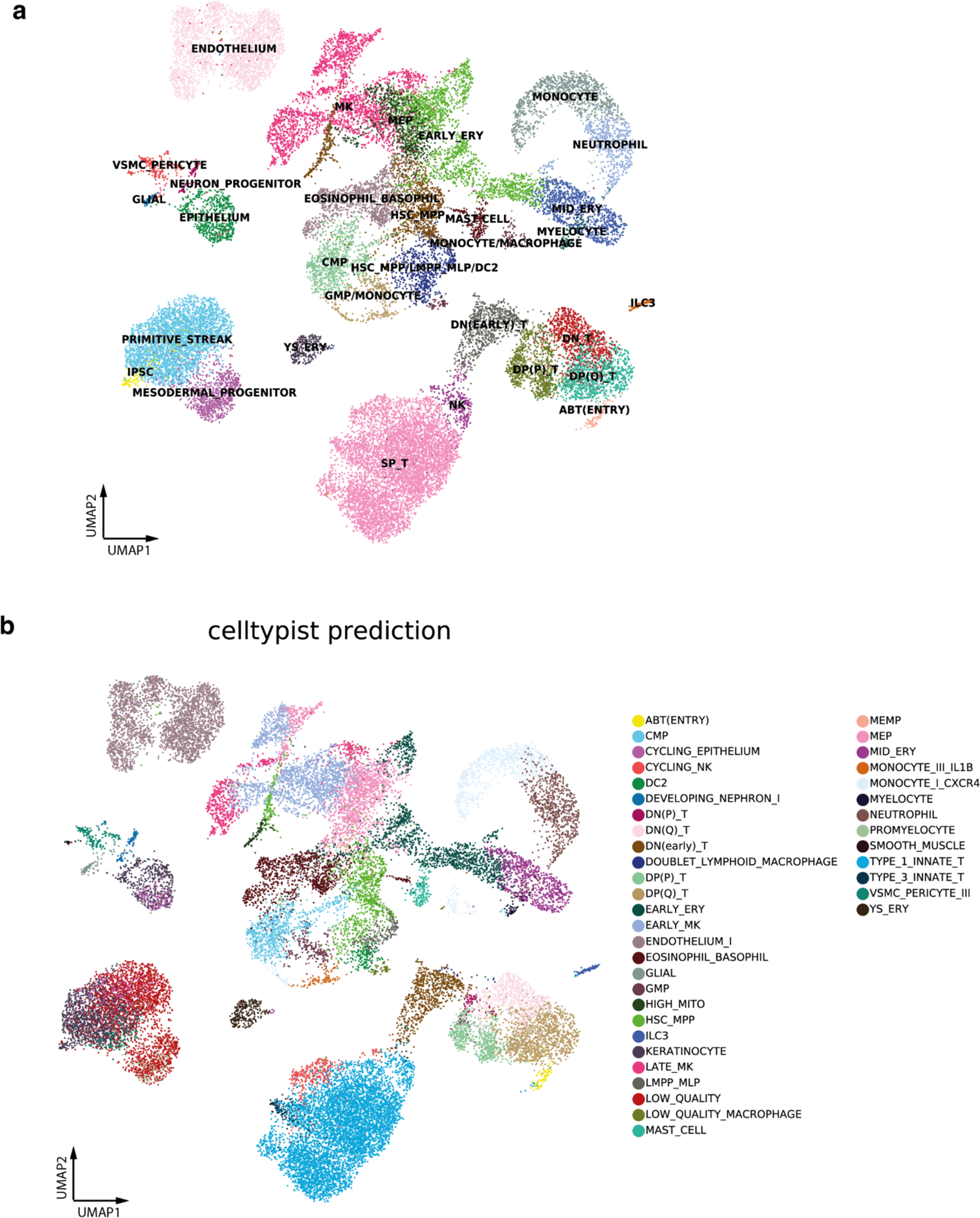
Analysis of artificial thymic organoid scRNA-seq data. **a**, UMAP visualization of different cell types in the ATO (refined annotation). iPSC: induced pluripotent stem cell, HSC_MPP: hematopoietic stem cell, and multipotent progenitor, LMPP_MLP: lymphoid-primed multipotent progenitor and multi lymphoid progenitor, DC: dendritic cell, CMP: common myeloid progenitor, GMP: granulocyte and monocyte progenitor, MK: megakaryocyte, MEP: megakaryocyte erythroid progenitor, YS_ERY: yolk sac-like erythrocyte, EARLY_ERY: early erythrocyte, MID_ERY: mid-stage erythrocyte, DN(EARLY) T: early double negative T cell, DN T: double negative T cell, DP(P) T: proliferating double positive T cell, DP(Q) T: quiescent double positive T cell, SP T: single positive T cell, NK: natural killer cell, ILC: innate lymphoid cell. **b**, Predicted annotations from logistic regression model with CellTypist^44^ using the developing human immune atlas^42^ as the training dataset, overlaid on the same UMAP plot as in **a**.

**Extended Data Fig. 8.**
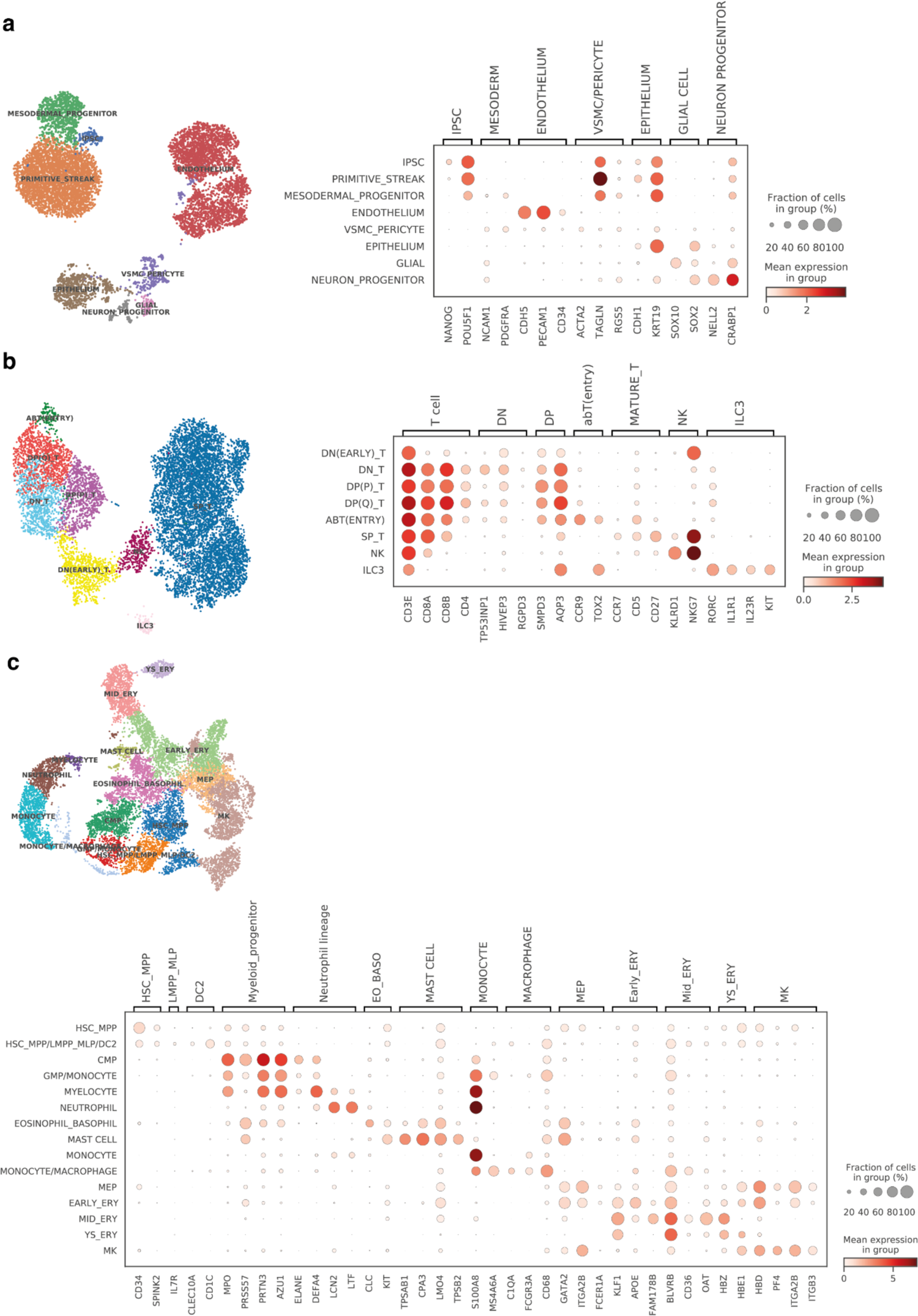
Annotation of artificial thymic organoid scRNA-seq data. For each subset lineage embedding generated through scVI, we show UMAP embeddings of cells colored by annotated cell populations and dot plots of mean expression (log-normalized counts, dot color) and fraction of expressing cells (dot size) of marker genes (columns) used for cell population annotation (rows). **a**, Annotation of non-hematopoietic cells. **b**, Annotation of T/ILC/NK lineage cells. **c**, Annotation of other hematopoietic cells that are not in **b**.

**Extended Data Fig. 9.**
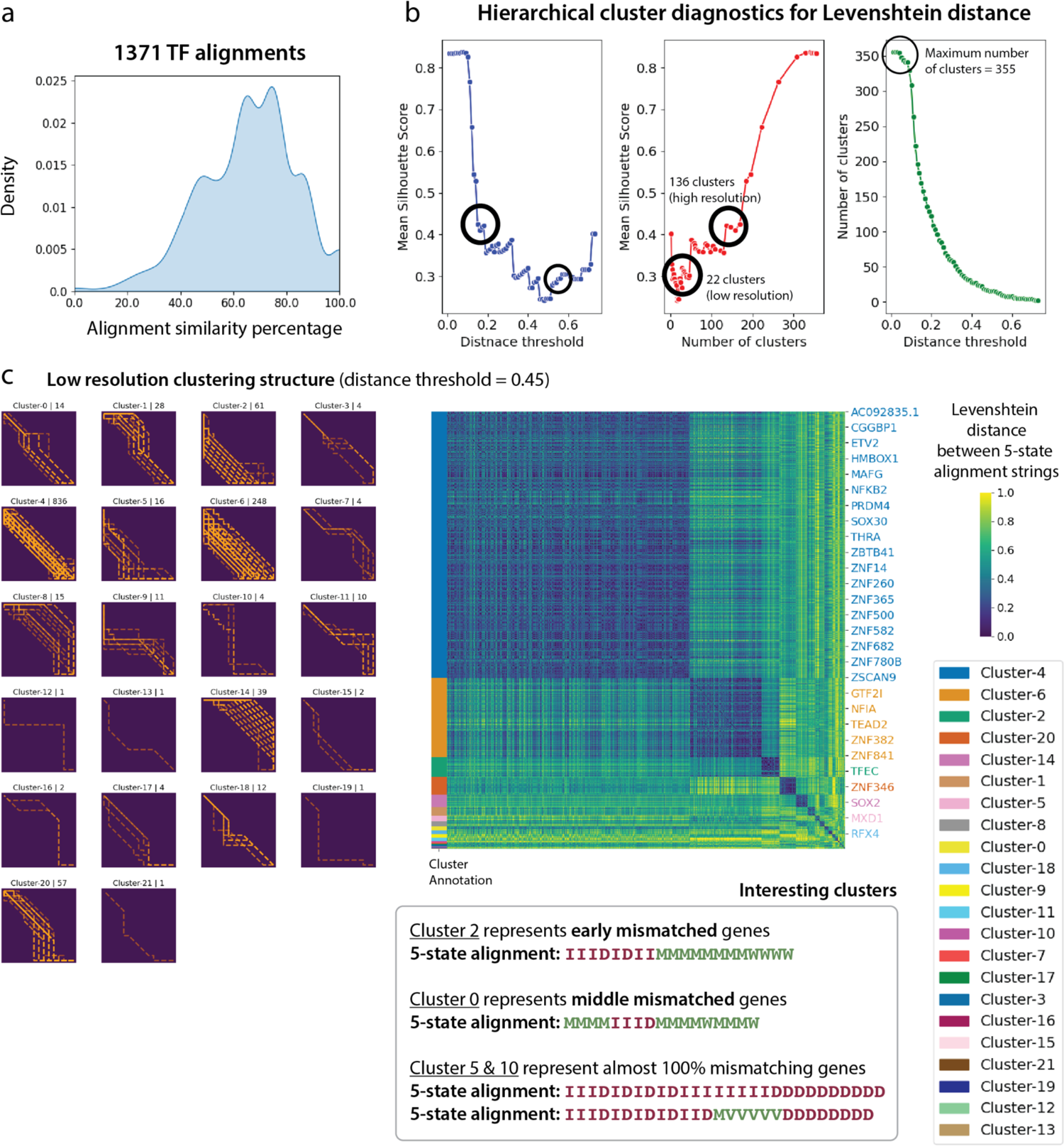
Pan fetal reference vs artificial thymic organoid alignment clustering outputs. **A**, Density plot of the alignment similarity distribution (i.e. distribution of the percentage of matches/warps across all the alignment outputs) for all 1371 transcription factors. **b**, Cluster diagnostic plots for the hierarchical agglomerative clustering of those 1371 TF alignments in terms of the mean Silhouette score when varying the Levenshtein distance threshold (or the number of clusters). The highest number of clusters represent the number of all unique 5-state alignment strings (i.e. 355 strings). Bold highlighted circles mark the local optimal mean Silhouette scores which give 22 optimal clusters for the genes at 0.45 distance threshold (low resolution), and 136 clusters at 0.18 distance threshold (high resolution). **c**, The identified clustering structure. Left: Each plot titled by “Cluster-x | n” is the pairwise matrix of reference and query time points, visualizing alignment paths for all the genes (one alignment per gene and a total of n genes in the cluster) in a cluster x. Right: The clustermap of the pairwise Levenshtein distance matrix of all TF alignments. Bottom: Identified interesting clusters (i.e. Cluster 2 representing early mismatched TFs, Cluster 0 representing middle mismatched TFs, Cluster 5 & 10 representing almost 100% mismatched TFs), with their aggregate alignments as 5-state strings.

**Extended Data Fig. 10.**
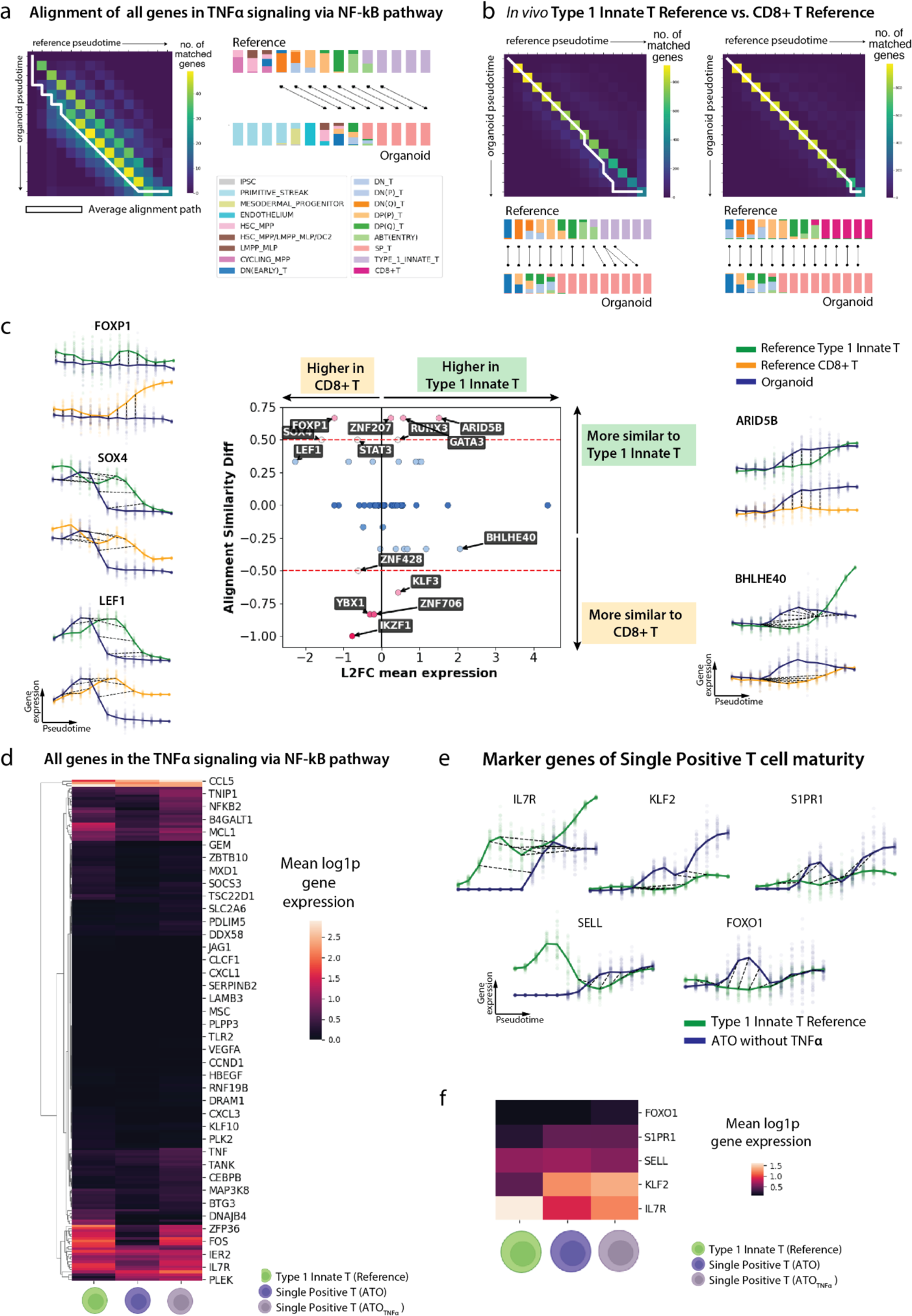
Further downstream analysis of pan fetal reference vs artificial thymic organoid alignment. a, Aggregate alignment result for all 196 genes in the TNFα signaling via NF-ϰB pathway, between *in vitro* organoid (i.e. ATO) and *in vivo* reference. Left: the pairwise time point matrix between organoid and reference pseudotime. The color represents the number of genes showing match or warp for the given pair of an organoid time point and a reference time point. The white line represents the main average alignment path. Right: Schematic illustration of the mapping between pseudotime points described by the aggregate alignment. The stacked bar plots represent the cell compositions at each time point (14 equispaced time points on pseudotime [0,1]), colored by the cell types, for reference and organoid separately. **b**, The aggregate alignment plots similar to **a**, reported for the alignment between Type 1 Innate T cell reference and ATO across 1220 TFs (left), and CD8+ Reference and ATO across 1219 TFs. **c**, Plots showing the alignment differences between *in vivo* conventional CD8+T lineage *versus in vitro* organoid, and *in vivo* type 1 innate T cell lineage *versus in vitro* organoid. Middle: plot of alignment similarity difference (*y*-axis) against log_2_ fold change of mean expression between CD8+T and type 1 innate T cells (*x*-axis). The color reflects the absolute value of alignment similarity difference. Surrounding plots: the interpolated log1p normalized expression (*y*-axis) against pseudotime (*x*-axis) showing the alignment between *in vivo* type 1 innate T cell lineage (green) and *in vitro* organoid (blue) (top), and the alignment between *in vivo* CD8+T lineage (orange) and *in vitro* organoid (blue) (bottom), for four selected genes. **d**, Heatmap of mean log1p normalized gene expression of all genes (196 genes) within TNFɑ signaling pathway in reference (*in vivo* type 1 innate T cells), ATO SP T cells without TNFɑ (i.e. wild type), and ATO SP T cells with TNFɑ treatment. **e**, Gene-level alignments for five markers of single positive T cell maturity (*IL7R, KLF2, S1PR1, SELL, FOXO1*) between the type 1 innate T reference and ATO. Each gives the interpolated log1p normalized expression (*y*-axis) of a gene against pseudotime (*x*-axis). The bold lines represent mean expression trends, while the faded data points are 50 random samples from the estimated expression distribution at each time point. The black dashed lines represent matches and warps between time points. **f**, Mean log1p normalized gene expression of the aforementioned marker genes compared across the reference type 1 innate T cells, SP T cells from wild-type ATO, and SP T cells from TNFα-treated ATO.

## Supplementary Figures

**Supplementary Fig. 1.**
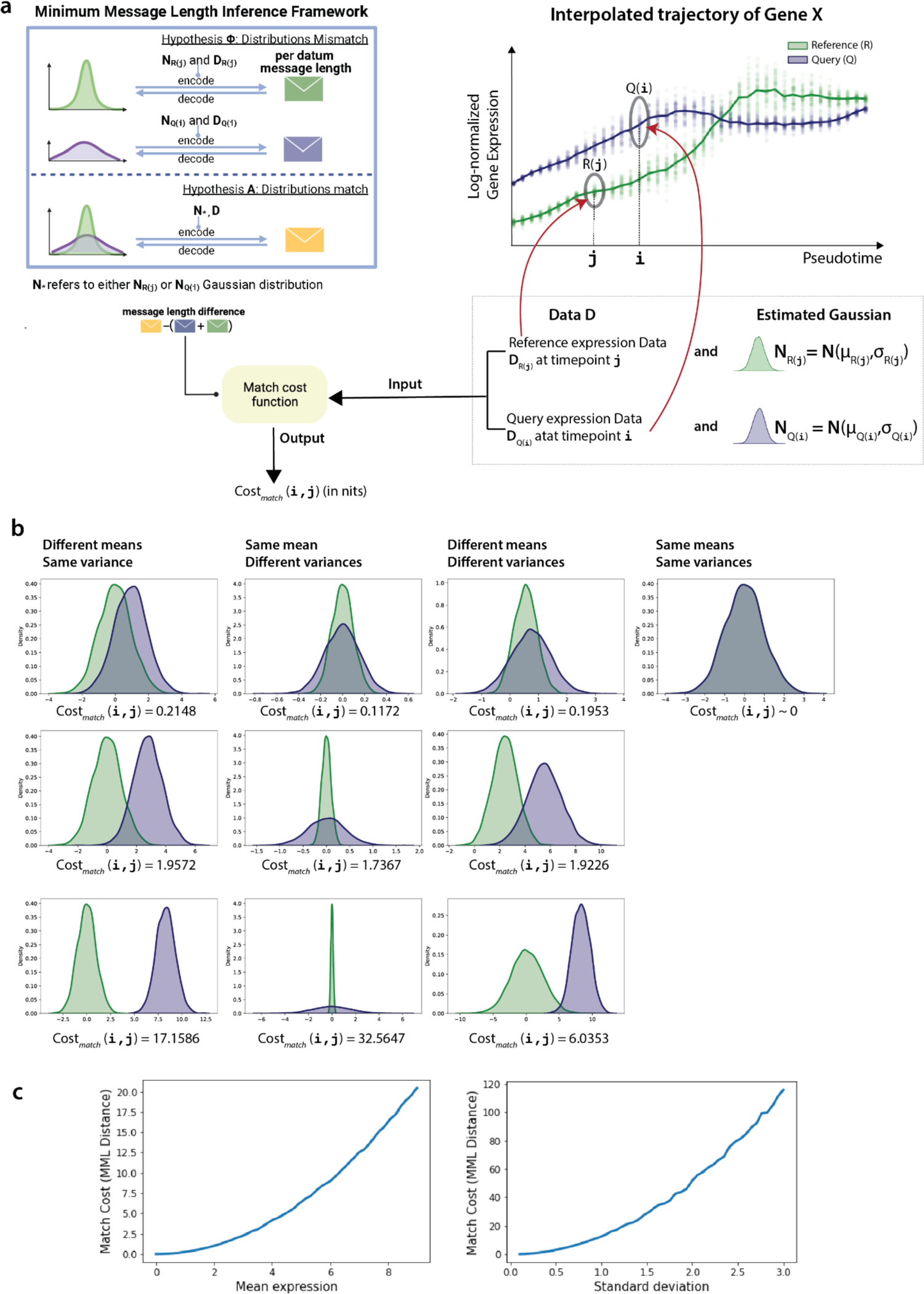
Minimum message length (MML) inference based distance function to compute the cost. *Cost*_*matc*<_(*i, j*) **of matching a reference time point** *j* **and query time point** *i***. a**, The top right plot gives the interpolated log-normalized expression (y-axis) of a particular gene X in the observed single-cell data of a reference *R* and query *Q*, against their pseudotime estimates (x-axis). The bold lines represent mean expression trends. The faded data points represent the interpolated data (i.e. 50 random samples from the estimated Gaussian distribution at each time point). As detailed in the Methods of the main text, the scoring scheme of the G2G DP algorithm computes the cost of matching every pair of time points between *R* and *Q* based on their corresponding interpolated expression distributions. Here we consider an example reference time point *j* and query time point *i*, and their respective single-cell expression datasets, *R*(*j*) and *Q*(*i*), as circled in the plot. Their interpolated expression distributions are *N*(μ_*R*(j)_, σ_*R*(j)_) and *N*(μ_&(i)_, σ_&(i)_), denoted by *N*_*R*(j)_ and *N*_&(i)_, respectively. Their interpolated expression data vectors are *D*_*R*(j)_ and *D*_&(i)_. The top left is the schematic illustration of our MML framework, extending the overview schematic **Fig. 2** (top left) in the main text. Under the MML framework, we define two hypotheses: Hypothesis **A:** the (*i, j*) time points match, and Hypothesis Φ: the (*i, j*) time points mismatch. Next we compute: (1) the total (per datum) message length of encoding **A** and *D* jointly, and (2) the total (per-datum) message length of encoding Φ and *D* jointly. Then we define *Cost*_*matc*<_(*i, j*) to be the difference between those two message lengths, measured in nits (a unit of information). **b**, Example cases of distributional differences (caused by the difference in means, the difference in variance, the difference in both mean and variance) between *R*(*j*) and *Q*(*i*), and their *Cost*_*matc*h_(*i, j*) values measured in nits. When the mean and variance is the same, *Cost*_*matc*h_(*i, j*)∼0. **c**, Behavior of *Cost*_*matc*h_(*i, j*) as the difference between the distributions increases. Left plot: *Cost*_*matc*h_(*i, j*) between the standard Gaussian distribution *N*(0,1) and *N*(μ, 1) Gaussian distribution for μ ∈ [0,9] at 50 equispaced points. 5000 data points have been randomly sampled from each *N*(μ, 1) distribution to represent itself. Right plot: *Cost*_*matc*h_(*i, j*) between the standard Gaussian distribution *N*(0,1) and *N*(0, σ) Gaussian distribution for σ ∈ [0.1,3] at 50 equispaced points. 5000 data points have been randomly sampled from each *N*(0, σ) distribution to represent itself.

**Supplementary Fig. 2.**
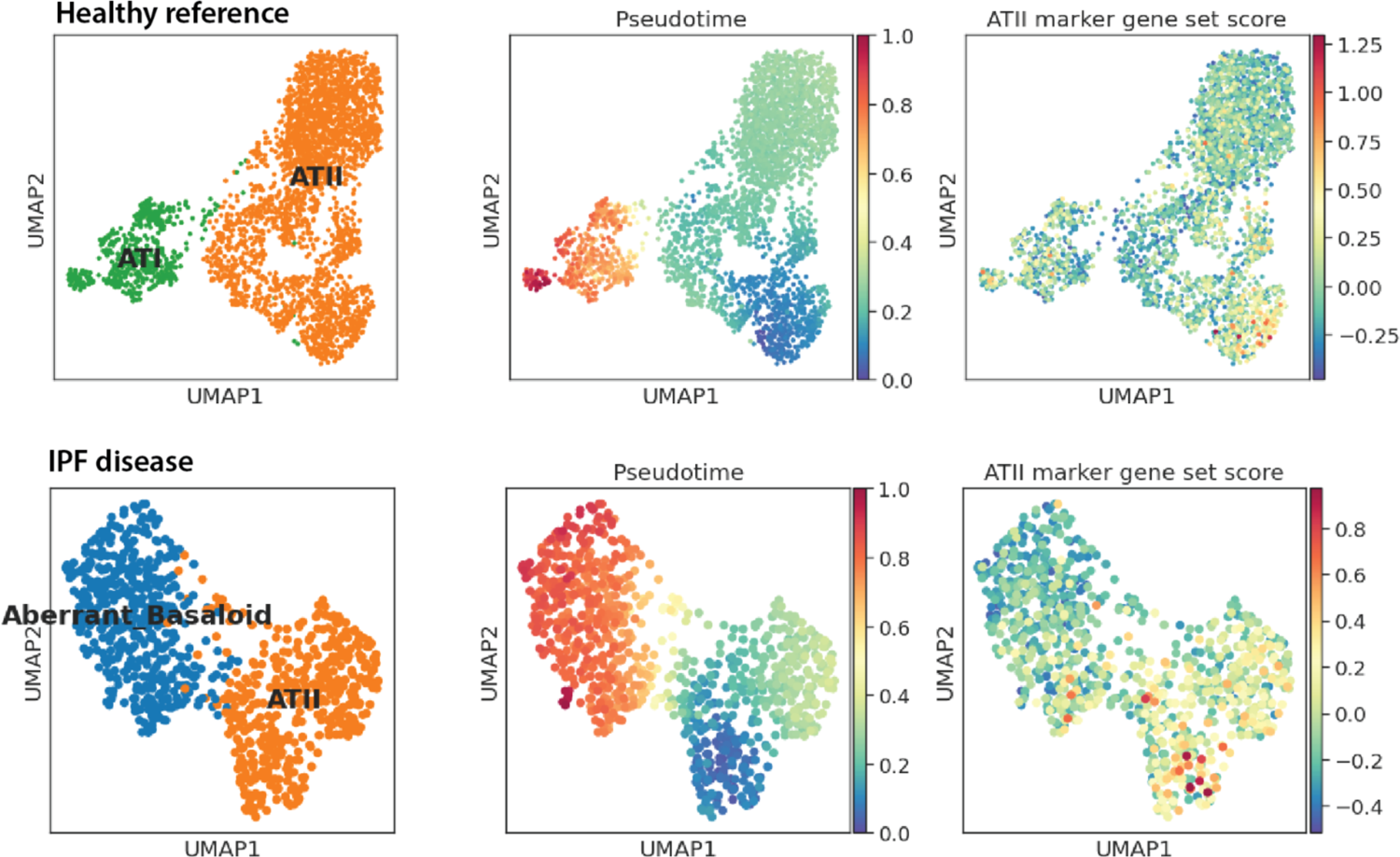
Healthy versus Idiopathic Pulmonary Fibrosis (IPF) disease case study. The differentiation trajectories of alveolar type 2 (AT2) cells into alveolar type 1 (AT1) cells in the healthy lung aberrant basaloid cells (ABC) in the IPF lung, their estimated Diffusion Pseudotime, and the gene set score of the AT2 progenitor marker genes (*AXIN2, FGFR2, ID2, FZD6, LRP5, LRP6*) on the UMAP projection of (top) healthy and (bottom) IPF cells in the Adams et al. (2020)^27^ dataset.

**Supplementary Fig. 3.**
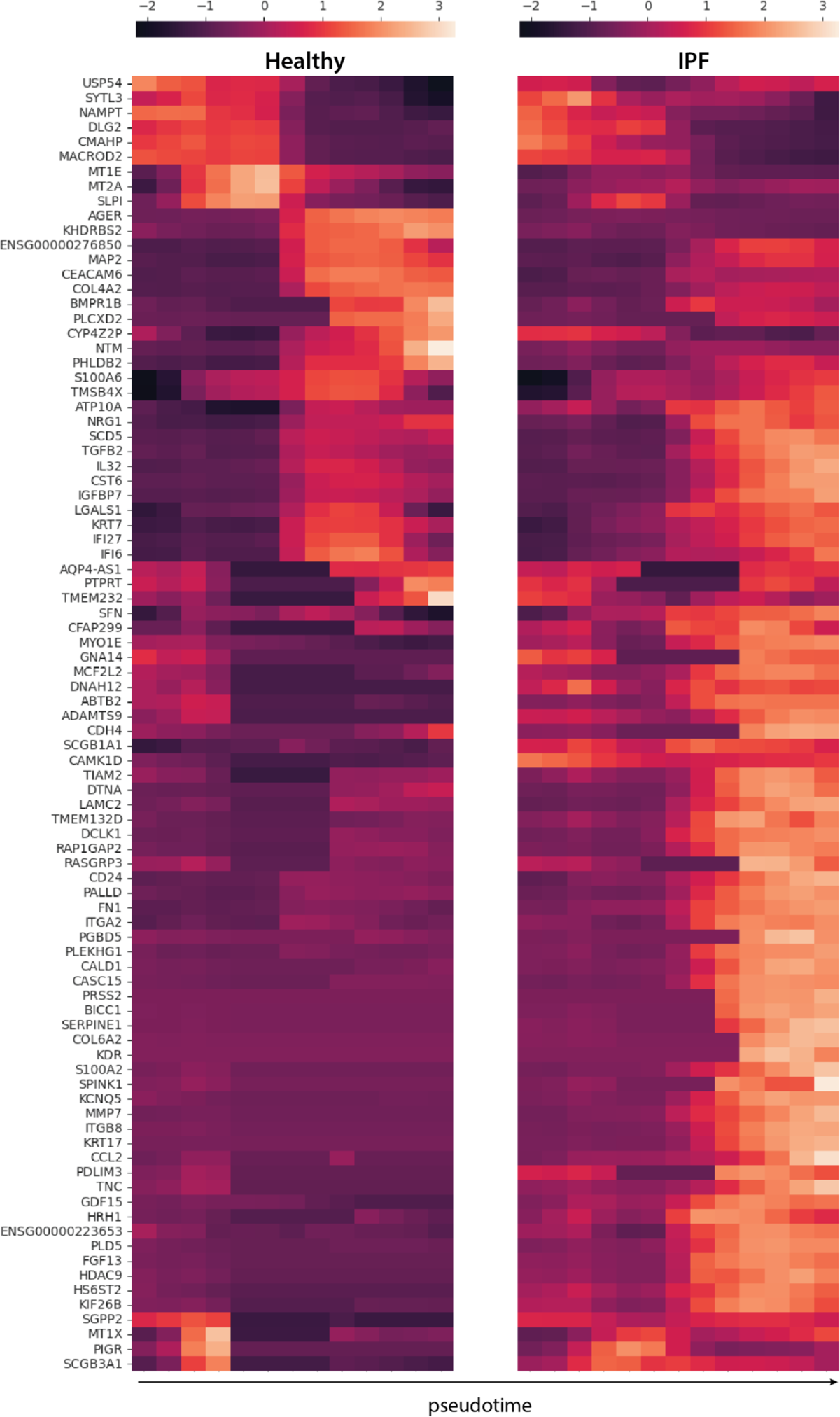
Healthy vs IPF case study: Heatmap of the smoothened (interpolated) and z-normalized mean log1p gene expression of 88 marker genes of aberrant basaloid cells (ABC).

**Supplementary Fig. 4.**
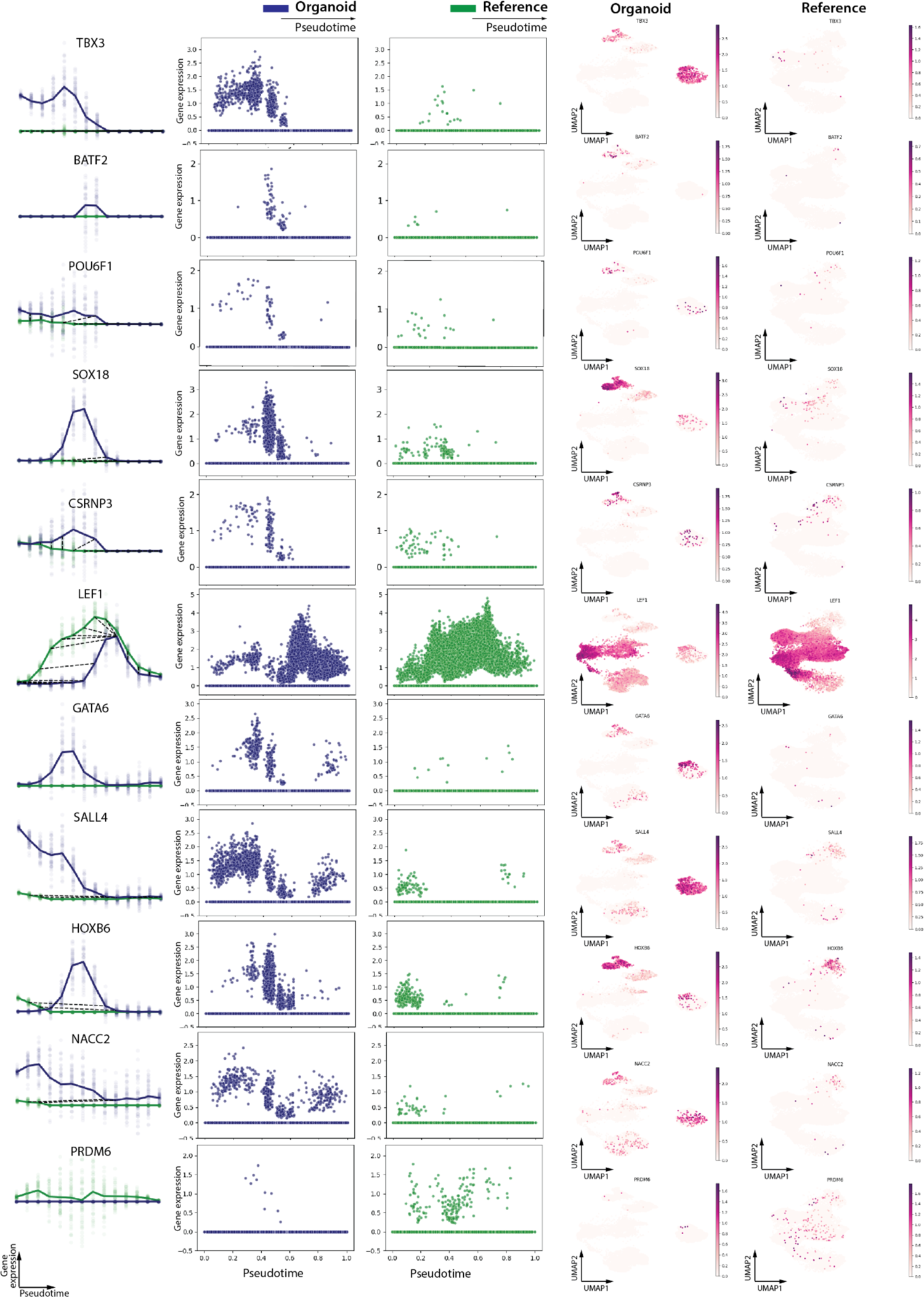
Gene-level alignments of example genes in the Pan fetal reference vs. artificial thymic organoid alignment. These are identified under the clusters with interesting alignment patterns as illustrated in **Extended Data Fig**. **9c**. Each row panel presents data for a single gene. Left plot: the interpolated log1p normalized expression (*y*-axis) against pseudotime (*x*-axis). The bold lines represent mean expression trends, while the faded data points are 50 random samples from the estimated expression distribution at each time point. The black dashed lines represent matches and warps between time points. Middle two plots: the actual log1p normalized expression (*y*-axis) against pseudotime (*x*-axis). Each point represents a cell. Right two plots: The same UMAP visualization as in **Extended Data Fig. 6**, subsetted to *in vitro* cells from ATO in the left plot, and *in vivo* cells from the pan fetal reference, colored by the corresponding gene expression value.

**Supplementary Fig. 5.**
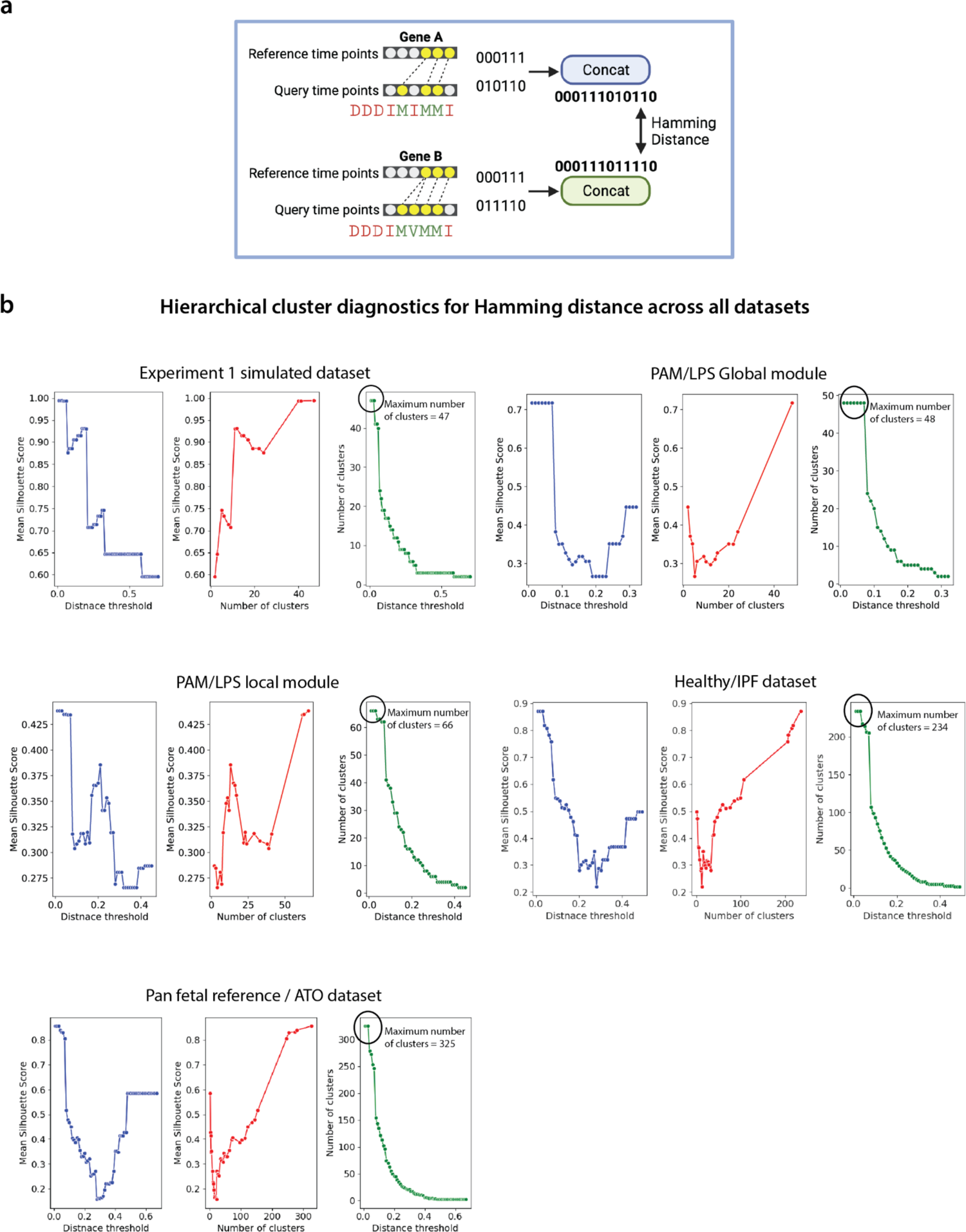
Cluster diagnostic plots for Hamming distance. a, Schematic illustration of how the Hamming distance is computed between two 5-state alignment strings. Both 5-state strings are first binary-encoded independently to obtain two equal-length binary strings (as described in Methods), which are then used to compute the Hamming distance between them. b, Cluster diagnostic plots for the Hamming distance based hierarchical agglomerative clustering of gene alignments across all the relevant datasets explored in the manuscript. These plots report the mean Silhouette score for the clustering structure when varying the Hamming distance threshold (or the number of clusters). Unlike when using Levenshtein distance, the highest number of clusters in these cases does not necessarily represent the number of all unique 5-state alignment strings.

**Supplementary Fig. 6.**
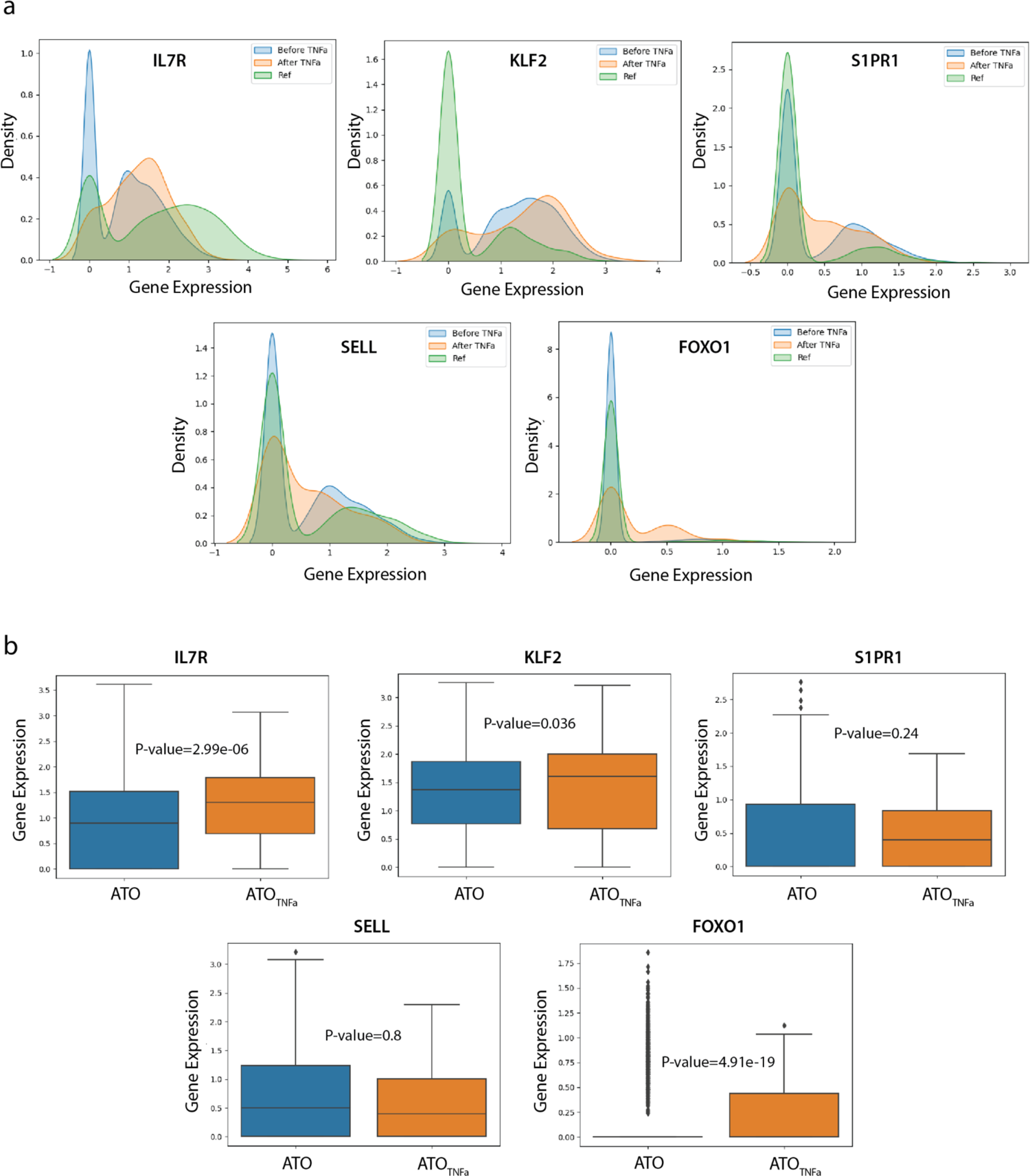
Expression distributions of the marker genes of Single Positive T cell maturation. **a**, Density plots of the gene expression distributions of *IL7R, KLF2, S1PR1, SELL*, and *FOXO1* before and after TNFα treatment of the ATO compared to the pan fetal reference. **b**, Boxplot visualization of the gene expression distributions of the same marker genes, compared between the wild-type ATO and the TNFα treated ATO, with P-values under the Man-Whitney U test reporting the significance of the increase in expression level after the treatment.

## Supplementary Tables

**Supplementary Table 1.** List of features included in trajectory alignment frameworks. A table outlining and comparing the features of CellAlign^6^, TrAGEDy^16^ and G2G.

| Framework component | CellAlign<sup>6</sup> | TrAGEDy<sup>16</sup> | Genes2Genes |
| --- | --- | --- | --- |
| Input | (1) Log-normalized single-cell gene expression data<br>(2) Pseudotime estimates of the cells inferred using any available method of choice |  |  |
| Algorithm | Uses Dynamic Time Warping (DTW) algorithm. | Builds on top of CellAlign<sup>6</sup>, and performs post-hoc changes to the DTW alignment to capture mismatches. | Combines DTW and Gotoh's biological sequence alignment<sup>15</sup> through a new dynamic programming algorithm. |
| Alignment states | Handles matches and warps only, subjected to a weight scheme with constant weight for warp open/extension | Identifies optimal start and end time points of the trajectories for DTW alignment to exclude regions of mismatch at the beginning and end. It further filters the DTW aligned regions based on alignment cost thresholding to identify mismatches. | Handles matches, warps, and mismatches jointly, subjected to a five-state machine with state transition probabilities, handling gap/warp open/extension. |
| Trajectory Interpolation | Interpolates data using a Gaussian kernel-based weighted mean expression. | Extends CellAlign<sup>6</sup> interpolation to use a cell density weighted window size. | Extends CellAlign<sup>6</sup> interpolation to distributional interpolation using weighted variance. |
| Distance measure between a pair of reference and query time points | Uses min-max normalized, mean gene expression based Euclidean distance measure, to identify similar trends of expression dynamics across two conditions. | Uses Spearman correlation, which does not require gene expression scaling as done in CellAlign<sup>6</sup>, but does not support gene-level alignment. | Uses a minimum message length inference based distributional distance measure, aiming to compare the gene expression distributions between two conditions. |
| Alignment output | Outputs only a single, high-dimensional alignment (across a given gene list). A single, gene-level alignment can be obtained by giving a single gene input. | Modified, high-dimensional DTW alignment after pruning the matches. A single, gene-level alignment can be obtained by giving a single gene input. | Outputs gene-specific alignments with an explicit alignment state description via a five-state alignment string for all the given genes. Can output an aggregate alignment for any given set of genes. |
| Alignment | Can cluster genes only | Does not explicitly | Can cluster genes based |
| clustering | based on the pseudotime shifts in their alignments. | discuss clustering. | on their five-state alignment strings, covering both matches and mismatches. |
| Differential expression capture | Requires additional downstream tasks to extract differential genes and regions (e.g. local DTW alignment with user-defined similarity threshold). | Sliding window soft clustering approach to extract DE using t-test/Mann-Whitney U test. | A gene-specific alignment output itself is a direct description of the differential expression status along the time axis. Provides a ranked list of genes based on their alignment similarity percentage. |

**Supplementary Table 2** Grid search results for optimal 5-state machine parameters

**Supplementary Table 3** Overrepresentation analysis results of healthy vs. IPF (under <40% alignment similarity threshold)

**Supplementary Table 4** Overrepresentation analysis results of T1 vs. ATO (under <40% alignment similarity threshold)

**Supplementary Table 5** Overrepresentation analysis results of T1 vs. ATO pluripotent genes cluster

**Supplementary Table 6** Overrepresentation analysis results of T1 vs. ATO DN onwards (under <40% alignment similarity threshold)

**Supplementary Table 7** Overrepresentation analysis results of CD8+T vs. ATO DN onwards (under <40% alignment similarity threshold)

**Supplementary Table 8** ATO metadata *ATO_manifest.csv*

**Supplementary Table 9** ATO CITE-seq metadata *ATO_hashtagging.csv*

## Supplementary Data (zip file)

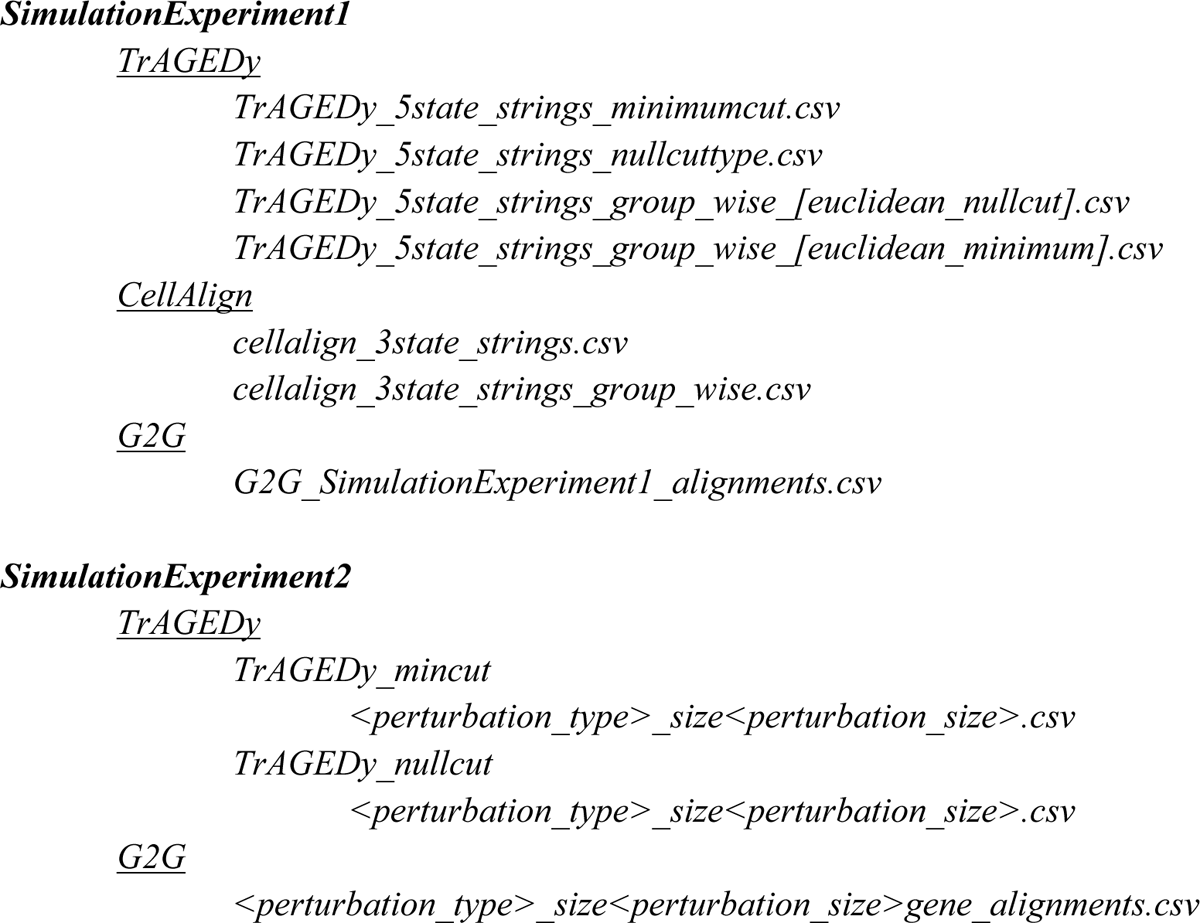

## Supplementary Note

### Additional analysis for *in vivo, in vitro* T cell comparison

We performed G2G alignment using *in vivo* conventional CD8+T cells and the relevant T lineage precursors (DN T cells onwards), with the results shown in Extended Data Fig. 10b. The most significantly enriched gene set among the mismatched genes for both cases remains the same (i.e. TNFɑ signaling via NF-ϰB pathway). To further explore the differences in the two alignment results, we focused on genes that showed the most dissimilar alignment results (genes that had alignment similarity differences > 0.5 or < −0.5 at the last stages of the trajectories) (Extended Data Fig. 10c). For this, we looked at the difference of alignment similarity percentage in the last six alignment states of the 5-state gene-level alignment strings, and the log_2_ fold change of the mean interpolated gene expression in the last time bins as heuristics to understand their differences.

Three of the genes, *SOX4, FOXP1* and *ARID5B* had large log_2_ fold change differences (absolute log_2_ fold change > 1) between type 1 innate T cells and CD8+T cells. For these three genes, the expression dynamics of *in vitro* T cell development are more similar to those of *in vivo* type 1 innate T cells, whereas *in vivo* CD8+T cells had higher *SOX4, FOXP1* and lower *ARID5B* expression in the last stages of development (Extended Data Fig. 10c). While the role of *SOX4* in CD8+T cell development is unclear, *FOXP1* has been shown to maintain a quiescent profile in naive CD8+T cells^88,89^, and our results are in keeping with a more activated profile in type 1 innate T cells. *ARID5B* has been reported to regulate metabolic programming and promote IFNγ production in NK cells^90^. The higher expression in type 1 innate T cells might explain some of their NK-like features^41,42^. We also note *LEF1* which shows a moderate alignment similarity difference and high log_2_ fold change; a gene that is known to be highly expressed in CD8+T cells compared to the type 1 innate T cells. On the other hand, for *BHLHE40*, which is downstream of TNFɑ signaling and other pro-inflammatory cytokines^91^, its expression dynamic in *in vitro* T cell development is more similar to that in CD8+T cells, while the *in vivo* type 1 innate T cells have increased expression at the end.

### TNFα validation experiment results

This section presents results for all statistical tests that check the MML distances in gene expression distributions of the SP T cells in the ATO and the type 1 innate T cells in the pan fetal reference, before and after TNFα treatment. Across all TFs and different relevant pathway gene sets (i.e. TNFα pathway, P38, JNK, NF-ϰB canonical signaling known to be targeted by TNFα signaling), we tested the mean change in MML distances of gene expression across all significantly distant genes, as well as across all genes.

The significance of distance in gene expression between reference and query was computed under the empirical CDF null model. When considering all genes, the mean change statistic may be misleading due to the noise in distances. Checking between the significant distances allows us to avoid such bias. Overall, we see that on average, the significantly changed genes have decreased in their gene expression distance.

| Gene set | Significance level of differential expression | # of significantly distant genes<br>diffset<sub>UNTREATED</sub> | # of significantly distant genes<br>diffset<sub>TREATED</sub> | Mean distance of diffset<sub>UNTREATED</sub><br>(nits) | Mean distance of diffset<sub>TREATED</sub><br>(nits) | P-value for change in distance<br>(Mann-Whitney U test) | Mean distance change of all genes<br>(nits) |
| --- | --- | --- | --- | --- | --- | --- | --- |
| All TFs (1371) | 0.005 | 12 | 35 | 31.51 | 12.91 | 0.0006131<br>978745461<br>366 | 0.404<br>→ 0.6<br>(-) |
| All TFs (1371) | 0.05 | 24 | 91 | 18.54 | 6.67 | 2.9617483<br>396576014<br>e-05 |  |
| TNFα signaling via NF-κB - TFs (42) | 0.005 | 11 | 13 | 28.54 | 9.98 | 0.0119253<br>644275241<br>57 | 7.77<br>→ 3.36<br>(+) |
| TNFα signaling via NF-κB - TFs (42) | 0.05 | 15 | 19 | 21.38 | 7.26 | 0.0446091<br>620161070<br>2 | |
| TNFα signaling via NF-κB - all genes (196) | 0.005 | 32 | 30 | 21.18 | 10.41 | 0.0084802<br>992372226<br>36 | 3.771<br>→ 1.899<br>(+) |
| TNFα signaling via NF-κB - all genes (196) | 0.05 | 57 | 54 | 12.6 | 6.5 | 0.1558075<br>120839859 | |
| All TFs excluding TNFα TFs (1329) | 0.005 | 3 | 27 | 24.84 | 12.47 | 0.0362068<br>965517241<br>34 | 0.171<br>→ 0.512<br>(-) |
| All TFs excluding TNFα TFs (1329) | 0.05 | 12 | 70 | 10.65 | 6.6 | 0.0144675<br>412638397<br>72 | |
| NF-κB canonical pathway (215) | 0.005 | 8 | 29 | 24.45 | 10.76 | 0.0003306<br>048846845<br>811 | 1.199<br>→ 1.858<br>(-) |
| NF-κB canonical pathway (215) | 0.05 | 18 | 77 | 12.48 | 5.01 | 0.0007696<br>929346291<br>883 | |
| P38 (50) | 0.005 | 4 | 6 | 40.5 | 12.23 | 0.0190476<br>190476190<br>5 | 3.361<br>→ 1.795<br>(+) |
| P38 (50) | 0.05 | 4 | 12 | 40.5 | 6.94 | 0.0021978<br>021978021<br>98 |  |
| Jnk(155) | 0.005 | 2 | 15 | 16.03 | 8.85 | 0.0882352<br>941176470<br>6 | 0.525<br>→ 1.307<br>(-) |
| Jnk (155) | 0.05 | 13 | 51 | 5.04 | 3.77 | 0.0296726<br>662308187<br>1 |  |

We note that for many cases, the distance has significantly dropped despite the increase in the number of genes with significant distances after TNFα treatment. This is because the highly distant genes before treatment have now got closer, outweighing the increase in the number of differential genes.

For TFs, there are 20 TFs that are significantly distant both before and after treatment, implying 4 TFs (*FOSB, GPBP1, KLF6, REL*) have rectified its distance after treatment. Among the 20 common TFs, 14 TFs (70%) are on the path to getting closer to the reference (i.e. *CEBPD, CREM, CSRNP1, FOS, JUN, JUNB, NME2, NR4A1, NR4A2, PA2G4, TSC22D1, YBX1, ZNF331, ZNF683*). There are 71 TFs that got significantly different after treatment, out of which, 11 TFs (i.e. *BHLHE40, HIF1A, JUND, KDM2A, NCOA3, NFKB2, RELB, RUNX3, STAT4, USF2, ZBTB1*) surpassed the reference expression in the right direction (high expression or low expression), suggesting the need for some modulation.

For genes in the TNFα signaling pathway, the number of significantly distant genes have dropped from 57 to 54 after TNFα treatment. There are 22 genes that have rectified its distance after treatment (*ATF3,BIRC2,CCL4,CCNL1,CD69,DUSP1,FOSB,FOSL2, ID2,IER2,KDM6B,KLF6,MAP3K8,NFE2L2,PDE4B,PTGER4,SGK1,SIK1,SLC2A3,SQSTM1, TNFAIP3,ZC3H12A*). 35 genes remained distant after treatment, out of which, 24 genes got closer to the reference than before (i.e. *BTG2, CEBPD, DNAJB4, DUSP2, DUSP4, FOS, GADD45B, GEM, GPR183, IER3, IL7R, JUN, JUNB, MXD1, NFKBIA, NR4A1, NR4A2, PHLDA1, PPP1R15A, REL, RHOB, SAT1, TSC22D1, ZFP36*). There are only 19 genes that were similar before, but got distant after treatment, out of which 5 genes (*BHLHE40, DUSP5, IER5, NFKB2, RELB*) surpassed the reference expression in the right direction.

We note that other TFs and genes could become more distant after TNFα treatment as a result of TNFα induced stress response. This may explain why the average distances across all genes in the NF-ϰB and JNK pathways have increased (as stress seems to be activating those pathways – References: https://www.ncbi.nlm.nih.gov/pmc/articles/PMC2823860/, https://www.ncbi.nlm.nih.gov/pmc/articles/PMC5046695/). However, despite such changes, the fact that CellTypist still annotates the TNFα treated ATO cells as type 1 innate T, and their increased expression of gene markers of SP T cell maturation suggest more maturity. Thus we conclude that TNFα signaling is a potential direction for the ATO protocol refinement, which is worth exploring further.

### Inspecting the known markers of SP T cell maturity

We also checked the differences in the expression distributions of known SP T cell maturation markers reported in literature. *IL7R* has shown to be initiated in mature SP thymocytes, and its expression is dependent on NF-ϰB signaling, triggered by TNFα signaling^92,93,94^.

#### Comparing the MML distances between ATO and reference before and after TNFα treatment (Supplementary Fig. 6a)

The MML distance of *IL7R* between the reference and ATO_TNFɑ_ is 13.347 nits, which is significantly different (P-value=∼0 under empirical CDF), while it significantly drops to 4.041 nits after TNFα treatment, with an increased induction of expression as expected from mature SP T cells. It remains different between reference and ATO_TNFɑ_, however with a higher p-value than before (P-value=∼0.0004 under empirical CDF). *KLF2* is more upregulated in the ATO than the reference, and has further upregulated in ATO_TNFɑ_, thus increasing the distance from 17.0983 nits to 21.9164 nits. *KLF2* upregulation is expected in mature SP T cells^95^. *S1PR1* becomes less distant between reference and ATO_TNFɑ_ (from 1.3549 nits to 1.2962 nits) even if it remains differential at 0.01 significance. *FOXO1* on the other hand is more upregulated in ATO_TNFɑ_ than before, increasing the distance to the reference. *SELL* expression is known to be maintained by *FOXO1* and *KLF2* ^96^. However, *SELL* shows no significant difference to the reference before and after TNFα treatment.

#### Complementary comparison of marker expression between ATO and ATO_TNFɑ_ (Supplementary Fig. 6b)

Complementary to the above analysis, we performed the Mann-Whitney U test to check if the ATO expression is significantly lesser than ATO_TNFɑ_ expression for the above marker genes. Accordingly, *IL7R, KLF2, FOXO1* have significantly increased expression, while *S1PR1* and *SELL* do not show significant difference before and after treatment.

## Notes

### Summary of Updates

A substantially revised manuscript which includes new benchmarking results.

